# Records of RNA localization through covalent tagging

**DOI:** 10.1101/785816

**Authors:** Hugo C. Medina-Muñoz, Christopher P. Lapointe, Douglas F. Porter, Marvin Wickens

## Abstract

RNA movements and localization pervade biology, from embryonic development to disease. To identify RNAs at specific subcellular locations, we anchored a uridine-adding enzyme at those sites, which then marked RNAs in its vicinity with 3’ terminal uridines. RNAs were tagged independent of their translation status, and included not only mRNAs, but also ncRNAs and ncRNA processing intermediates. A battery of RNAs, including the stress sensor, *IRE1*, were tagged at both ER and mitochondria, and reveal RNAs whose dual localization is conserved from yeast to human cells.

## INTRODUCTION

Localization of specific RNAs to discrete sub-cellular locations was first observed in striking examples during early development^1-6^ and in yeast^7^. We now know RNA localization is widespread, and critical in secretion, patterning, cell fate determination, and neurobiology^8,9^. Many mRNAs in embryos and mammalian cells exhibit discrete patterns of localization, emphasizing its breadth. Localization hinges on interplay between sequences in the RNAs, RNA binding proteins, molecular motors, and subcellular structures, such as the ER or cytoskeleton^10-13^. Advancements in FISH^14-17^, live imaging^18-20^ and sequencing-based methods including proximity-specific ribosome profiling^21,22^, APEX-RIP^23^ and APEX-Seq^24,25^ are very powerful, but typically provide snapshots of localized RNAs, since cells do not survive the required treatments. They also often require either custom oligonucleotide probes or sophisticated equipment. Approaches are needed to identify RNAs at any subcellular location across the entire transcriptome, and to do so in living cells.

We developed a strategy that registers RNAs as they interact with a cellular site *in vivo*. Our approach builds on RNA Tagging, used initially to identify RNAs that bind cognate proteins *in vivo*^26^. In those studies, the RBP to be tested was linked to an enzyme that adds uridine residues (termed a “PUP,” “TUTase”^27^, or TENT^28^) to the RNA. When the chimeric protein bound an RNA molecule *in vivo*, the end was “tagged” with uridine residues. The number of uridines added to each mRNA molecule mirrored the affinity of its sites for the RBP, and likely the integrated dwell time of the RBP on that particular RNA molecule^26^, as observed *in vitro*^29^.

Here we describe Localized RNA Tagging, in which RNAs at specific sites are identified, focusing on the ER and mitochondria of the yeast, *S cerevisiae.* A U-adding enzyme is attached to an anchoring protein that resides at a specific location. The anchored enzyme tags RNA molecules it encounters *in vivo*, which are identified through deep sequencing. Since cells live during tagging, the number of U’s added likely mirrors the cumulative time that an individual RNA molecule spends at that location.

## RESULTS

### Broad-specificity tagging

We first designed a protein construct intended to tag most or all cellular RNAs, and so provide a baseline for comparison. We selected *C. elegans* PUP-2^27^ as the tagging agent. This enzyme adds uridines to RNA 3’ ends, lacks RNA-binding domains, and has been used to identify RNAs bound to specific proteins in living cells^26,30,31^ (Figure 1A). To facilitate tagging of most RNAs in the cytoplasm, we linked PUP-2 to the RNA-recognition motifs (RRMs) of yeast poly(A)-binding protein, generating a construct here termed “PUP alone (+PAB).” A control chimera, “PUP alone (-PAB),” was constructed that lacked the PAB RRMs (Figure 1A). Both proteins were expressed under control of the *SEC63* promoter^21^ (used later to enable direct comparison to RNAs at the ER).

**Figure 1.**
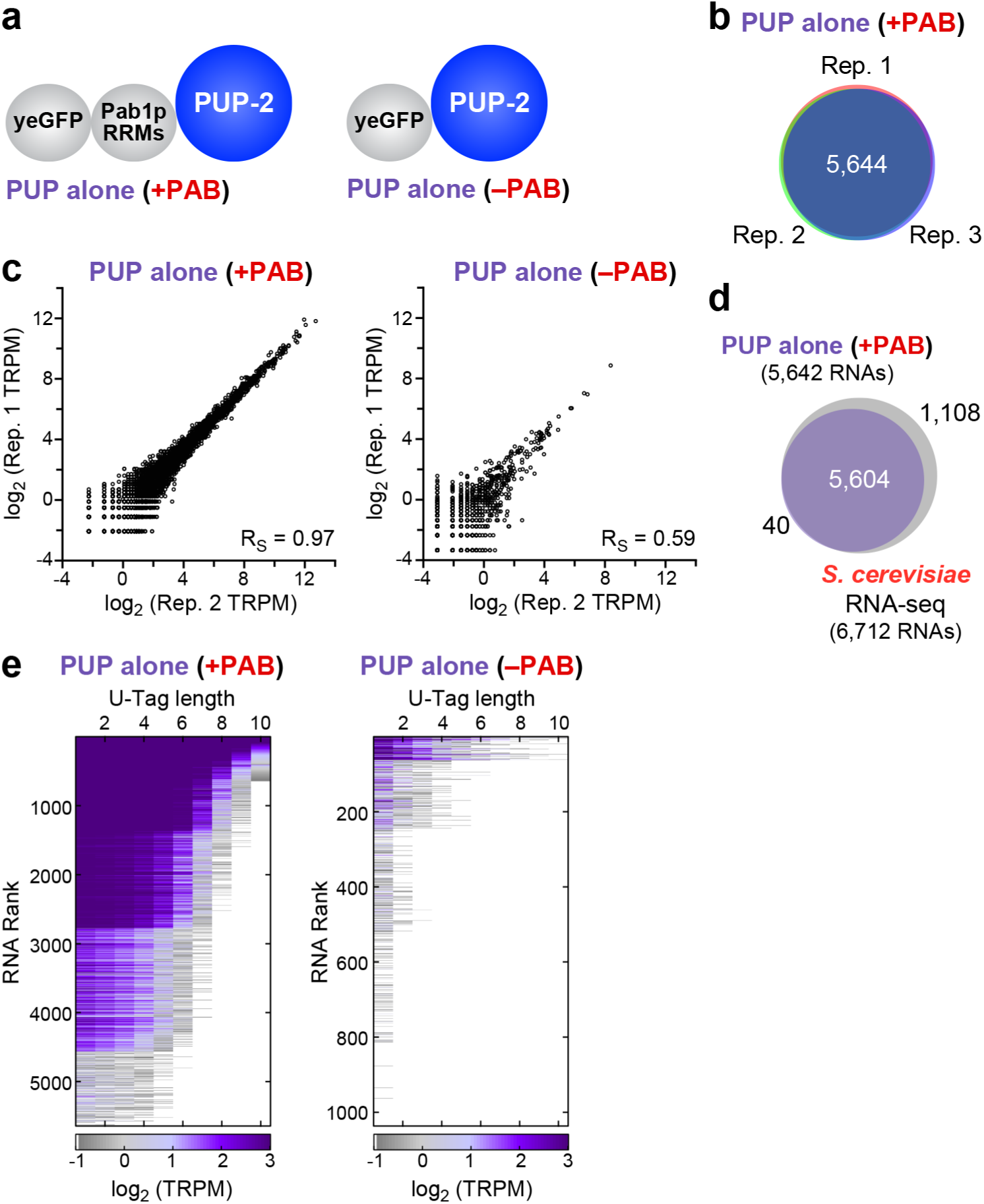
Figure 1. Broad specificity cytoplasmic RNA tagging: a baseline. **a)** PUP-2 (“PUP alone”) chimeras with or without Pab1p RRMs. **b)** Reproducibility of U-tagged RNA species with the PUP alone (+PAB) construct and of **c)** tagged reads <u>p</u>er <u>m</u>illion (TRPM) for both chimeras. Each dot represents a single RNA species. **d)** RNAs tagged by PUP (+PAB) vs all yeast RNAs. **e)** Ranked U-tagged RNAs (y-axis) with and without RRMs. Each row indicates an individual RNA species; columns represent U-tag length intervals, from 1 to 10 U’s added. Color relates to abundance of reads, as indicated in the key (bottom): purple indicates frequent reads; grey, low reads.

To identify tagged RNAs and the number of U’s they received, we prepared polyadenylated RNA via oligo(dT) selection and ribosomal RNA depletion^26^ (see *Methods*). RNAs then were reverse-transcribed using a primer designed to enrich uridylated RNAs (Supp. Figure 1A). The resulting DNA libraries were analyzed on an Illumina sequencer using paired-end sequencing (Supplementary Fig. 1A). Sequencing data were processed using a computational pipeline^26^ that identified all tagged RNAs, and for all tagged RNAs, the number of reads obtained and number of U’s added (Supplementary Fig. 1B, C).

The two “PUP alone” proteins added U’s to cellular RNAs with very different efficiencies. PUP alone (+PAB) yielded 3-4 orders of magnitude more tagged reads per million (TRPMs) across all U-tag lengths (Supplementary Fig. 2A). PUP alone (+PAB) was more reproducible (Figure 1B, C, Supplementary Fig. 2B, C), and yielded more U-tagged species (Figure 1D, Supplementary Fig. 2D). Tagging efficiency was correlated with RNA abundance with both proteins (Supplementary Fig. 2E). Tagged RNAs were ranked based on the number of U’s added and number of reads obtained (Figure 1E), which revealed the dramatic differences with and without the PAB RRMs. We adopted the protein with RRMs for subsequent experiments due to its efficiency and ability to tag most cellular RNAs. We refer to it simply as “PUP alone” hereafter.

### ER-localized RNA tagging

To detect RNAs that encounter the endoplasmic reticulum (ER), we fused the PUP alone chimera to the C-terminus of Sec63p, a protein embedded in the ER (Figure 2A), and expressed the chimeric protein from the endogenous *SEC63* locus. The Sec63p chimera, termed “ER-PUP”, is predicted to be co-translationally embedded into the ER membrane by three trans-membrane segments of Sec63p, and place the C-terminal PUP-2 domain in the adjacent cytosol^32^. As predicted, GFP fluorescence from ER-PUP mirrored the pattern reported for Sec63p, and co-localized with signal from the Sec61p-mCherry ER-marker^33,34^ (Figure 2B), indicating that ER-PUP was anchored to the ER membrane.

**Figure 2.**
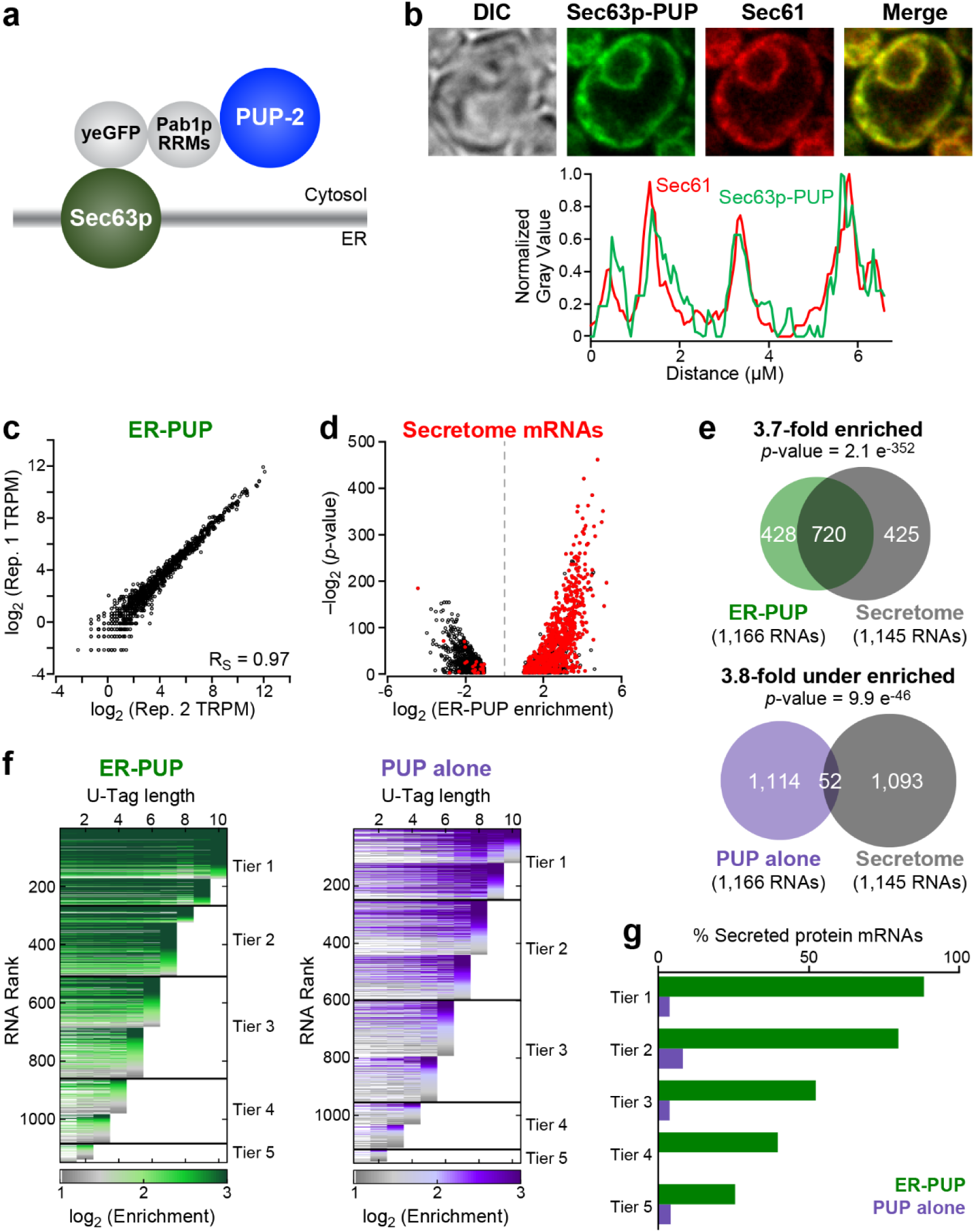
ER-localized tagging provides an *in vivo* record. **a)** ER-PUP chimera designed to tag ER-proximal RNAs. PUP(+PAB) was fused to the C-terminus of Sec63p, which tethers the chimera to the cytosolic surface of the ER.^32^ **b)** Subcellular distribution: ER-PUP GFP fluorescence (green) vs an ER marker^33,34^, Sec61p-mCherry (red); merged (yellow). Fluorescence intensities for ER-PUP (green) and the ER marker (red) across a representative cell. **c)** Reproducibility of ER-PUP tagging across biological replicates. **d)** Distribution of mRNAs that encode secreted proteins (“secretome”) among the ER-PUP-tagged mRNAs. **e)** Secretome mRNAs in tagging by ER-PUP and PUP alone. **f)** Distribution of relative enrichment (ER-PUP vs PUP alone) across each U-tag length. RNA species that had significant (adjusted *p*-value < 0.05) difference (log_2_(Δ tagged reads) ≥ 1)) in tagging efficiency (enrichment) in any of ten U-tag lengths (1U-10Us) were isolated and plotted as horizontal lines. Each row is one mRNA. Lines are segmented, and segments colored in accord with enrichment values: most enriched denoted in green (ER-PUP) or purple (PUP) and the lowest grey (both). RNAs were binned into five groups (“tiers”), and tiers ranked by the highest tag length (e.g., Tier 1 RNAs had a minimum of 9-10 U’s; Tier 5 had a minimum of 1-2U’s). **g)** Proportion (%) secretome mRNAs in each tier of ER-PUP-tagged (green) or PUP alone-tagged (purple) tier.

To identify RNAs tagged by ER-PUP, we compared RNAs tagged with and without the Sec63p anchor, using the analytical tool DESeq2^35^, a method that identifies the statistical strengths of observed differences in RNA populations (Supplementary Fig. 3A, B, C, D). The data were highly reproducible (Figure 2C, Supplementary Fig. 3E). The ER-anchored PUP selectively tagged 1,148 RNAs, which we refer to as “ER-enriched,” while the unanchored PUP preferentially tagged 1,167 (“ER-depleted”). Many mRNAs tagged by ER-PUP encoded secreted proteins^36^ (“secretome mRNAs”, *p*-value = 2.1 e^-352^) and had ER-related gene ontology (GO) associations^37,38^, which were reduced or missing among RNAs tagged less efficiently at the ER (Figure 3D, E; Supplementary Fig. 3F; and Table 1). Thus, the Sec63p anchor directed tagging to RNAs that encounter the ER *in vivo*.

**Figure 3.**
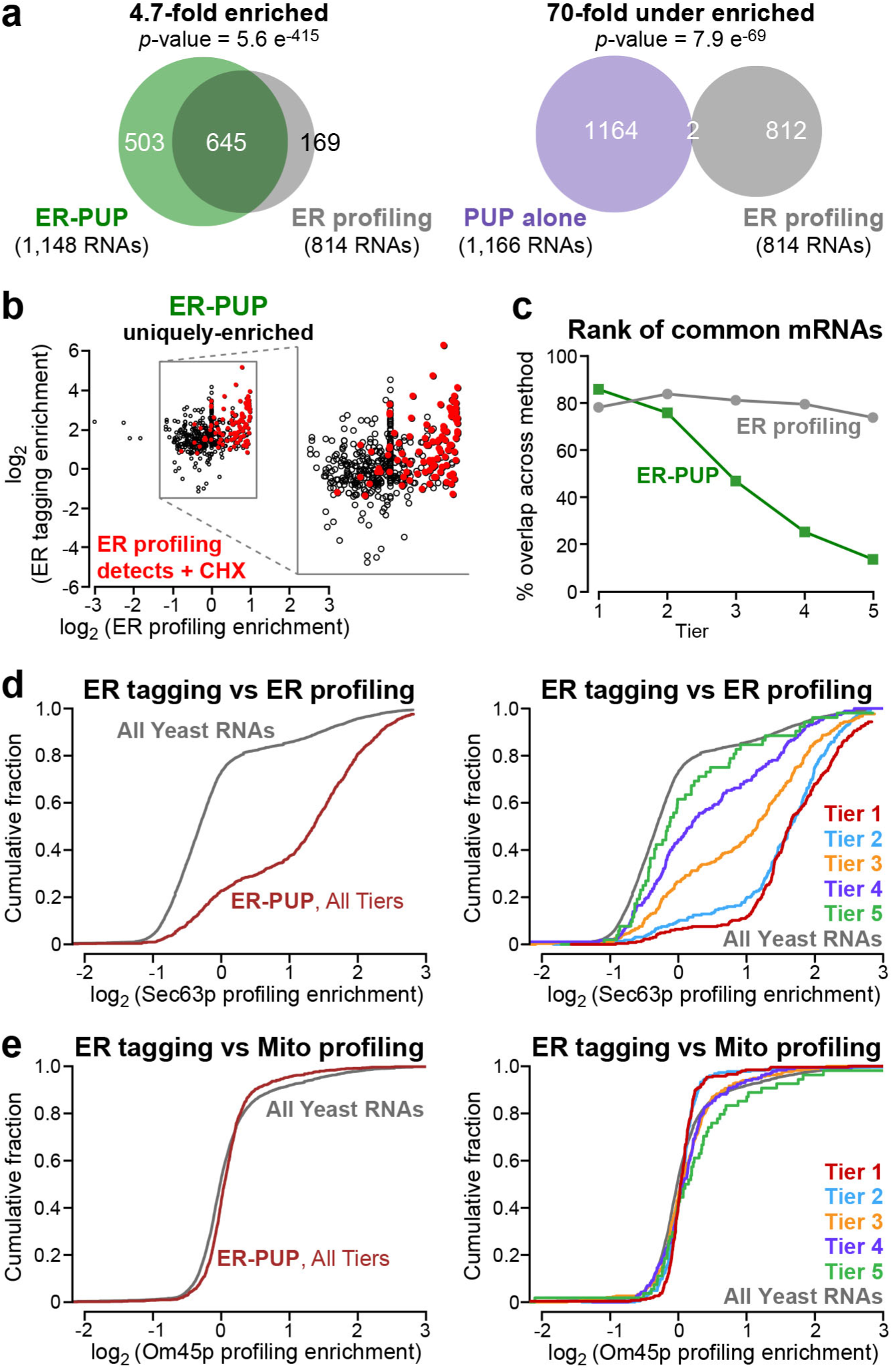
Many but not all ER-enriched RNAs are bound by ER proximal ribosomes. **a)** mRNAs enriched by ER-PUP or PUP alone vs positional profiling at the ER.^21^ **b)** Enrichment of RNAs uniquely enriched (log_2_(Δ tagged reads) ≥ 1)) by ER-PUP, and their enrichment in ER profiling. Red dots mark RNAs whose enrichment increases above the cutoff in profiling after arresting elongation with cycloheximide.^21^ **c)** Rank of the commonly enriched RNAs in tagging (green) and profiling (black). **d)** Cumulative fraction distribution of ER-enriched mRNAs vs ER profiling; per tier analysis in right panel. **e)** Cumulative fraction distribution of ER-enriched mRNAs vs mitochondrial profiling^22^, with per tier analysis on the right.

To aid in further analyses, we grouped RNAs tagged by ER-PUP and the control, PUP alone, into five tiers based on U-tag lengths. RNAs with the longest tags were grouped in Tier 1 and those with the shortest tags in Tier 5 (Figure 2F). Within a tier, each RNA was ranked by the fold-enrichments in that dataset relative to the other. With both ER-PUP and PUP alone, RNAs with the longest U-tags generally had the highest enrichment (Figure 2F). Among RNAs tagged by ER-PUP, the fraction of secretome mRNAs was highest in Tiers 1 and 2 (88% and 79%) and declined progressively to Tier 5 (25%) (Figure 2G). The control, PUP alone, yielded little enrichment or correlation with tiers (Figure 2F, G), demonstrating that ER-PUP preferentially tagged RNAs with ER-related functions.

To assess the relationship between ER-PUP enrichment and mRNA abundance, we binned all yeast RNAs into five tiers based on RNA seq^26^ (FPKM), from most (Tier 1) to least abundant (Tier 5) (Supplementary Fig. 4A, B). The distribution of abundances of RNAs tagged by ER-PUP was much more similar across tiers as compared to mRNA abundances in the cell (Supplementary Fig. 4B). Indeed, highly abundant RNAs (Abundance Tiers 1&2) with secretome association were dramatically enriched by ER-PUP (4.6-fold enriched, *p*-value = 3.98e^-233^) while ones that lack secretome association were depleted (2.9-fold depleted, *p*-value = 1.8e^-74^, Supplementary Fig. 4C). Further, the gamut of RNA abundances was represented across all ER-PUP tiers, while PUP alone tiers primarily contained the most abundant species (Supplementary Fig. 4D). Finally, the fraction of secretome mRNAs among ER-PUP tagged mRNAs dramatically exceeds that seen with PUP alone in tagging tiers 1, 2 and 3 (Supplementary Fig. 4E; see also Figure 2F, G), and is highest for the best-tagged RNAs in every abundance tier. Thus, the primary driver of ER-tagging is localization rather than abundance.

### Many but not all ER-enriched RNAs are bound by ER-proximal ribosomes

mRNAs tagged by ER-PUP are predicted to include ones translated at or near the ER. We compared mRNAs identified in ER-tagging with those detected in proximity-specific ribosome profiling experiments that had used the same Sec63p anchor^21^ (termed “ER profiling”). ER-profiling identifies ribosome-bound mRNAs near an ER-anchored biotin ligase that biotinylates Avi-tagged ribosomes^21^. 79% of mRNAs associated with Sec63-proximal ribosomes were also preferentially tagged by ER-PUP (4.7-fold enriched, *p*-value = 5.6 e^-415^), and only 0.2% were tagged better by PUP alone than ER-PUP (70-fold depleted, *p*-value = 7.9 e^-69^, Figure 3A). However, a sizable fraction (43%) of ER-PUP-enriched mRNAs were not identified via ER ribosome profiling (Figure 3A). By contrast, ER-PUP enriched mRNAs were depleted in mRNAs identified by mitochondria-specific ribosome profiling^22^ (1.7-fold depleted, *p*-value = 7.3 e^-6^, Supplementary Fig. 5A). Tagging also uniquely identified 453 mRNAs and 50 non-coding (nc) at the ER, while profiling detected 169 mRNAs not detected by tagging (Figure 3A, left panel, and Supplementary Fig. 5B). Thus, many but not all tagged RNAs were detected as translated at the ER under normal conditions.

ER-tagging detects RNAs seen in profiling only when ribosomes are arrested. Cycloheximide blocks translation elongation and so likely increases the time that ribosome-nascent chain complexes are near the ER-anchored biotin ligase in profiling ^21,39^. mRNAs enriched by ER-tagging include those that are enriched by profiling only when ribosomes are trapped by cycloheximide treatment (Figure 3B). We suspect that tagging detects ribosome-bound mRNAs that have only brief interactions with the ER, and so can be detected independent of elongation arrest (Figure 3B). In that sense, tagging is more sensitive, detecting transient proximity to the ER.

We next analyzed the relationship between U-tag length and ribosome profiling. mRNAs detected by Sec63p-mediated profiling^21^ were grouped into five tiers using k-means clustering, from highest ribosome association (Tier 1) to least (Tier 5, Supplementary Fig. 5C). mRNAs with the longest U-tags were more likely to associate with ER-proximal ribosomes (Tiers 1-3, *p*-value = 1.2e^-40^), and mRNAs were tagged regardless of their rank in profiling (Figure 3C). The most robustly ER-tagged mRNAs (Tiers 1 and 2) were more engaged with ribosomes, as inferred from profiling; poorly tagged RNAs progressively decreased in their associations with ribosomes^21^ (Figure 3D). As expected, ER-tagged RNAs did not exhibit this correlation with RNAs translated by mitochondria-proximal ribosomes, as inferred from Om45p-mediated profiling^22^ (Figure 3E).

Taken together, these findings indicate that tagging identified RNAs with ER-ribosome association, as well as RNAs that do not. Thus, tagging and profiling yield overlapping but non-identical sets of RNAs, and together provide a more complete view of RNAs that encounter the ER than does either approach alone.

### Mitochondria-localized RNA tagging

To detect RNAs near the mitochondrial outer membrane, we inserted PUP downstream of the *OM45* gene, resulting in an Om45p-PUP-2 chimera, which we refer to as “Mito-PUP” hereafter. Om45p is predicted to be co-translationally inserted into the mitochondrial membrane with its C-terminal PUP-2 domain in the cytosol^40-45^ (Figure 4A). GFP from Mito-PUP co-localized with the Tom70p-mCherry mitochondrial marker in yeast, and both proteins yielded fluorescence patterns comparable to those of the endogenous proteins fused to GFP^34^ (Figure 4B).

**Figure 4.**
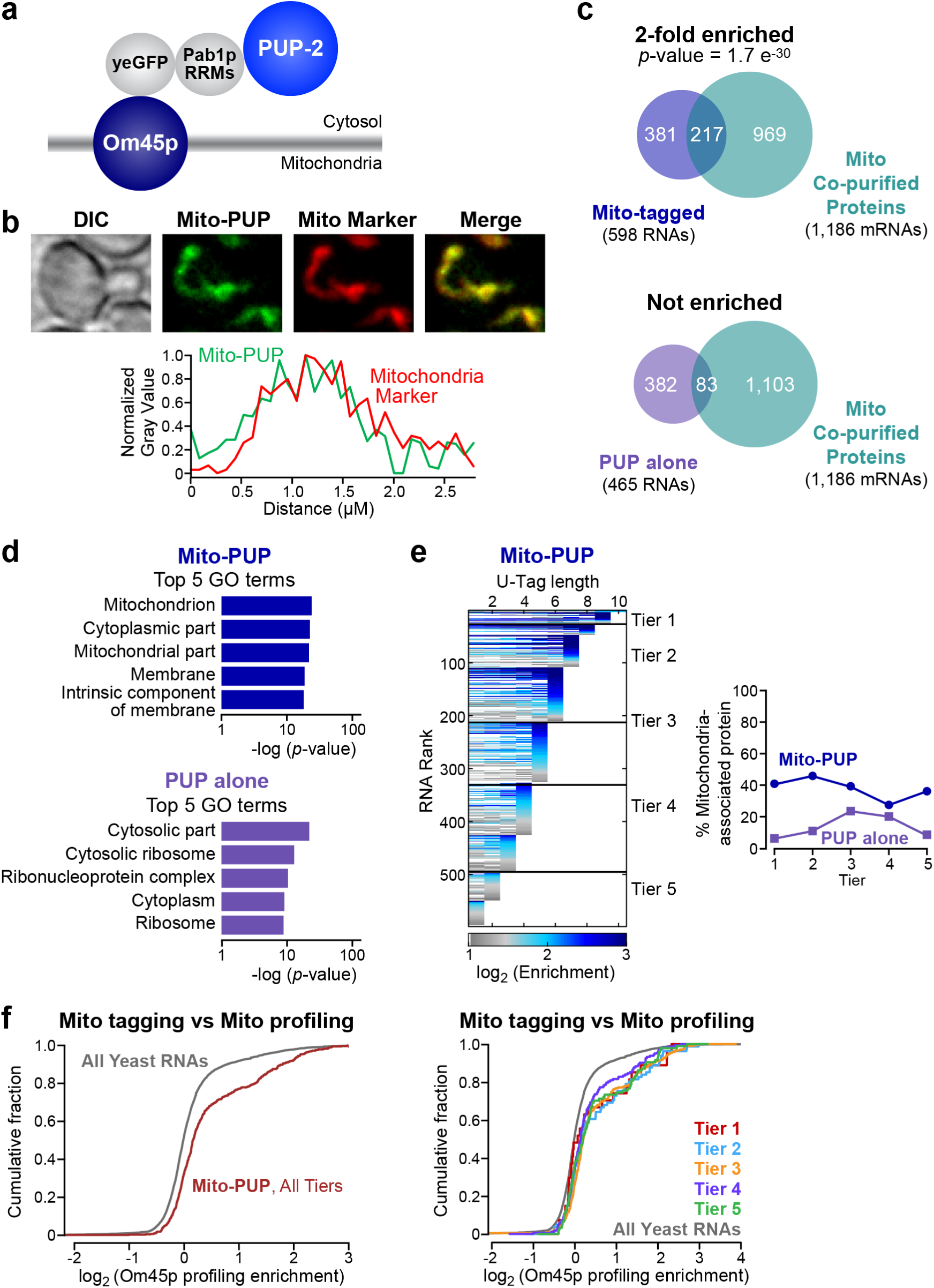
RNA contacts with the mitochondrial outer membrane. **a)** Architecture of Mito-PUP. PUP alone was fused to the cytosolic C-terminal end of the mitochondrial membrane protein, Om45p.^40-45^ **b)** Confocal localization of Mito-PUP fluorescence (green) and the mitochondrial marker, Tom70p-mCherry^34^ (red), including fluorescence intensity across a line that bisects the cell (below). **c)** Overlaps of Mito-PUP- and PUP alone-enriched mRNAs, and those that encode proteins that co-purify with biochemically isolated mitochondria.^46,47^ **d)** Top five gene ontology (GO) associations for Mito-PUP (top) and PUP alone (bottom). **e)** Rank of Mito-PUP-enriched RNAs plotted across each of ten U-tag lengths (left). The fraction of the RNAs in each Mito-PUP (blue line) and PUP alone tier (purple line) that encode mitochondrial proteins is plotted on the right. **f)** Cumulative fraction of all (left) and per-tier (right) Mito-PUP-enriched mRNAs with a given degree of mitochondrial profiling^22^ enrichment.

Mito-PUP preferentially tagged mRNAs that encode proteins physically associated with mitochondria and mitochondrial functions. Compared to PUP alone, Mito-PUP tagged 598 RNAs at least two-fold more efficiently (mitochondria-enriched), and 465 RNAs less efficiently (mitochondria-depleted, Figure 4C, and Supplementary Fig. 6). The entire set of RNAs that were preferentially tagged by Mito-PUP had statistically significant association with mitochondria-related GO terms^37,38^, and for mRNAs that encode mitochondria-associated proteins^46,47^ (2-fold enriched, p = 1.7e^-30^, Figure 4C, D, Table 2). mRNAs depleted from mitochondria by our DESeq2 analyses did not have these associations (Figure 4C, D). RNAs in the highest tagging tiers were more likely to encode mitochondrial proteins than those in lower tiers (Figure 4E, and Supplementary Fig. 7A). The efficiency of tagging at mitochondria also correlated with mitochondrial translation at that location, as judged by ribosome profiling^22^ (3-fold enriched, *p*-value = 1.8 e^-34^, Figure 4F, and Supplementary Fig. 7B). Together, these analyses strongly suggest that Mito-PUP preferentially tags mRNAs near mitochondria.

Despite the overlap between tagging and profiling, each method identified a unique set of mRNAs. Tagging and profiling detected 133 mRNAs in common, and these were enriched for mitochondrial functions by GO analysis (Table 3). The rank of commonly detected RNAs greatly differed between methods; for example, they trended toward longer uridine tags (Tiers 1-3) but tagging rank was reduced among RNAs detected most efficiently in ribosome profiling (Tier 1, Supplementary Fig. 7C, D). Instead, the common RNAs were distributed nearly evenly across the mid-range ribosome profiling tiers (Tiers 2-4), which likely indicates that differences in detection requirements influence the rank for each method (Supplementary Fig. 7C, D).

Mito-PUP tagged hundreds of RNAs that were not identified by ribosome profiling. Of these, 457 were mRNAs, and eight were ncRNAs (Supplementary Fig. 7E). Conversely, profiling identified 325 mRNAs that were not detected by tagging, but these were mostly lower abundance RNAs (Supplementary Fig. 7E, F). Of the RNAs uniquely identified by each method, those unique to tagging were associated with ion transport processes, and those unique to profiling were associated with tRNAs and respiration (Tables 4 & 5). Thus, tagging and profiling yield unique, but complementary results.

mRNAs localized to the outer periphery of mitochondria fall into two classes: Class 1, which require Puf3p for localization, and Class II, whose localization is independent of Puf3p.^48^ A third group of mRNAs, termed Class 3, are translated in the cytoplasm, not near mitochondria.^48^ Mito-PUP tags mRNAs localized near mitochondria, whether they require Puf3p for localization (Class 1, 3.4-fold enriched, p = 6.0e^-24^) or not (Class 2, 3.5-fold enriched, p = 3.1e^-22^). In contrast, MitoPUP tags Class 3 mRNAs poorly, indicating those mRNAs are too far from the Om45p anchor to be tagged. Thus, localized tagging specifically discriminates localization among classes of RNAs whose proteins are destined for the same organelle.

### RNAs at both ER and mitochondria in yeast and human cells

Most RNAs were tagged by either ER-PUP or by Mito-PUP (Figure 5A), and so yielded GO enrichments anticipated for that organelle (Tables 6 & 7). For example, several components of the TOM (Tom70p, Tom40p. and Tom71p) and TIM (Tim18p, Tim50p, Tim22p, Tim44p, and Tim54p) protein import complexes exhibit high tagging only by Mito-PUP, while those that encode certain secreted proteins (e.g., Ecm14p and Pff1p), ER-resident chaperones and translocon components (e.g. Kar2p, Ssh1p, and Sec63p), and the ncRNA of the SRP particle, *SCR1*^*49*^, were tagged only at the ER (Figure 5B).

**Figure 5.**
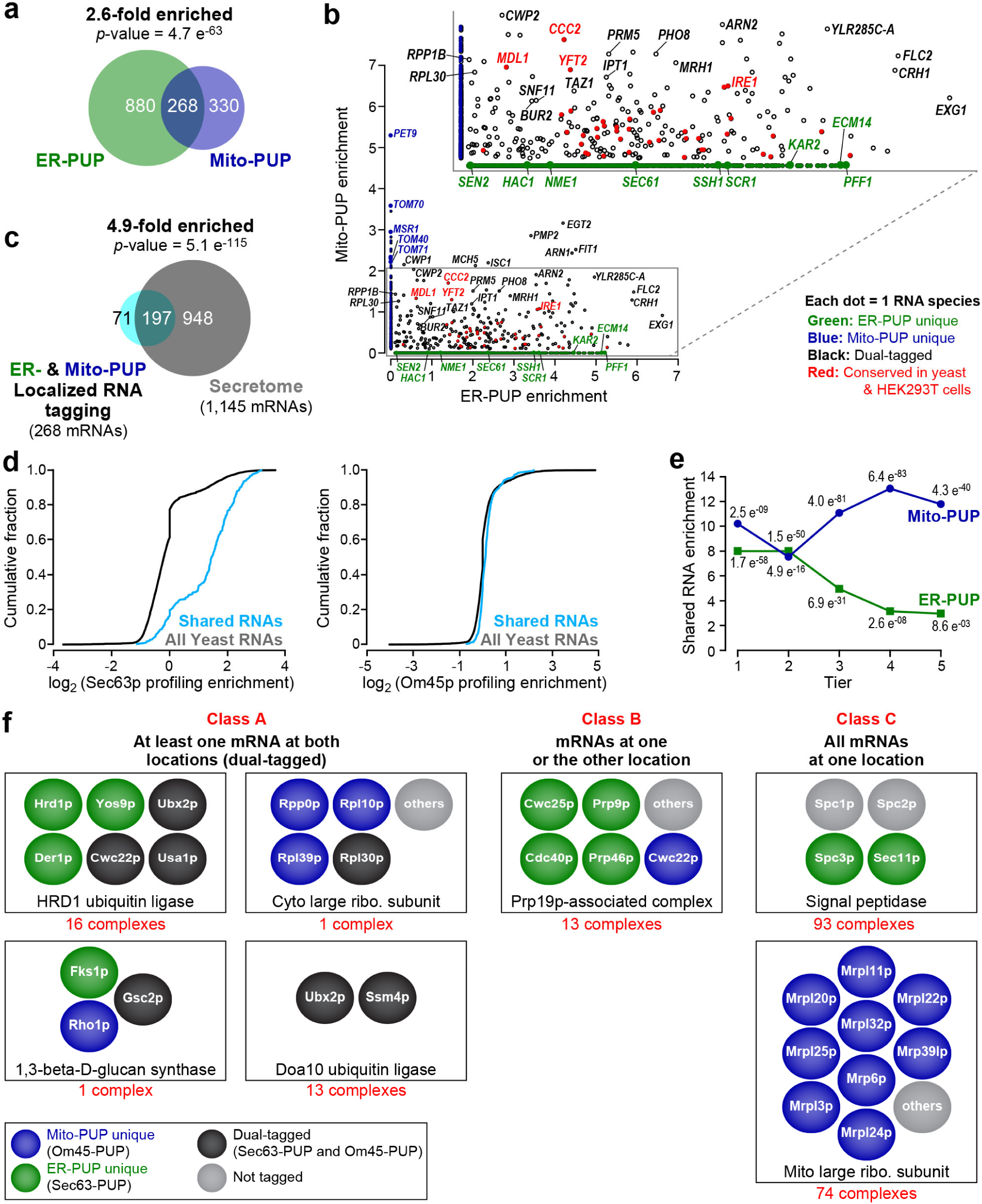
Dual tagging at both ER and mitochondria. **a)** mRNAs tagged by ER-PUP and Mito-PUP. **b)** Comparison of RNAs uniquely enriched by the ER- (green dots) or Mito-PUP (blue dots), and those that were common to both (black, and red). Larger circles represent mRNAs that encode proteins typical of each organelle, while the red and purple dots denote mRNAs that are detected at the ER (ER-PUP unique) and mitochondria (Mito-PUP unique) by independent, APEX-seq studies in HEK293T cells.^25^ **c)** Dual-tagged RNAs vs secretome mRNAs. **d)** Cumulative fraction of common RNAs that have ER-profiling^21^ (left) and mitochondrial profiling^22^ (right) enrichment. **e)** Distribution of common RNAs among ER- (green line) and Mito-PUP (blue line) rank. **f)** mRNAs that encode subunits of multiprotein complexes exhibit three localization patterns. Representative complexes depicted for each class; physical interactions between proteins not implied. For complete list, see Table 11.

Strikingly however, a substantial fraction (ER: 23%, Mito: 44%) of all tagged mRNAs were detected by both ER-PUP and Mito-PUP (p-value = 4.7 e^-63^; Figure 5A). Tagging of these mRNAs with both Om45p and Sec63p anchors suggests these mRNAs come near both the ER and mitochondrial outer membrane. As a whole, the dual-tagged mRNAs encoded primarily secreted proteins (4.9-fold enriched, p-value = 5.1e^-115^, Figure 5C, Table 8), were translated (Figure 5D) and received longer tails at the ER (Figure 5E), suggesting a longer cumulative time at that location.

A second group of 45 shared RNAs were tagged roughly equally by ER- and mitochondria-anchored proteins (normalized to PUP alone in each location; Figure 5B, and Table 9). High-ranking, shared mRNAs of this type include ones associated with lipid biosynthesis (*ISC1, IPT1, YFT2*, and *TAZ1*), ion transport (*MDL1, MCH5*, and *CCC2*), RNA polymerase II transcription (*SNF11* and *BUR2*), a plasma membrane-associated proteolipid (*PMP2*), and a GPI-anchored cell wall endonuclease (*EGT2*) (Figure 5B, and Table 9). These commonly-enriched mRNAs may reside where the two organelles are in close proximity^50,51^ or move from one location to the other (see Discussion).

Localization of specific RNAs to the proximity of both ER and the outer mitochondrial membrane is conserved. We compared our tagging results to data recently reported from human embryonic kidney (HEK293T) cells using APEX-Seq^25^. The high fraction of RNAs tagged at both locations in yeast (ER: 268 of 1,148; Mitochondria: 268 of 598, Figure 5A, B) was mirrored in HEK293T cells^25^, as was the identity of many of the RNAs (ERM: 50%; OMM: 67%, Supplementary Fig. 8A, B, C). Among the dual-tagged RNAs detected in both yeast (Figure 5B) and human cells (Supplementary Fig. 8C) were ones that encode functions linked to ERMES^51^ (ER-mitochondria encounter structure), formed where ER and mitochondria are in close proximity. These included mRNAs that encode proteins and functions that are associated with MAMs^50,52^ (ER-mitochondrial associated membranes) including transmembrane transporters (Figure 5B and Supplementary Fig. 8C, and Table 10), and proteins involved in lipid (*CAX4, LAC1, TGL1*, and *ALE1*) or glycoprotein (*ROT2, CAX4, OST6*) metabolism. Strikingly, the ER stress sensor, *IRE1/ERN1*^*53-59*^ was detected at both organelles, consistent with the presence of Ire1p protein at both organelles^52^, and the requirement for Ire1p for the increase of mitochondrial respiration during the UPR^60^ (Figure 5B and Supplementary Fig. 8C, and Table 10).Conservation of many shared targets suggests the importance of their presence at both locales, particularly given the substantial differences between the two techniques.

mRNAs that encode components of multi-protein complexes (identified through GO annotations^37,38^) comprise three classes based on the range of locations between ER and mitochondria (Figure 5F, Table 11). Thirty-one complexes fell into Class A, and included dual tagged mRNAs (Figure 5F, Table 11; e.g. HRD1 ubiquitin ligase complex). Most strikingly, thirteen complexes fell into Class B, with mixtures of mRNAs tagged at one or the other location (Figure 5F; e.g. Prp19 complex). Ninety-three complexes were uniquely enriched by ER-PUP (e.g. the signal peptidase) and seventy-four by Mito-PUP (Class C; Figure 5F, Table 11, e.g. the 54S mitochondria ribosomal subunit). The presence of mRNAs for subunits in different locales suggests coordination to assemble the complexes, or that certain of the proteins have other roles.

### Regulatory sites and RNA-binding proteins

RNA localization often is controlled by RNA-binding proteins (RBPs). We therefore identified sequence motifs in the 3’UTRs of RNAs identified by localized tagging using MEME^61^, and compared those motifs to known RNA-binding specificities^26,62^.

The top motif among RNAs tagged by Mito-PUP was a degenerate form of the Puf3p binding site (Supplementary Fig. 9A). Puf3p binds and controls nuclear-encoded mRNAs with mitochondrial functions, and participates in their localization near mitochondria^26,30,48,63-66^. The proportion of mito-tagged RNAs with such elements was similar across all tagging tiers (Supplementary Fig. 9B, dark blue line). This prompted re-analysis of Om45p-mediated profiling data^22^), which revealed the same phenomenon (Fig 9A, B, light blue line). Our findings suggest that Puf3p promotes mitochondrial localization of only certain mRNAs, and that others arrive there through other mechanisms^48^.

ER-tagged RNAs were enriched for Bfr1p targets (*p*-value = 1.9 e^-280^), consistent with the role of Bfr1p in the secretory pathway and RNA metabolism^26,67-71^; similarly, mitochondria-tagged RNAs were enriched for Puf3p targets (*p*-value = 1.2 e^-26^) (Supplementary Fig. 9C). ER-tagged RNAs were moderately enriched among RNAs that bind Pub1p (*p*-value = 2.8e^-36^), Mrn1p (*p*-value = 4.5e^-16^), and Scp160p (*p*-value = 4.6e^-11^); of these, only Scp160p is known to localize to the ER^72,73^ (Supplementary Fig. 9D). Sec63-tagged mRNAs revealed AU- or U-rich motifs (Supplementary Fig. 9E). These analyses point to proteins likely involved in control of these mRNAs.

### Non-coding RNAs and RNA metabolism

Analysis of tagged RNAs revealed connections between RNA metabolism and cell biology. ncRNAs from 53 genes were significantly enriched at mitochondria or the ER relative to the PUP alone control (Figure 6; three representative paired reads are shown for each type of RNA, and Table 12). *SCR1*, the RNA component of the Signal Recognition Particle^49^, was the ncRNA tagged best by ER-PUP (Figure 5B, and Figure 6A); *HAC1* mRNA, which is critical in the UPR^53-59^, also was detected (Figure 6A; see also Figure 5B).

**Figure 6.**
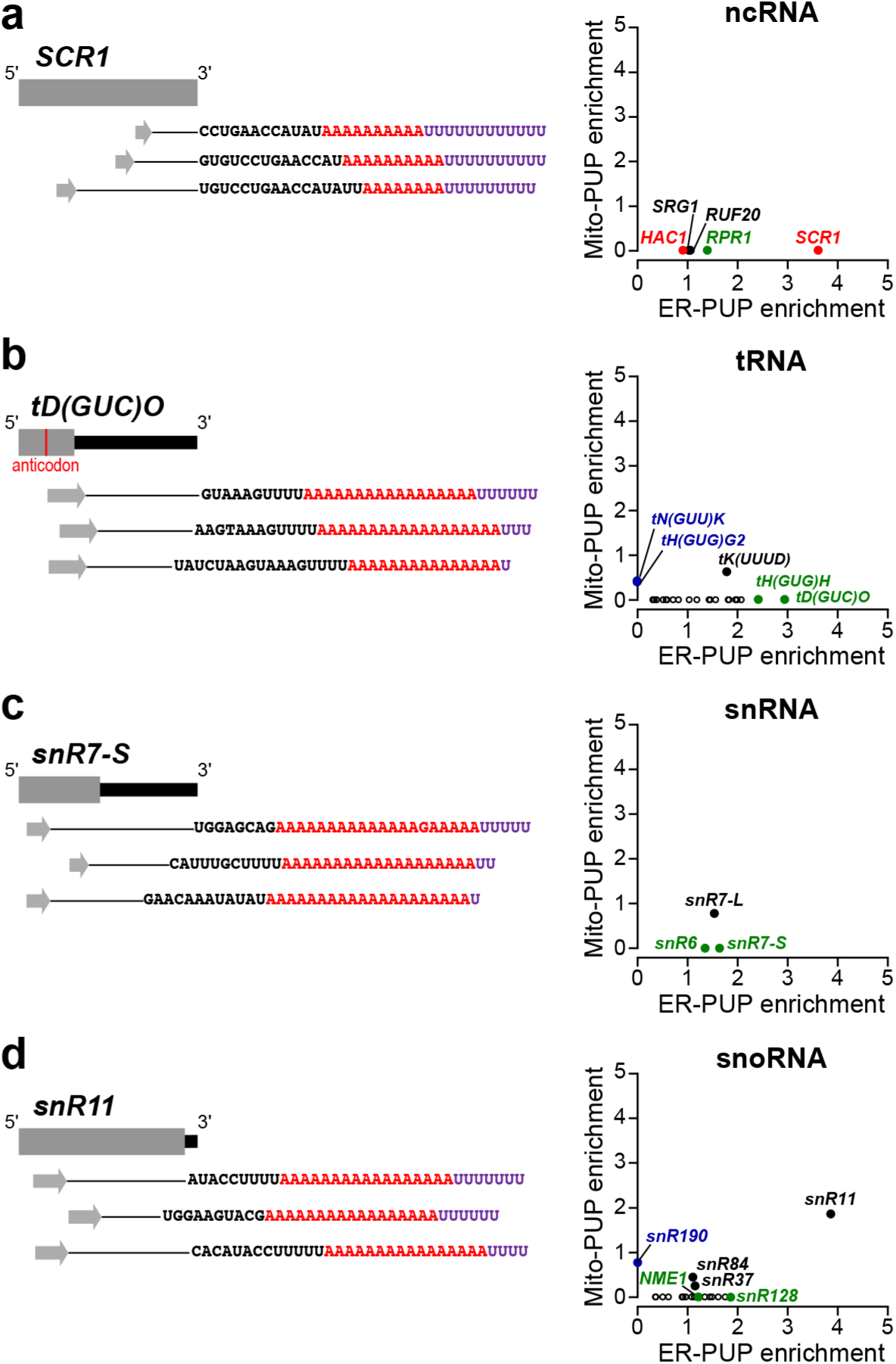
ncRNAs enriched at the ER and mitochondria. ER- and Mito-PUP scatter analyses on right demonstrate the enrichment distributions for each of the four classes of non-protein coding RNA genes, which includes RNAs derived from **a)** ncRNAs, **b)** tRNAs **c)** snRNAs, and **d)** snoRNAs. Each RNA is represented by a single dot, and the dots are colored to reflect unique enrichment by ER-PUP (green) or Mito-PUP (blue), or both (black). Larger circles highlight notable RNAs such as the top RNAs for each site or, for the ncRNAs, RNAs characteristic of the ER (red dots). Three RNAs from each class are diagrammed to the left. Forward reads are represented by gray arrows, while black sequences represent the DNA-encoded bit, presumably the 3’ end of the RNA, detected in the reverse read. This is followed by the sequence of non-templated adenosines (red A’s) and uridines (purple U’s) that followed the 3’ sequence. The line in between these two features represents the section of the RNA inferred from the mapped pairedend reads.

21 tRNA-related transcripts were tagged at the ER. These lacked CCA, contained 3’ extensions beyond the 3’ end of the mature tRNA, and possessed poly(A) tails upstream of the uridines added in tagging (Figure 6B). Poly(A) is added to some ncRNAs as part of nuclear RNA surveillance, mediated by the TRAMP complex and exosome^74-77^. We obtained no reads upstream of anticodons, which may be due to the presence of modified bases that arrest reverse transcriptase^78-80^ or RNA cleavage events that may leave tRNA halves^78,81^. Among proteins that participate in tRNA splicing^82^, mRNAs encoding Sen2p, a subunit of the endonuclease, were detected at the ER; while those encoding Tom70p, an enhancer of tRNA splicing *in vivo*,^83^ was detected at mitochondria, where it also is part of a translocase (Figure 5B).

25 snRNA- and snoRNA-related RNAs were tagged by either ER- or Mito-PUPs. These included snRN7-L, which was tagged at both organelles (ER: Tier 3, Mito: Tier 4), and snR7-S and snR6, which were only tagged at the ER (Figure 6C). Yeast snRNAs can shuttle to the cytoplasm as part of their maturation, perhaps to help prevent inclusion of misprocessed snRNAs in the spliceosome^84^. Some snoRNAs were also tagged at one or both sites, including *NME1*, the RNA component of MRP that catalyzes RNA cleavage events^85^ (Figure 6D; see also Figure 5B). While snoRNAs are thought to be restricted to the nucleus, failure to properly process their guanosine cap can cause their accumulation in the cytoplasm^86^. Together, the data on ncRNAs suggest tagging captures mature ncRNAs as well as ones undergoing maturation and surveillance.

## DISCUSSION

Localized RNA Tagging identifies RNAs located to specific sites transcriptome-wide and is independent of hybridization, affinity purification, fractionation, cross-linking, or chemical treatments. RNAs tagged using Sec63p or Om45p displayed distinct properties consistent with each anchor. The number of uridines added to an individual RNA molecule likely reflects the integrated time it was at that location. mRNAs with longer U-tags were enriched for biological functions consistent with their location and association with local ribosomes. These findings mirror analyses of RNA-protein interactions using tagging, in which the number of uridines added to an individual RNA molecule correlated well with its binding affinity for the protein, as determined *in vitro*.^26,29^ Other variables, such as the structure of specific RNAs, may also effect tagging efficiency.

Tagging provides a cumulative “record” of an RNA’s movements, while the recently described APEX-Seq^25^ approach yields a snapshot – a “registry” of RNA locations at a specific time. The two approaches are complementary and provide a fuller view of RNA movements than either alone. Similarly, localized tagging and global studies of RBPs are synergistic (e.g., Supp. Figure 9). For example, Puf3p binds to and controls expression and localization of nuclear-encoded mitochondrial mRNAs.^26,30,48,63-66^ Mito-tagged RNAs often contained suboptimal Puf3p binding sites, consistent with the existence Puf3p-independent mechanisms for mitochondrial localization. Comparison with profiling suggests other roles for optimal binding, perhaps including its ability to enhance rather than repress translation^87^ (Supp. Figure 9B, C). ER-tagged RNAs contained novel motifs, some of which may be involved in the SRP-independent localization of mRNAs to the ER^88^ or may bind RBPs, such as Bfr1p. ^26,71^ Indeed, Bfr1p is implicated in the secretory pathway, co-purifies with secretory mRNPs and mRNAs, and localizes to the ER in a manner that requires its RNA binding activity.^26,67-71^

A specific and large subset of RNAs were tagged using both ER (Sec63p) and mitochondrial (Om45p) anchors. These “dual-tagged” RNAs might arise in three ways. First, they may be located where the two organelles are very close, perhaps as promoted by ERMES^51,89^ (ER-mitochondria encounter structure). Second, a single RNA molecule may move between organelles. Certain dual-tagged RNAs are tagged poorly at one or both locations, suggesting transient interactions, much as RNAs tagged poorly by an RBP possess poor (if any) cognate binding sites^26^. Third, individual RNA molecules may go from the nucleus to either one or the other location, rather than between the two organelles. The mechanism of dual localization we detect appears to be largely independent of translation, since dual localization is very rare among mRNAs detected by yeast ribosome profiling at these two organelles. ^21,22^ However, mRNAs detected by localized tagging might be translated, but in a manner that escapes detection in profiling, which requires ribosomes with their exit tunnels near the membrane^21,22^. Underlying mechanisms may be identified through use of multiple tagging devices in the same cell, live imaging, ^8,18-20,90^ or strategies that track localization and translation simultaneously^91,92^.

Dual localization of specific mRNAs is conserved between yeast and human cells, and may reflect proximity to MAMs. Dual localization may help integrate events between ER and mitochondria. IRE1 mRNA is exemplary, and is dual-localized in both yeast and human cells. Its dual localization may integrate inputs from both organelles and facilitate coordinated responses, such as the Ire1p-dependent rise in respiration after ER stress in yeast^60^ or sustained UPR-induced apoptosis in mammalian cells^93^.

Non-coding RNAs were readily detected in tagging, and include *SCR1 and NME1*, the RNA components of SRP^49^ and MRP^85^. Indeed, SCR1 was the most strongly tagged ncRNA at the ER. Tagged RNAs related to tRNAs, snRNAs and snoRNAs appear to have been caught during their maturation or surveillance. Tagged tRNAs were 3’-extended and polyadenylated upstream of the U-tag. Their polyadenylation suggests action of the nuclear TRAMP complex prior to their encounter with the tagging enzyme. 3’-extended tRNAs may subsequently be processed by the endonuclease Trz1p^94^, which is both nuclear and mitochondrial^34,94,95^. The presence of tagged snRNAs supports the recent finding that yeast snRNAs are shuttled to the cytoplasm as part of their maturation to prevent inclusion of misprocessed snRNAs in the spliceosome^84^. Tagged snoRNAs may arise through a different form of surveillance, as failure to properly process the guanosine cap of snoRNAs leads to their accumulation into the cytoplasm^86^.

Tagging can be performed in living organisms, as is true of “TRIBE,” in which ADAR-catalyzed deaminations mark the binding of specific proteins^96,97^. APEX-seq, as currently configured, is not suitable in intact organisms. Conversely, in cells with endogenous uridylation activities, application of tagging would be simplified through enzymes with different nucleotide specificities^98^ or by reduction of endogenous uridylation activities.

The use of localized devices to covalently mark RNAs provides an entrée to creating histories of individual RNA molecules. Combinations of Localized RNA Tagging, APEX-Seq, or TRIBE, performed in the same cell, will yield biographies of individual RNAs, in which their movements and protein encounters are written in unique combinations of modifications and tail sequences.

## MATERIAL AND METHODS

### RNA extraction and library preparation

Total RNAs and libraries were prepared in accordance with previous reports^26^. Sequencing libraries were submitted to the UW-Madison biotechnology center for paired-end sequencing (2X50 bp) on the HiSeq2500 platform. Libraries were loaded at equal concentrations with PhiX loaded at 30% of the total concentration.

### Mapping reads to the genome

FastQ files were processed with a custom sequencing pipeline^26^. Reads from libraries that were sequenced on multiple lanes were pooled and processed together.

### Processing read counts

Read counts were processed differently to determine either (A) absolute RNA tagging values or (B) tagging enrichment at a given site.

#### A) Calculation of tagged reads per million across ten minimum U-tag lengths

RNAs species that were tagged reproducibly across three replicates were identified with BioVenn^99^ (http://www.biovenn.nl/). The Read counts in each replicate were then normalized to the total number of reads (in millions) for that replicate, yielding the tagged reads per million (TRPM) across ten minimum U-tag lengths (1U to 10U) for each tagged RNA species. The TRPM values for each RNA species in each replicate were then averaged to yield the average TRPM (TRPM Avg.) for each RNA species at each of the U-tag length levels. RNAs species were then ranked by U-tag length and reads, and RNAs with the longest U-tag length reads, but less total reads having priority over those with more reads but shorter U-tag length.

#### B) Deseq2 analyses to determine enrichment at the ER and Mitochondria

Raw read counts for RNAs detected in each experimental replicate were analyzed with Deseq2^35^. For each individual analysis (ER- or Mitochondria-localized RNA tagging data), the PUP alone (+PAB) read counts were used as a control. In all our analyses, a log_2_ fold change of two-fold or greater (log_2_(FC ≥ 2)) and an adjusted p-value (*p*-adj) cutoff of < 0.05 was applied. RNAs that met these criteria were considered “enriched (i.e. “ER-enriched”) or depleted (i.e. “ER-depleted” or PUP “alone-enriched”).

### K-means clustering to generate tiers and heatmaps

Clustering was done using Cluster 3.0^100^ (C clustering Library 1.52) using the K-means option as previously reported^26^. Five groups (Tiers) were arbitrarily selected, and log_2_-transformed data were clustered based on detection efficiency (i.e. L_2_FC). Tiers were ranked based on highest to lowest efficiency of detection (i.e. most enriched to least enriched), with priority given to enrichment at the highest U-tag lengths. Heat maps were generated with MatLab.

### Data Mining

#### Proximity-specific ribosome profiling

Proximity-specific ribosome profiling data were mined published experiments. ^21,22^ mRNAs with greater than or equal to two-fold or higher ribosome-protected fragment reads relative to the input were determined to be “enriched”. Since we omitted did not use cycloheximide in our experiments, we selected Sec63p^21^ (1 min biotin pulse) and Om45p^22^ (2 min biotin pulse) mediated profiling data that omitted the translation inhibitor.

#### Mitochondrial proteome mRNAs

Mitochondria-copurified proteins from two experiments^46,47^ were consolidated into a single list termed, “mitochondrial proteins”.

#### RBP targets

RNA that interact with RNA-binding proteins were mined from published RNA tagging^26^ and RIP-Chip^62^ data. RNA tagging data were reprocessed using the DESeq2^35^ approach described here for ER-Pup and Mito-PUP, also using PUP alone (+PAB) as a control set. RNAs that were tagged log_2_(Δ Tagged Reads) ≥ 1)) and significance (p-adj) < 0.05 relative to PUP alone (+PAB) were considered enriched. These analyses were done across ten minimum U-tag lengths, and the RNA were ranked by highest to lowest U-tag length enrichment. For RIP-Chip, RNAs that had a Significance Analysis of Microarrays (SAM) q-value < 10% were considered targets. The classes of Puf3p mRNA targets were retrieved from published biochemical experiments^48^.

#### APEX-Seq

APEX-Seq data^25^ as used for these analyses. Similar cutoffs applied to our data (log_2_(Fold Change) ≥ 1, p-adj < 0.05) were applied to ER (ERM) and mitochondrial (OMM) APEX-Seq data to facilitate comparison.

#### Dual-tagged RNA conservation

Human homologs of dual-tagged yeast genes were retrieved using YeastMine^37,38^ (https://yeastmine.yeastgenome.org/yeastmine/begin.do).

### 3’ UTR motif enrichment analyses

Command line MEME^61^ Version 5.0.4 was used for all analyses. The order (Rank) of all yeast 3’ UTRs from^26^ was randomized using the excel rand (=RAND()) function, and the resulting list was used as background (-neg) for all MEME analyses. Motifs that were enriched in the 3 UTRs of tagged RNAs (ranked by longest U-tag length) were done using the differential enrichment (-objfun de) function, and MEME was prometed to return the top ten motifs (-nmotifs 10) that range from 8-10 nts in length (-minw 8 -maxw 12) with all lengths between those limits included in the scan (-allw). The rank of the 3’ UTR within the list of tagged RNAs or the list of 3’ UTRs with randomized rank was taken into account (-norand) in the analyses. Puf3p binding element incidence in the 3’ UTRs of RNAs was determined using a custom perl script^31^.

### Tools used

#### Gene ontology (GO)

All GO analyses were done using yeast mine lists^37,38^ (https://yeastmine.yeastgenome.org/yeastmine/bag.do). The analyses used the default background, and considered enrichments with a maximum p-value of 0.05 after Holm-Bonferroni correction.

#### Venn Diagrams

Venn diagrams were generated using BioVenn^99^ (http://www.biovenn.nl).

#### Hypergeometric Distribution Analyses

Hypergeometric distribution calculations were done with the online calculator available from the Graeber Lab (https://systems.crump.ucla.edu/hypergeometric/). The total number of yeast transcripts used was 6,712 RNAs, as defined by RNA-seq^26^.

#### Cumulative Fraction Plots

Cumulative fraction plots were done in either RStudio (Figures 3D, E, 4F) (as previously reported^26^) or Excel (this report, Figure 5D).

#### Tab file conversion to Fasta

Tab files that contained 3’ UTR sequences were converted to fasta format using the HIV sequence database Format Converter (https://www.hiv.lanl.gov/content/sequence/FORMAT_CONVERSION/form.html)

#### Confocal Microscopy

Strains were grown to mid-log (OD_660_ 0.5-0.8) phase in 25 mL cultures at 30 °C in a horizontal shaker at 180 RPM. Cells were grown in synthetic complete^101^ (SC) Low fluorescence^102^ (LOFLO) media

Yeast cells were immobilized on Concanavalin A (ConA Sigma: 11028-71-0) coated coverslips for imaging with a modified version of a published imaging protocol^103^.

#### Coverslip Preparation

Coverslips were first incubated in a methanol/hydrochloric acid (1:1) solution in 50 mL conical tube in a fume hood overnight. The next day, coverslips were rinsed 3Xs with deionized water, and then placed on one edge inside 65 °C oven until fully dry (∼ 30 mins). The dry coverslips were then cooled to RT, and 400 uL of 2 mg/mL concanavalin A (ConA) was spread evenly in the center. After drying for 60 mins at RT, the coverslips were then tilted on their side to remove excess ConA. The coverslips were then covered and dried at RT overnight.

#### Fixation

Mid-log phase cultures were spun down, and resuspended in 500 uL of fresh SC LOFLO media. 100 uL of cells were then placed evenly on the center of the ConA-coated coverslip and allowed to bind for 30 mins in a 30 °C incubator without shaking. Once immobilized on the coverslip, the cells were fixed by submerging the coverslip in SC LOFLO (5% Formaldehyde, Fisher: BP531-500) for 20 minutes. After, the coverslip was washed with 3Xs with fresh SC LOFLO media, and the coverslip was then placed on a slide containing SC LOFLO.

#### Confocal settings

Cells were imaged on a Leica TCS SP8 confocal microscope using LAS X 3.1.1.15751. The microscope is equipped with a Photomultiplier (PMT) and Hybrid detectors (HyD). A 63x 1.4NA HC Plan Apochromat oil immersion objective was used with 3.01 zoom and standard scanner with 400Hz scanning speed. Z-stacks with a 0.3 uM step size were collected, and yeGFP (495nm - 530nm) and mCherry (600nm - 650nm) were sequentially imaged.

#### Fluorescence quantitation

Images were processed using FIJI^104^/ImageJ^105^ Version: 2.0.0-rc-69/1.52i. Noise reduction was done on the images using the “Despeckle” function. Fluorescence intensities (gray values) quantified using the measure (Ctrl+M) function along a straight line that was drawn across individual cells. The measurements were then normalized for each individual sample to get the Normalized Gray Value at each individual measurement. This was done with the following equation, Normalized Gray Value at a specific point on the line = (Raw intensity value at a specific point on the line – Minimum of all intensities on the line)/(Maximum of all intensities on the line – Minimum of all intensities on the line). The normalized Gray values for each sample were plotted on a line graph using Microsoft Excel.

## Yeast Strains (BY4742 background)

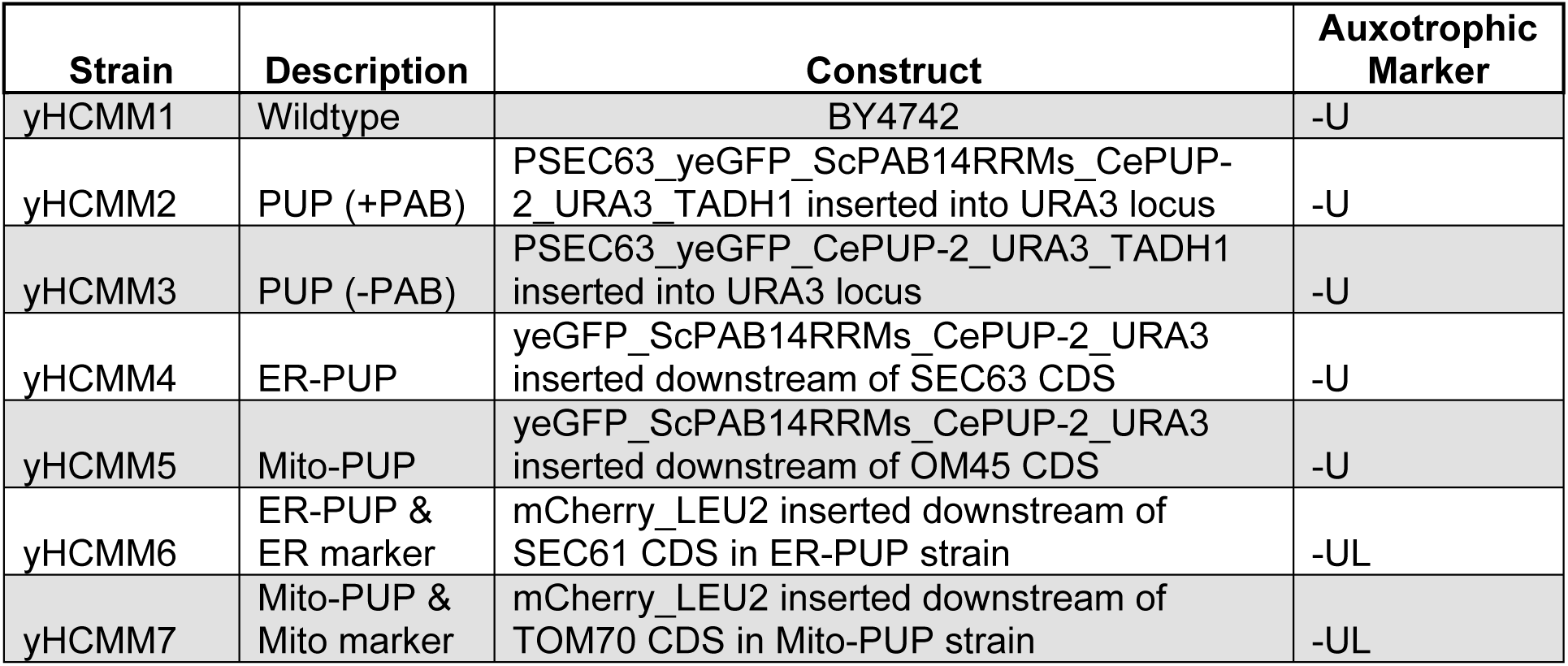

## Plasmids

| **Plasmid** | **Construct** | **Auxotrophic Marker** |
| --- | --- | --- |
| pHCMM1 | PTEF1_yeGFP_ScPAB14RRMs_CePUP-2_URA3_TADH1 | -UL |
| pHCMM2 | PSEC63_yeGFP_ScPAB14RRMs_CePUP-2_URA3_TADH1 | -UL |
| pHCMM3 | PSEC63_yeGFP_CePUP-2_URA3_TADH1 | -UL |
| pHCMM4 | PCYC1_SEC61_mCherry_LEU2_TCYC1 | -L |

## Primers

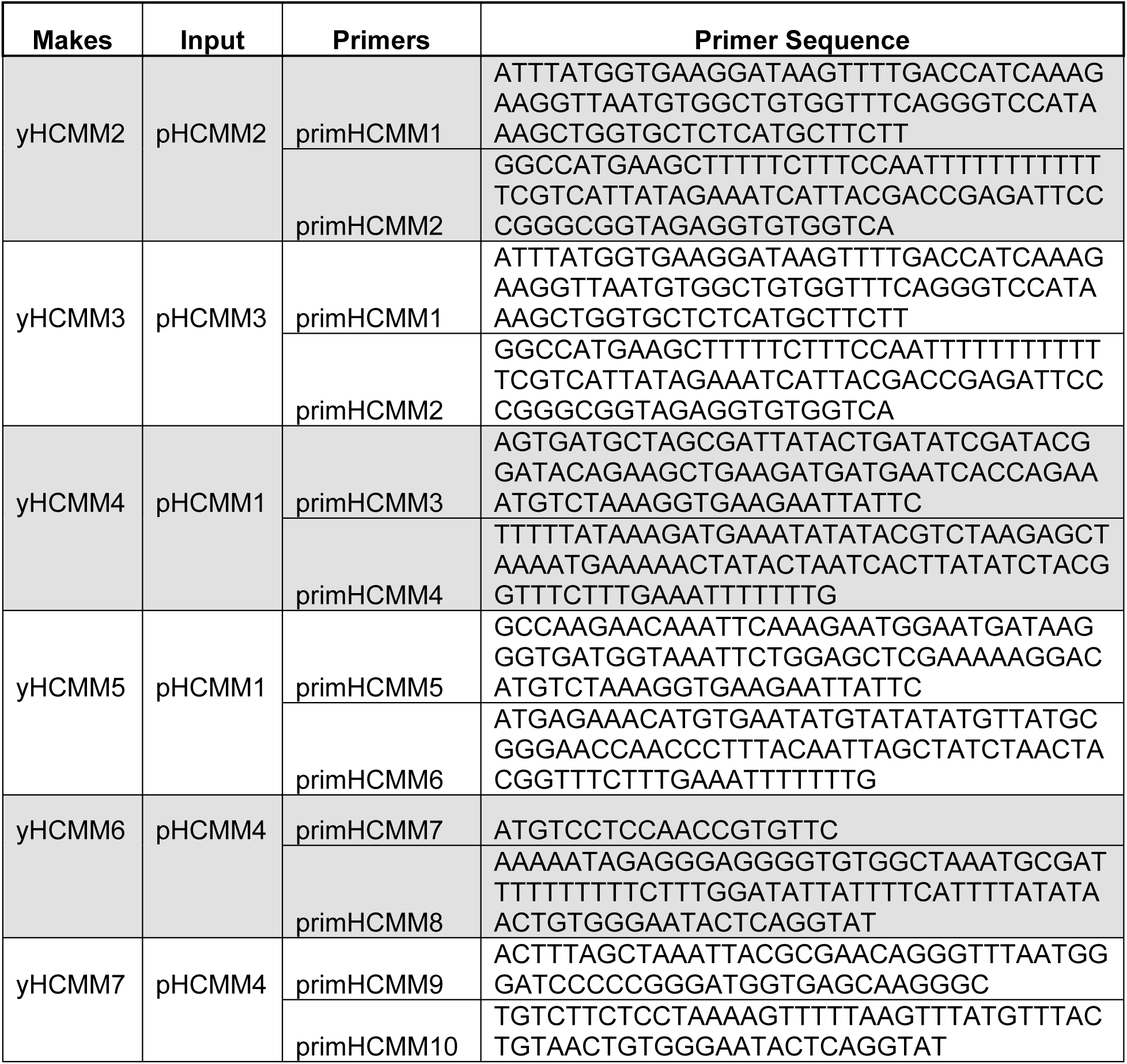

## ACKNOWLEDGMENTS

We thank the Wickens and Kimble laboratories, and members of the UW-Madison community, particularly Judith Kimble and Dave Brow, for helpful advice and discussions. We thank Eric Phizicky and Elizabeth Grayhack (University of Rochester), and Scott Aoki (Indiana University) for their insights. We are grateful to Sarah Crittenden for help with confocal microscopy, Kim Haupt and Elle Kielar Grevstad (UW-Madison Optical Core) for assistance with image analyses, and Aaron Hoskins, Tucker Carrocci, and Ian Norden for help with fluorescent proteins. We are grateful to Joshua R. Hyman, Molly Zeller, Amanda Maegli and Mike Sussman (UW-Madison Biotechnology Center) for help with sequencing, and to Laura Vanderploeg and the Biochemistry Media Lab for help preparing figures. This work was supported by an NIH grant to M.W. (GM50942) and an E.W. Hopkins Fellowship to H.C.M.M. While in UW-Madison, C.P.L. was supported by Wharton and Biochemistry Scholar Fellowships, and D.F.P. was supported by T32 GM008349 (NIH) and a WARF Scholarship.

## AUTHOR CONTRIBUTIONS

H.C.M-M. and M.W. conceived the method and designed experiments; H.C.M-M. performed all experiments, and analyzed the data; H.C.M-M. and M.W. interpreted results and wrote the paper and figures; C.P.L. provided advice on tagging and helped with revisions, and D.F.P. provided advice on protein design.

