## Supplementary Figures for "Records of RNA localization through covalent tagging"

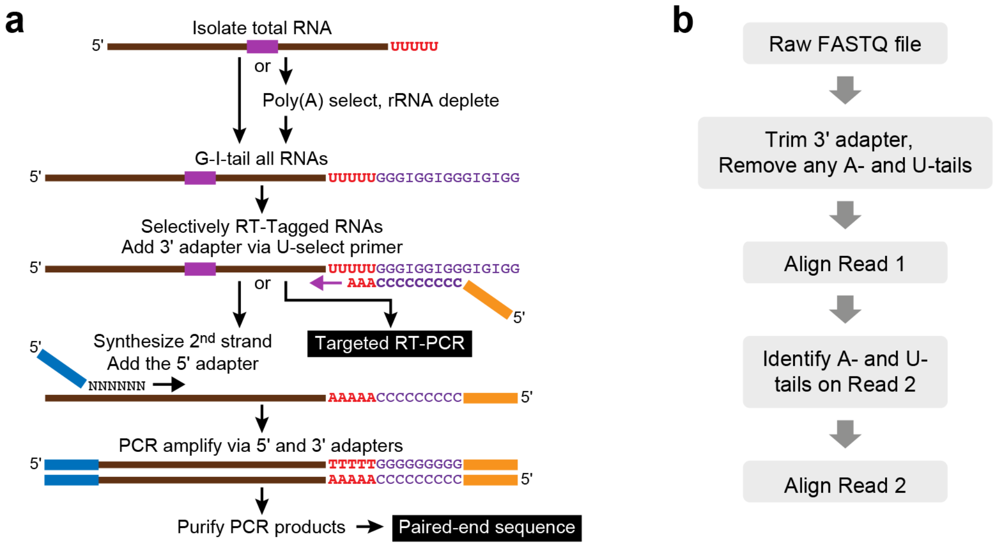


**c**

**PUP alone (+PAB)**

| Replicate | Total Reads | Uniquely Mapped Reads | Number of Genes Identified |
|---|---|---|---|
| 1 | 26,116,295 | 4,941,967 | 5,999 |
| 2 | 25,021,422 | 3,780,003 | 5,982 |
| 3 | 21,865,004 | 4,163,971 | 5,991 |

**PUP alone (–PAB)**

| Replicate | Total Reads | Uniquely Mapped Reads | Number of Genes Identified |
|---|---|---|---|
| 1 | 16,143,092 | 8,368,921 | 2,090 |
| 2 | 20,430,380 | 10,462,818 | 2,208 |
| 3 | 16,713,968 | 8,274,280 | 2,080 |

**Figure S1. Detecting U-tagged RNAs.**

**a)** *Experimental protocol.* From a preparation of total yeast RNA, RNAs with 3' terminal uridines were identified using sequential depletion of rRNAs and oligo(dT) selection, G/I-tailing, reverse transcription with a U-selective primer, and amplification.[26] Sequences of the 3' ends and tails were identified by paired-end sequencing (Illumina HiSeq2500) **b)** *Computational analysis.* Flowchart to identify RNAs tagged *in vivo.* **c)** *Representative data.* Sequencing statistics with PUP (+PAB) and PUP (-PAB). Experiments were performed with a minimum of three biological replicates and a minimum of two technical replicates for each one.

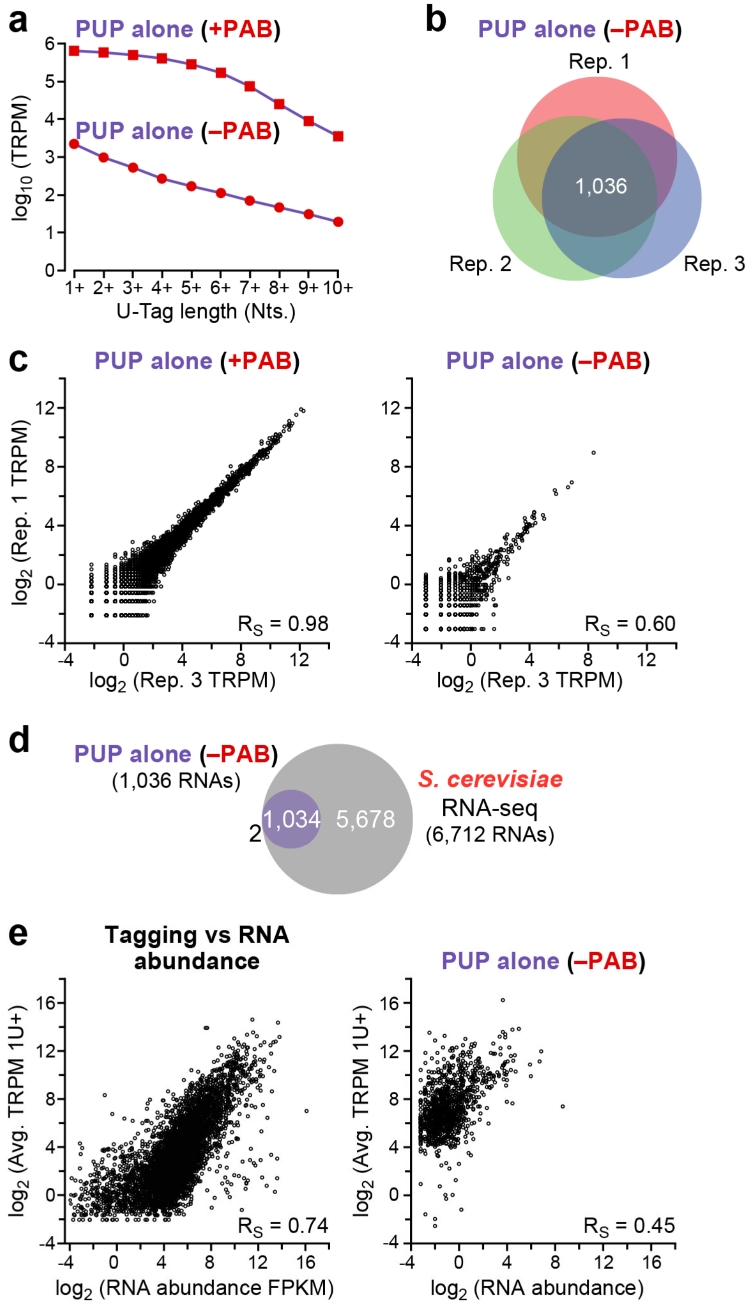


**Figure S2. PUP tagging efficiency.**

**a)** U-tag length intervals (1U-10U) and analogous PUP variant U-tag reads (TRPM)**. b)** Reproducibility across three PUP alone (-PAB) biological replicates. **c)** Reproducibility and abundances of PUP alone, with (left) or without (right) PAB, for individual tagged mRNAs. Individual RNA species (black dots) reads (TRPM) are compared across two experiments. **d)**

Comparison of the number of RNAs identified with PUP (-PAB) vs RNA seq.[26]. **e)** Tagged reads vs RNA abundance[26] for PUP alone with (left) or without (right) PAB.



**a** ER-PUP

| Replicate | Total Reads | Uniquely Mapped Reads | Number of Genes Identified |
|---|---|---|---|
| 1 | 15,779,019 | 4,455,578 | 5,285 |
| 2 | 20,712,699 | 4,716,985 | 5,684 |
| 3 | 15,357,796 | 4,345,423 | 5,506 |

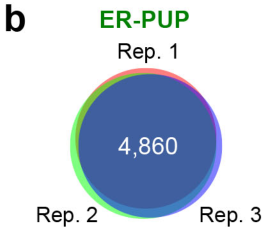


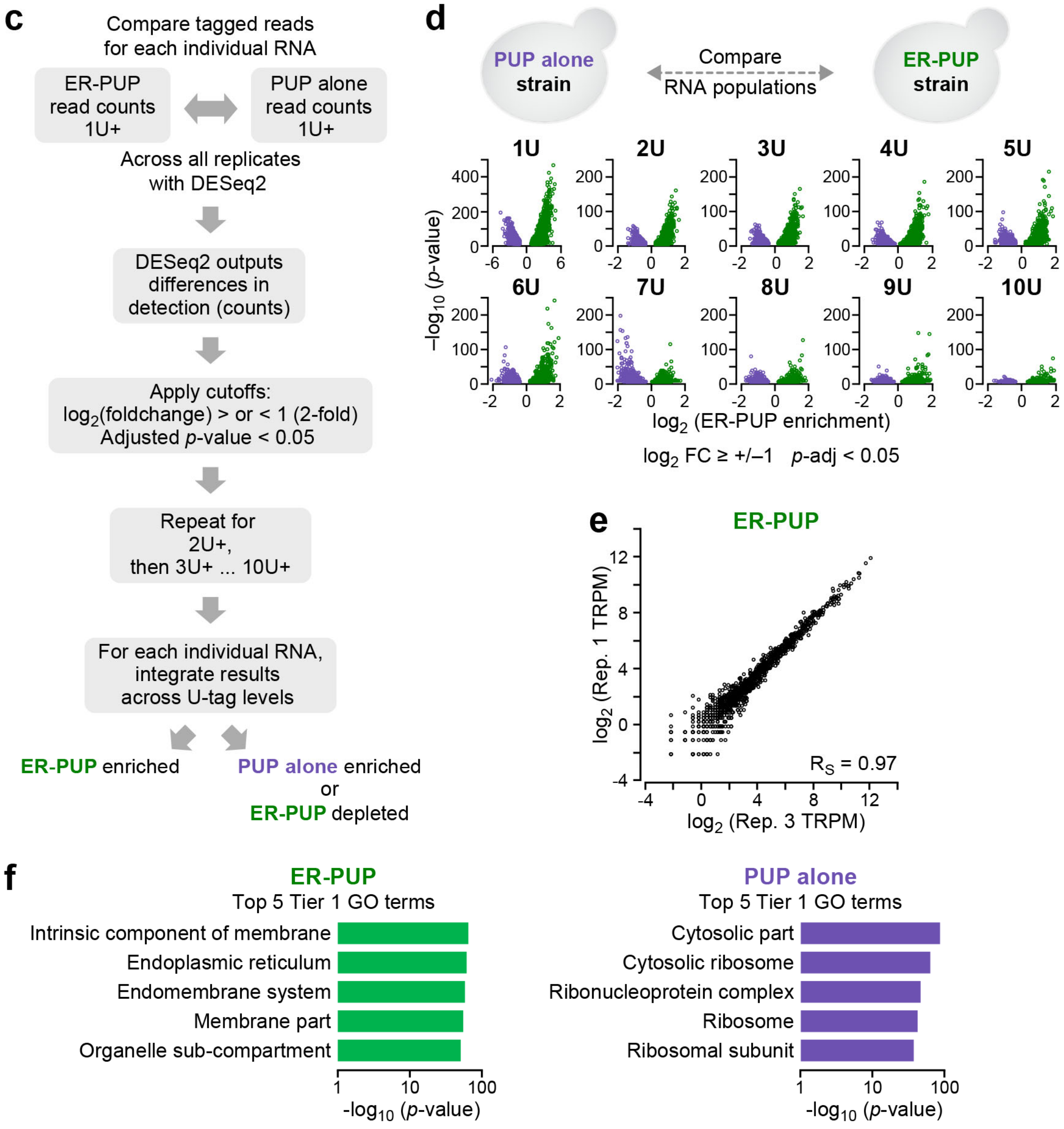


**Figure S3. ER-enriched tagging events.**

**a)** *Statistics.* ER-PUP sequencing statistics across three biological replicates. **b)** *Reproducibility.* Data as in (a). **c)** *Flowchart of computational analysis.* The steps used to identify RNAs whose tagging was enriched with ER-PUP or PUP alone are depicted. DESeq2[35] was used to identify statistically significant differences (adjusted *p*-value < 0.05), ($\log_2(\Delta$ tagged reads$) \geq 1$)). **d)** Enrichment of individual RNAs vs number of U's added. Differences (x-axis) and significance (y-axis) values distinguish individual RNA species (dots). Each U-tag length (1U-10U) was analyzed separately. RNAs to the right of zero on the x-axis are enriched by ER-PUP (green dots), while the ones on the left are depleted from ER PUP (aka enriched by PUP alone, purple dots). **e)** *Reproducibility.* Comparison of data from two biological ER-PUP replicates. **f)** *Functional enrichments.* Top five ER-PUP (top) and PUP alone (bottom) gene ontology (GO) terms are depicted.

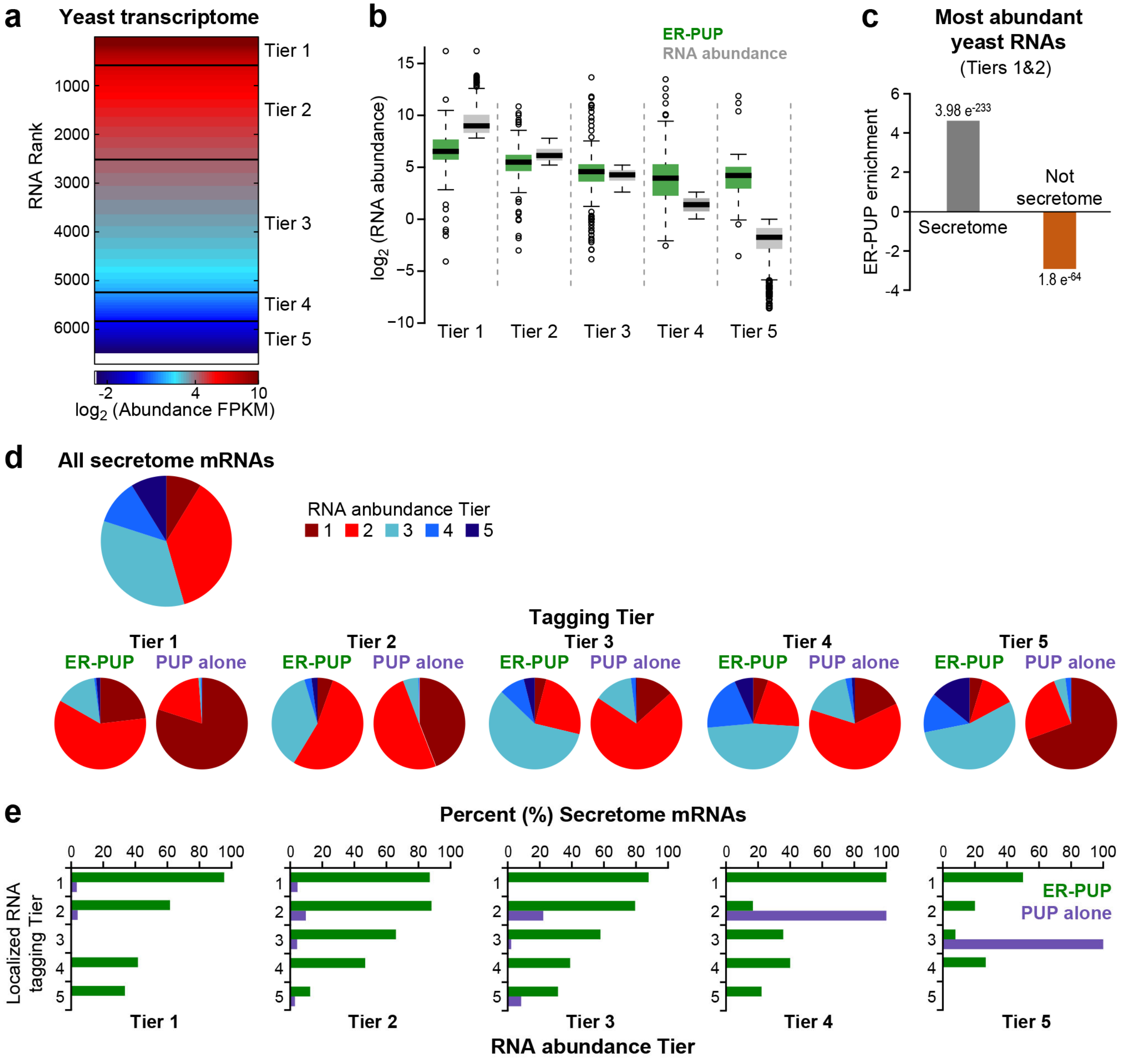


**Figure S4. DESeq2 analysis of tagging and abundance.**

**a)** The yeast transcriptome comprises five RNA abundance[26] (FPKM) tiers (clusters, K-means), ranked from most (Tier 1) to least abundant (Tier 5). **b)** Per tier RNA abundance of ER-enriched RNAs (green) and of all yeast transcripts (grey). **c)** Relationship of ER-PUP enrichment and secretome mRNAs in abundance Tiers 1 & 2. Hypergeometric distribution significance ($p$-values) are reported **d)** Per Tier ER-PUP and PUP alone abundance composition. **e)** Tagged RNAs populate five distinct RNA abundance bins (bar graphs). For each, the ER-PUP (green) and PUP alone Tiers (purple) (y-axes) project the proportion (%, x-axes) of secretome mRNAs.

Medina-Muñoz et al. 2019 Figure S5

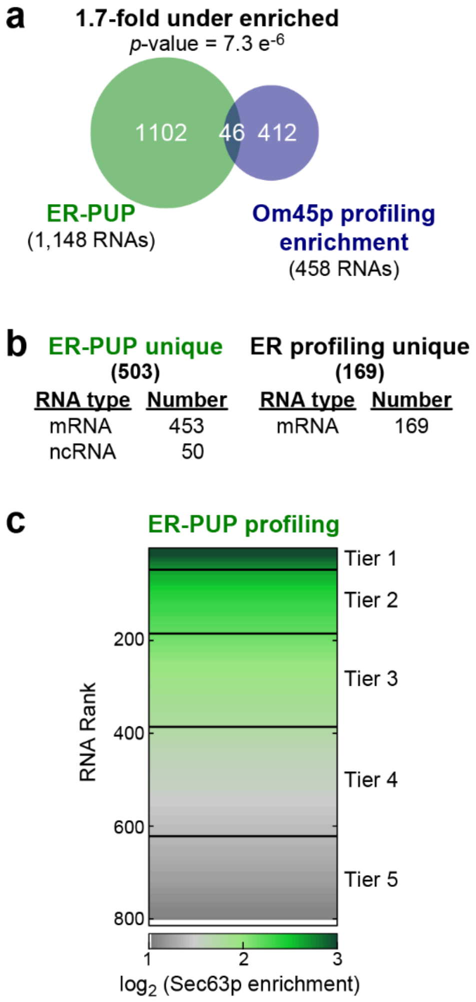


**Figure S5. ER tagging and ribosome association.**

**a)** Comparison of ER-tagged RNAs with mRNAs identified by mitochondrial (Om45) profiling.[22] **b)** RNAs identified uniquely by either tagging or profiling**. c)** ER (Sec63p) profiling data[21] comprises five enrichment clusters (tiers from K-means), from most (Tier 1) to least ribosome association (Tier 5).

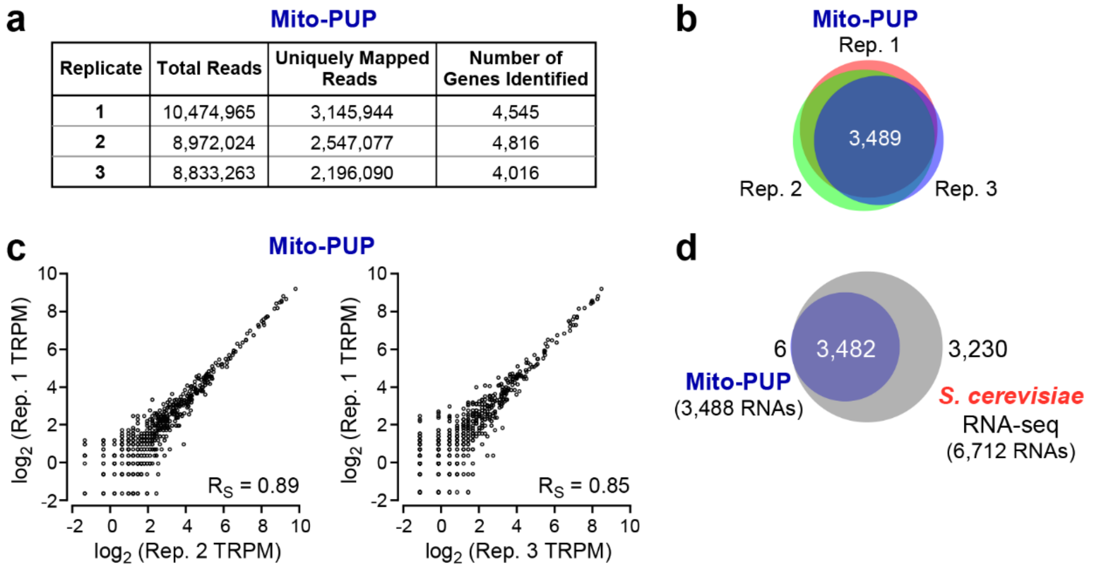

a — Mito-PUP

| Replicate | Total Reads | Uniquely Mapped Reads | Number of Genes Identified |
|---|---|---|---|
| 1 | 10,474,965 | 3,145,944 | 4,545 |
| 2 | 8,972,024 | 2,547,077 | 4,816 |
| 3 | 8,833,263 | 2,196,090 | 4,016 |



**Figure S6. Mito-PUP tagging statistics and reproducibility.**

**a)** *Statistics*. Data from three biological replicates. **b)** RNA species detected reproducibly across three replicates. **c)** Reproducibility of Mito-PUP tagging data. Relative abundance of Mito-PUP tagged RNAs (expressed as TRPM) (black dots) in pairs of biological replicates. **d)** Mito-PUP tagged RNAs vs the yeast transcriptome.[26]

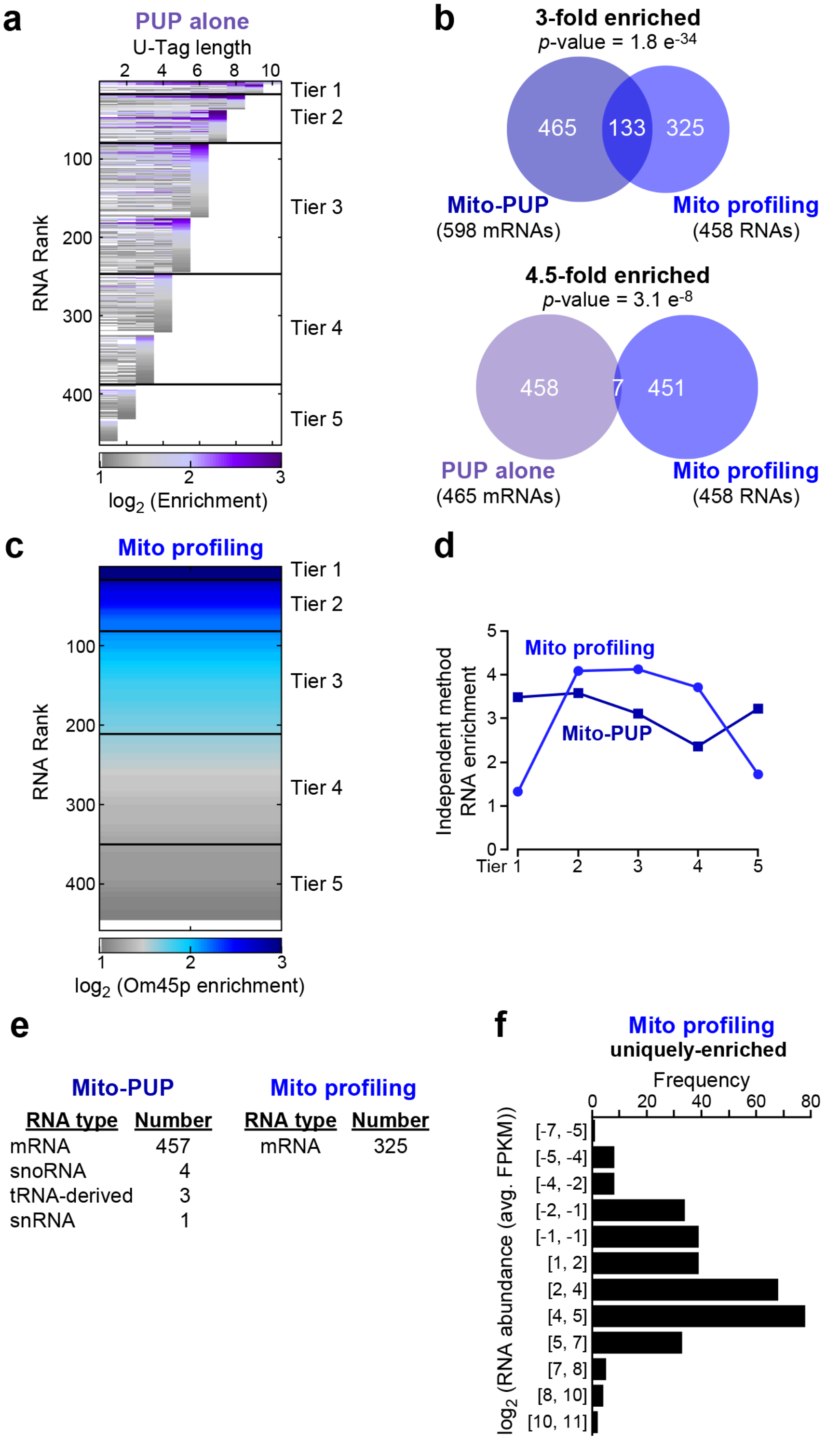


**Figure S7. Mito-PUP tagging.**

**a)** PUP alone-enriched RNAs (relative to Mito-PUP) clustered and ranked. Higher U-tag length yields a higher rank, from Tier 1 (longest) to Tier 5 (shortest). **b)** RNAs tagged by Mito-PUP (top) or PUP alone (bottom) vs mitochondria-proximal ribosome associated mRNAs (obtained using Om45 as an anchor[22]). **c)** Mitochondrial ribosome profiling data[22] used to define five (K-means) RNA clusters (Tiers) from the highest (Tier 1) to lowest (Tier 5) association with ribosomes. **d)** RNAs detected by both mitochondrial tagging and profiling. For RNAs detected by both tagging and profiling, the rank in each tier of the two individual methods is depicted. **e)** RNAs unique to tagging and profiling. **f)** Distribution of mRNAs unique to mitochondrial profiling across a series of RNA abundance[26] bins. Each bin (y-axis) represents a range of RNA abundance limits ($log_2$(FPKM)). The number of RNAs that fall within those limits is projected by a black bar along the x-axis.

Medina-Muñoz et al. 2019 Figure S8

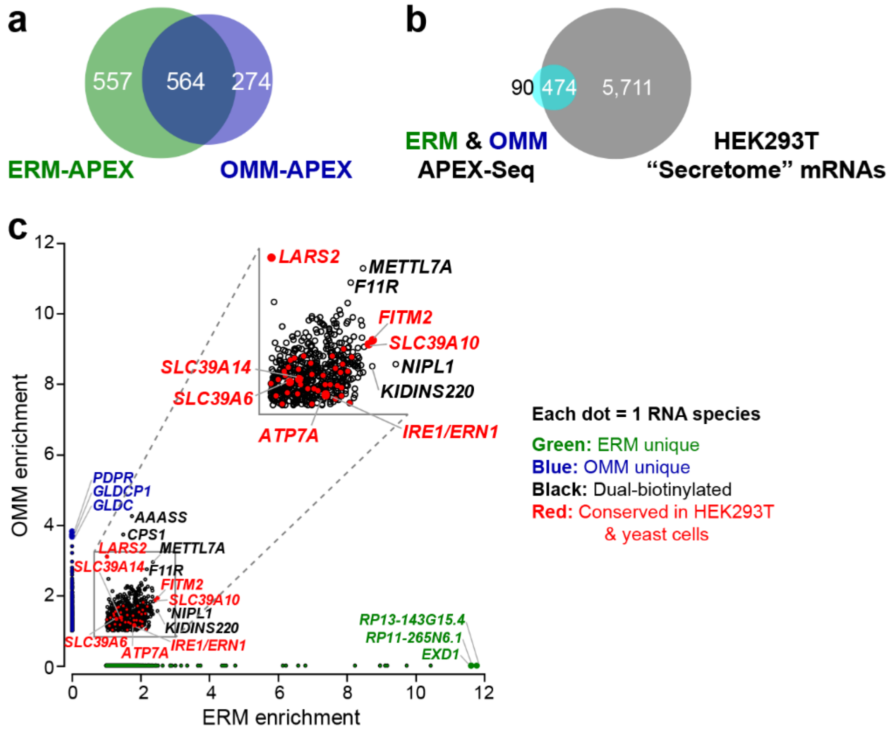


**Figure S8. Conservation of dual-tagged RNA species in yeast and human cultured cells.**

**a**) RNAs detected by APEX-seq in human (HEK293T) cells at ER membrane (ERM) vs outer mitochondrial membrane (OMM) (data reprocessed from *ref 25*). **b)** Dual-labeled secretome mRNAs from APEX-Seq. **c)** RNAs identified in tagging vs APEX-seq: conservation of RNAs at both ER and mitochondria. Each mRNA is represented by a dot, and plotted vs enrichment in ERM (x-axis) and OMM (y-axis). Organelle-specific mRNAs are green (ER) or blue (OMM), and lie on the x- or y-axes, respectively. Dual localized mRNAs in HEK293T cells are black. Red indicates RNAs that are identified at both the ER and mitochondria in both yeast and HEK293T cells. Blow-up insert highlights dual-tagged RNAs.

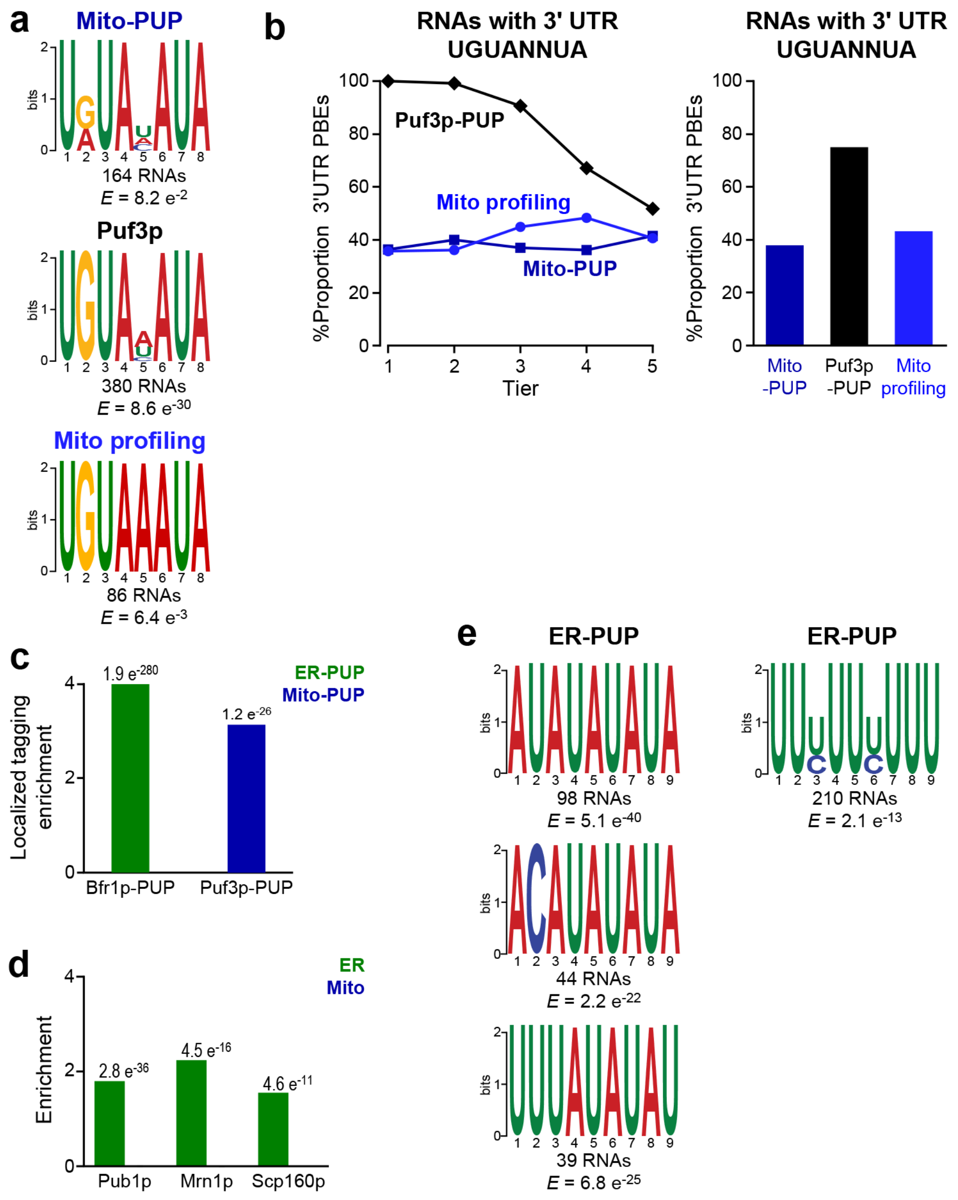


**Figure S9. Sequence elements correlated with ER or mitochondrial localization.**

**a)** Enrichment of motifs associated with mRNAs tagged by Mito-PUP (top), physically associated with Puf3p[26] (center), or bound by mitochondria-proximal ribosomes inferred from profiling[22] (bottom). **b)** fraction of mRNAs that with one or more Puf3p-binding element (PBE) in their 3' UTR, as a function of their tier in Mito-PUP tagging (dark blue), Puf3p-tagging[26] (black) or mitochondrial profiling[22] (light blue) tier Bar graph shows percent (%) of all RNAs detected that contain at least one PBE in the 3' UTR. **c)** RNAs that physically interact with Bfr1p and Puf3p targets[26] from among ER-PUP- (green) and Mito-PUP-tagged (blue) RNAs. Hypergeometric distribution significance (p-values) are shown. **d)** ER-PUP enriches additional RNA-binding protein targets (identified in RIP-chip[62]). **e)** Enrichment of motifs associated with mRNAs tagged by ER-PUP.
