## Supplementary Tables for "Records of RNA localization through covalent tagging"

### Supplementary Table 1. ER-enriched RNAs: GO Terms.

| Biological Process GO Terms | p-Value | Matches |
|---|---|---|
| transmembrane transport [GO:0055085] | 1.1E-44 | 205 |
| ion transport [GO:0006811] | 1.7E-40 | 177 |
| glycosylation [GO:0070085] | 2.0E-32 | 67 |
| ion transmembrane transport [GO:0034220] | 4.2E-32 | 130 |
| glycoprotein metabolic process [GO:0009100] | 3.5E-31 | 66 |
| transport [GO:0006810] | 5.5E-30 | 376 |
| glycoprotein biosynthetic process [GO:0009101] | 1.5E-29 | 62 |
| protein glycosylation [GO:0006486] | 2.5E-29 | 60 |
| macromolecule glycosylation [GO:0043413] | 2.5E-29 | 60 |
| establishment of localization [GO:0051234] | 6.3E-28 | 380 |
| localization [GO:0051179] | 4.0E-27 | 416 |
| mannosylation [GO:0097502] | 1.1E-26 | 43 |
| lipid metabolic process [GO:0006629] | 9.8E-25 | 142 |
| cation transport [GO:0006812] | 1.4E-23 | 106 |
| membrane lipid metabolic process [GO:0006643] | 2.1E-23 | 58 |
| anion transport [GO:0006820] | 1.9E-22 | 96 |
| membrane lipid biosynthetic process [GO:0046467] | 1.9E-21 | 50 |
| cellular lipid metabolic process [GO:0044255] | 2.5E-21 | 130 |
| metal ion transport [GO:0030001] | 1.7E-20 | 56 |
| protein N-linked glycosylation [GO:0006487] | 6.0E-19 | 39 |
| lipid biosynthetic process [GO:0008610] | 7.2E-19 | 96 |
| cation transmembrane transport [GO:0098655] | 1.4E-18 | 88 |
| metal ion homeostasis [GO:0055065] | 1.1E-17 | 65 |
| cation homeostasis [GO:0055080] | 1.7E-16 | 76 |
| anion transmembrane transport [GO:0098656] | 2.3E-16 | 59 |
| cellular metal ion homeostasis [GO:0006875] | 8.1E-16 | 59 |
| inorganic ion homeostasis [GO:0098771] | 2.1E-15 | 72 |

| | | |
|---|---|---|
| carbohydrate derivative biosynthetic process [GO:1901137] | 5.9E-15 | 104 |
| organic anion transport [GO:0015711] | 6.0E-15 | 74 |
| glycolipid metabolic process [GO:0006664] | 1.3E-14 | 31 |
| liposaccharide metabolic process [GO:1903509] | 1.3E-14 | 31 |
| glycolipid biosynthetic process [GO:0009247] | 1.5E-14 | 29 |
| transition metal ion transport [GO:0000041] | 3.3E-14 | 36 |
| cellular cation homeostasis [GO:0030003] | 3.5E-14 | 69 |
| ion homeostasis [GO:0050801] | 8.1E-14 | 76 |
| chemical homeostasis [GO:0048878] | 1.3E-13 | 87 |
| protein lipidation [GO:0006497] | 1.5E-13 | 35 |
| lipoprotein biosynthetic process [GO:0042158] | 1.5E-13 | 35 |
| cell wall organization or biogenesis [GO:0071554] | 2.4E-13 | 108 |
| GPI anchor metabolic process [GO:0006505] | 3.7E-13 | 28 |
| lipoprotein metabolic process [GO:0042157] | 4.1E-13 | 35 |
| inorganic ion transmembrane transport [GO:0098660] | 1.1E-12 | 73 |
| GPI anchor biosynthetic process [GO:0006506] | 2.1E-12 | 26 |
| transition metal ion homeostasis [GO:0055076] | 2.4E-12 | 49 |
| protein O-linked glycosylation [GO:0006493] | 4.4E-12 | 21 |
| cellular homeostasis [GO:0019725] | 5.3E-12 | 88 |
| cellular chemical homeostasis [GO:0055082] | 5.4E-12 | 76 |
| cellular ion homeostasis [GO:0006873] | 5.8E-12 | 69 |
| cellular transition metal ion homeostasis [GO:0046916] | 1.5E-11 | 45 |
| inorganic cation transmembrane transport [GO:0098662] | 4.5E-11 | 65 |
| divalent inorganic cation transport [GO:0072511] | 6.5E-11 | 28 |
| divalent metal ion transport [GO:0070838] | 9.4E-11 | 27 |
| organic acid transport [GO:0015849] | 1.5E-10 | 46 |
| phospholipid metabolic process [GO:0006644] | 1.9E-10 | 67 |
| carboxylic acid transport [GO:0046942] | 4.7E-10 | 45 |
| sphingolipid metabolic process [GO:0006665] | 1.0E-09 | 32 |
| cell wall macromolecule metabolic process [GO:0044036] | 1.0E-09 | 32 |
| glycerolipid metabolic process [GO:0046486] | 1.1E-09 | 59 |
| organic substance transport [GO:0071702] | 1.8E-09 | 237 |
| amino acid transmembrane transport [GO:0003333] | 2.2E-09 | 29 |

| | | |
|---|---|---|
| carbohydrate derivative metabolic process [GO:1901135] | 4.5E-09 | 120 |
| amino acid transport [GO:0006865] | 5.4E-09 | 33 |
| glycerophospholipid metabolic process [GO:0006650] | 7.1E-09 | 54 |
| glycerolipid biosynthetic process [GO:0045017] | 8.6E-09 | 42 |
| phospholipid biosynthetic process [GO:0008654] | 8.6E-09 | 51 |
| divalent inorganic cation homeostasis [GO:0072507] | 8.7E-09 | 24 |
| organic acid transmembrane transport [GO:1903825] | 2.2E-08 | 35 |
| cellular divalent inorganic cation homeostasis [GO:0072503] | 3.9E-08 | 23 |
| glycerophospholipid biosynthetic process [GO:0046474] | 4.9E-08 | 40 |
| carboxylic acid transmembrane transport [GO:1905039] | 7.5E-08 | 34 |
| phosphatidylinositol biosynthetic process [GO:0006661] | 9.8E-08 | 28 |
| sphingolipid biosynthetic process [GO:0030148] | 1.0E-07 | 25 |
| phosphatidylinositol metabolic process [GO:0046488] | 1.0E-07 | 38 |
| external encapsulating structure organization [GO:0045229] | 3.5E-07 | 83 |
| cell wall organization [GO:0071555] | 3.5E-07 | 83 |
| cell wall macromolecule biosynthetic process [GO:0044038] | 3.6E-07 | 27 |
| cellular component macromolecule biosynthetic process [GO:0070589] | 3.6E-07 | 27 |
| homeostatic process [GO:0042592] | 6.2E-07 | 103 |
| cell wall biogenesis [GO:0042546] | 6.9E-07 | 47 |
| nitrogen compound transport [GO:0071705] | 1.0E-06 | 206 |
| fungal-type cell wall organization or biogenesis [GO:0071852] | 3.1E-06 | 76 |
| response to endoplasmic reticulum stress [GO:0034976] | 1.5E-05 | 38 |
| maintenance of protein localization in endoplasmic reticulum [GO:0035437] | 1.5E-05 | 11 |
| drug transport [GO:0015893] | 1.8E-05 | 37 |
| cell wall mannoprotein biosynthetic process [GO:0000032] | 2.3E-05 | 15 |
| mannoprotein metabolic process [GO:0006056] | 2.3E-05 | 15 |
| mannoprotein biosynthetic process [GO:0006057] | 2.3E-05 | 15 |
| cell wall glycoprotein biosynthetic process [GO:0031506] | 2.3E-05 | 15 |
| iron ion transport [GO:0006826] | 3.3E-05 | 17 |
| protein retention in ER lumen [GO:0006621] | 9.1E-05 | 10 |
| polysaccharide metabolic process [GO:0005976] | 1.1E-04 | 39 |
| aromatic amino acid transport [GO:0015801] | 1.5E-04 | 11 |
| protein mannosylation [GO:0035268] | 1.5E-04 | 11 |

| | | |
|---|---|---|
| drug transmembrane transport [GO:0006855] | 1.6E-04 | 26 |
| oligosaccharide-lipid intermediate biosynthetic process [GO:0006490] | 1.6E-04 | 12 |
| regulation of biological quality [GO:0065008] | 2.0E-04 | 145 |
| protein localization to endoplasmic reticulum [GO:0070972] | 3.8E-04 | 27 |
| ERAD pathway [GO:0036503] | 4.4E-04 | 26 |
| N-glycan processing [GO:0006491] | 5.5E-04 | 9 |
| calcium ion transport [GO:0006816] | 8.5E-04 | 10 |
| beta-glucan metabolic process [GO:0051273] | 8.5E-04 | 16 |
| beta-glucan biosynthetic process [GO:0051274] | 8.5E-04 | 16 |
| calcium ion transmembrane transport [GO:0070588] | 3.3E-03 | 8 |
| cofactor transport [GO:0051181] | 4.3E-03 | 18 |
| zinc ion transport [GO:0006829] | 4.3E-03 | 10 |
| nucleobase transport [GO:0015851] | 4.6E-03 | 9 |
| cellular manganese ion homeostasis [GO:0030026] | 4.6E-03 | 9 |
| protein O-linked mannosylation [GO:0035269] | 4.6E-03 | 9 |
| manganese ion homeostasis [GO:0055071] | 4.6E-03 | 9 |
| fungal-type cell wall organization [GO:0031505] | 4.9E-03 | 61 |
| regulation of membrane lipid distribution [GO:0097035] | 5.8E-03 | 14 |
| basic amino acid transport [GO:0015802] | 6.8E-03 | 12 |
| lipid translocation [GO:0034204] | 1.0E-02 | 13 |
| cellular potassium ion transport [GO:0071804] | 1.1E-02 | 11 |
| potassium ion transmembrane transport [GO:0071805] | 1.1E-02 | 11 |
| inorganic anion transport [GO:0015698] | 1.3E-02 | 18 |
| cellular zinc ion homeostasis [GO:0006882] | 1.6E-02 | 10 |
| zinc ion homeostasis [GO:0055069] | 1.6E-02 | 10 |
| basic amino acid transmembrane transport [GO:1990822] | 1.6E-02 | 10 |
| polysaccharide biosynthetic process [GO:0000271] | 1.8E-02 | 26 |
| tyrosine transport [GO:0015828] | 2.0E-02 | 7 |
| ceramide metabolic process [GO:0006672] | 2.2E-02 | 9 |
| copper ion import [GO:0015677] | 2.2E-02 | 9 |
| zinc ion transmembrane transport [GO:0071577] | 2.2E-02 | 9 |
| iron ion homeostasis [GO:0055072] | 2.2E-02 | 25 |
| manganese ion transport [GO:0006828] | 2.5E-02 | 8 |

| | | |
|---|---|---|
| fungal-type cell wall beta-glucan metabolic process [GO:0070879] | 2.5E-02 | 8 |
| fungal-type cell wall beta-glucan biosynthetic process [GO:0070880] | 2.5E-02 | 8 |
| response to unfolded protein [GO:0006986] | 2.9E-02 | 22 |
| calcium ion homeostasis [GO:0055074] | 2.9E-02 | 11 |
| maintenance of protein localization in organelle [GO:0072595] | 2.9E-02 | 11 |
| phospholipid translocation [GO:0045332] | 4.0E-02 | 12 |
| cellular polysaccharide biosynthetic process [GO:0033692] | 4.5E-02 | 25 |
| response to organonitrogen compound [GO:0010243] | 4.6E-02 | 31 |
| coenzyme transport [GO:0051182] | 4.7E-02 | 10 |

| Molecular Function GO Terms | p-Value | Matches |
|---|---|---|
| transmembrane transporter activity [GO:0022857] | 5.1E-54 | 185 |
| transporter activity [GO:0005215] | 7.5E-53 | 192 |
| ion transmembrane transporter activity [GO:0015075] | 5.7E-39 | 136 |
| inorganic molecular entity transmembrane transporter activity [GO:0015318] | 5.1E-31 | 119 |
| transferase activity, transferring hexosyl groups [GO:0016758] | 2.7E-30 | 63 |
| mannosyltransferase activity [GO:0000030] | 9.4E-27 | 41 |
| transferase activity, transferring glycosyl groups [GO:0016757] | 2.5E-26 | 68 |
| cation transmembrane transporter activity [GO:0008324] | 1.0E-24 | 94 |
| metal ion transmembrane transporter activity [GO:0046873] | 1.4E-20 | 44 |
| anion transmembrane transporter activity [GO:0008509] | 2.8E-18 | 60 |
| active transmembrane transporter activity [GO:0022804] | 4.7E-17 | 77 |
| organic anion transmembrane transporter activity [GO:0008514] | 4.6E-15 | 50 |
| inorganic cation transmembrane transporter activity [GO:0022890] | 7.3E-15 | 69 |
| secondary active transmembrane transporter activity [GO:0015291] | 3.0E-14 | 50 |
| transition metal ion transmembrane transporter activity [GO:0046915] | 6.0E-13 | 27 |
| organic acid transmembrane transporter activity [GO:0005342] | 2.6E-11 | 39 |
| carboxylic acid transmembrane transporter activity [GO:0046943] | 9.6E-11 | 38 |
| drug transmembrane transporter activity [GO:0015238] | 1.9E-10 | 35 |
| amino acid transmembrane transporter activity [GO:0015171] | 1.6E-08 | 27 |
| alpha-1,2-mannosyltransferase activity [GO:0000026] | 8.2E-08 | 13 |
| signaling receptor activity [GO:0038023] | 1.6E-07 | 14 |
| hydrolase activity, hydrolyzing O-glycosyl compounds [GO:0004553] | 3.4E-06 | 26 |
| hydrolase activity, acting on glycosyl bonds [GO:0016798] | 1.7E-05 | 29 |
| symporter activity [GO:0015293] | 2.3E-05 | 24 |
| antiporter activity [GO:0015297] | 1.0E-04 | 19 |
| alpha-1,6-mannosyltransferase activity [GO:0000009] | 1.4E-04 | 9 |
| aromatic amino acid transmembrane transporter activity [GO:0015173] | 1.4E-04 | 9 |
| transmembrane signaling receptor activity [GO:0004888] | 2.1E-04 | 10 |

| | | |
|---|---|---|
| nucleobase-containing compound transmembrane transporter activity [GO:0015932] | 3.4E-04 | 16 |
| basic amino acid transmembrane transporter activity [GO:0015174] | 7.8E-04 | 11 |
| cofactor transmembrane transporter activity [GO:0051184] | 8.8E-04 | 13 |
| calcium ion transmembrane transporter activity [GO:0015085] | 9.1E-04 | 8 |
| nucleobase transmembrane transporter activity [GO:0015205] | 1.2E-03 | 9 |
| ATPase activity, coupled to movement of substances [GO:0043492] | 2.6E-03 | 28 |
| oxidoreductase activity, oxidizing metal ions [GO:0016722] | 4.0E-03 | 10 |
| xenobiotic transmembrane transporter activity [GO:0042910] | 4.0E-03 | 10 |
| solute:cation symporter activity [GO:0015294] | 5.3E-03 | 19 |
| zinc ion transmembrane transporter activity [GO:0005385] | 5.7E-03 | 9 |
| dolichyl-phosphate-mannose-protein mannosyltransferase activity [GO:0004169] | 5.8E-03 | 7 |
| monovalent inorganic cation transmembrane transporter activity [GO:0015077] | 9.0E-03 | 36 |
| carbohydrate derivative transmembrane transporter activity [GO:1901505] | 1.0E-02 | 14 |
| potassium ion transmembrane transporter activity [GO:0015079] | 1.2E-02 | 10 |
| ammonium transmembrane transporter activity [GO:0008519] | 1.7E-02 | 11 |
| ATPase activity, coupled to transmembrane movement of substances [GO:0042626] | 1.7E-02 | 26 |
| divalent inorganic cation transmembrane transporter activity [GO:0072509] | 2.0E-02 | 9 |
| alpha-1,3-mannosyltransferase activity [GO:0000033] | 3.7E-02 | 6 |
| L-tyrosine transmembrane transporter activity [GO:0005302] | 3.7E-02 | 6 |
| manganese ion transmembrane transporter activity [GO:0005384] | 4.0E-02 | 7 |
| protein-cysteine S-palmitoyltransferase activity [GO:0019706] | 4.0E-02 | 7 |
| protein-cysteine S-acyltransferase activity [GO:0019707] | 4.0E-02 | 7 |
| vitamin transmembrane transporter activity [GO:0090482] | 4.0E-02 | 7 |
| exopeptidase activity [GO:0008238] | 4.8E-02 | 15 |

| Pathways | p-Value | Matches |
|---|---|---|
| sphingolipid metabolism | 1.3E-08 | 15 |
| lipid-linked oligosaccharide biosynthesis | 2.9E-07 | 11 |
| triglyceride biosynthesis | 3.6E-02 | 5 |

**Supplementary Table 2. Mitochondria-enriched RNAs: GO Terms.**

| Biological Process GO Terms | p-Value | Matches |
|---|---|---|
| transmembrane transport [GO:0055085] | 7.8E-17 | 108 |
| metal ion homeostasis [GO:0055065] | 5.6E-16 | 48 |
| metal ion transport [GO:0030001] | 8.5E-16 | 40 |
| ion transport [GO:0006811] | 5.9E-14 | 91 |
| cation homeostasis [GO:0055080] | 7.4E-13 | 52 |
| transition metal ion homeostasis [GO:0055076] | 1.1E-12 | 38 |
| inorganic ion homeostasis [GO:0098771] | 1.7E-12 | 50 |
| chemical homeostasis [GO:0048878] | 2.5E-12 | 61 |
| ion homeostasis [GO:0050801] | 8.7E-12 | 53 |
| cellular metal ion homeostasis [GO:0006875] | 2.4E-11 | 40 |
| transition metal ion transport [GO:0000041] | 1.1E-09 | 25 |
| ion transmembrane transport [GO:0034220] | 2.6E-09 | 64 |
| cellular chemical homeostasis [GO:0055082] | 2.6E-09 | 51 |
| iron ion homeostasis [GO:0055072] | 3.1E-09 | 27 |
| inorganic ion transmembrane transport [GO:0098660] | 4.9E-09 | 48 |
| cellular cation homeostasis [GO:0030003] | 6.2E-09 | 44 |
| cellular transition metal ion homeostasis [GO:0046916] | 1.7E-08 | 31 |
| cellular ion homeostasis [GO:0006873] | 3.0E-08 | 45 |
| inorganic cation transmembrane transport [GO:0098662] | 5.7E-08 | 43 |
| cellular homeostasis [GO:0019725] | 1.6E-07 | 55 |
| cation transport [GO:0006812] | 2.6E-07 | 54 |
| iron ion transport [GO:0006826] | 1.2E-06 | 15 |
| homeostatic process [GO:0042592] | 1.4E-06 | 69 |
| cation transmembrane transport [GO:0098655] | 4.7E-06 | 46 |
| cellular iron ion homeostasis [GO:0006879] | 9.3E-05 | 20 |
| mitochondrion organization [GO:0007005] | 1.4E-04 | 56 |
| transport [GO:0006810] | 1.7E-04 | 180 |
| divalent inorganic cation transport [GO:0072511] | 5.7E-04 | 16 |
| regulation of biological quality [GO:0065008] | 6.2E-04 | 92 |
| establishment of localization [GO:0051234] | 8.3E-04 | 182 |

| | | |
|---|---|---|
| divalent metal ion transport [GO:0070838] | 1.7E-03 | 15 |
| mitochondrial transport [GO:0006839] | 2.2E-03 | 26 |
| cofactor transport [GO:0051181] | 3.3E-03 | 14 |
| divalent inorganic cation homeostasis [GO:0072507] | 3.3E-03 | 14 |
| phosphate ion transport [GO:0006817] | 4.1E-03 | 9 |
| protein localization to mitochondrion [GO:0070585] | 9.2E-03 | 21 |
| establishment of protein localization to mitochondrion [GO:0072655] | 9.2E-03 | 21 |
| anion transport [GO:0006820] | 1.1E-02 | 40 |
| iron ion transmembrane transport [GO:0034755] | 1.2E-02 | 8 |
| mitochondrial genome maintenance [GO:0000002] | 1.2E-02 | 16 |
| mitochondrial transmembrane transport [GO:1990542] | 1.4E-02 | 20 |
| cellular divalent inorganic cation homeostasis [GO:0072503] | 1.5E-02 | 13 |
| iron coordination entity transport [GO:1901678] | 2.0E-02 | 9 |
| localization [GO:0051179] | 3.0E-02 | 195 |

| Cellular Component GO Terms | p-Value | Matches |
|---|---|---|
| mitochondrion [GO:0005739] | 1.1E-23 | 220 |
| mitochondrial part [GO:0044429] | 2.8E-21 | 141 |
| cytoplasmic part [GO:0044444] | 6.2E-21 | 446 |
| intrinsic component of membrane [GO:0031224] | 3.1E-20 | 253 |
| membrane [GO:0016020] | 2.8E-18 | 326 |
| membrane part [GO:0044425] | 1.5E-17 | 279 |
| integral component of membrane [GO:0016021] | 7.8E-15 | 231 |
| mitochondrial membrane [GO:0031966] | 1.2E-14 | 94 |
| organelle membrane [GO:0031090] | 1.2E-14 | 170 |
| mitochondrial envelope [GO:0005740] | 1.4E-14 | 100 |
| organelle inner membrane [GO:0019866] | 3.0E-12 | 69 |
| mitochondrial inner membrane [GO:0005743] | 3.5E-12 | 67 |
| cell periphery [GO:0071944] | 1.5E-11 | 142 |
| organelle envelope [GO:0031967] | 1.1E-10 | 111 |
| envelope [GO:0031975] | 1.1E-10 | 111 |
| cytoplasm [GO:0005737] | 2.5E-10 | 509 |
| intracellular membrane-bounded organelle [GO:0043231] | 3.2E-09 | 486 |
| extracellular region [GO:0005576] | 3.9E-09 | 38 |
| membrane-bounded organelle [GO:0043227] | 5.2E-09 | 490 |
| endoplasmic reticulum [GO:0005783] | 6.5E-09 | 117 |
| intracellular organelle [GO:0043229] | 2.0E-08 | 511 |
| organelle [GO:0043226] | 2.3E-08 | 511 |
| storage vacuole [GO:0000322] | 3.6E-08 | 87 |
| fungal-type vacuole [GO:0000324] | 3.6E-08 | 87 |
| lytic vacuole [GO:0000323] | 4.0E-08 | 87 |
| plasma membrane [GO:0005886] | 8.0E-08 | 100 |
| vacuole [GO:0005773] | 1.1E-06 | 92 |
| cell part [GO:0044464] | 3.2E-06 | 573 |

| | | |
|---|---|---|
| mitochondrial matrix [GO:0005759] | 4.4E-06 | 52 |
| cell wall [GO:0005618] | 6.0E-06 | 36 |
| external encapsulating structure [GO:0030312] | 6.0E-06 | 36 |
| fungal-type cell wall [GO:0009277] | 6.1E-06 | 35 |
| cell [GO:0005623] | 6.6E-06 | 573 |
| endomembrane system [GO:0012505] | 1.6E-05 | 153 |
| endoplasmic reticulum part [GO:0044432] | 2.4E-05 | 73 |
| endoplasmic reticulum subcompartment [GO:0098827] | 4.3E-05 | 70 |
| mitochondrial membrane part [GO:0044455] | 1.0E-04 | 42 |
| endoplasmic reticulum membrane [GO:0005789] | 1.9E-04 | 67 |
| anchored component of membrane [GO:0031225] | 2.6E-04 | 21 |
| intrinsic component of mitochondrial membrane [GO:0098573] | 3.0E-04 | 22 |
| intracellular organelle part [GO:0044446] | 3.2E-04 | 351 |
| organelle subcompartment [GO:0031984] | 3.8E-04 | 84 |
| organelle part [GO:0044422] | 4.7E-04 | 351 |
| nuclear outer membrane-endoplasmic reticulum membrane network [GO:0042175] | 6.9E-04 | 67 |
| integral component of mitochondrial membrane [GO:0032592] | 2.0E-03 | 20 |
| intrinsic component of organelle membrane [GO:0031300] | 2.1E-03 | 31 |
| bounding membrane of organelle [GO:0098588] | 2.5E-03 | 92 |
| mitochondrial protein complex [GO:0098798] | 3.1E-03 | 42 |
| integral component of organelle membrane [GO:0031301] | 7.0E-03 | 29 |
| intrinsic component of mitochondrial inner membrane [GO:0031304] | 7.7E-03 | 14 |
| plasma membrane part [GO:0044459] | 9.1E-03 | 35 |
| intrinsic component of plasma membrane [GO:0031226] | 1.3E-02 | 28 |
| whole membrane [GO:0098805] | 1.4E-02 | 82 |
| vacuolar membrane [GO:0005774] | 2.4E-02 | 48 |
| integral component of plasma membrane [GO:0005887] | 3.6E-02 | 26 |

| Molecular Function GO Terms | p-Value | Matches |
|---|---|---|
| transmembrane transporter activity [GO:0022857] | 7.3E-15 | 92 |
| transporter activity [GO:0005215] | 1.5E-13 | 94 |
| metal ion transmembrane transporter activity [GO:0046873] | 2.1E-11 | 29 |
| inorganic molecular entity transmembrane transporter activity [GO:0015318] | 2.5E-07 | 58 |
| ion transmembrane transporter activity [GO:0015075] | 4.1E-07 | 61 |
| transition metal ion transmembrane transporter activity [GO:0046915] | 6.0E-07 | 18 |
| inorganic cation transmembrane transporter activity [GO:0022890] | 2.8E-05 | 39 |
| active transmembrane transporter activity [GO:0022804] | 2.0E-04 | 40 |
| cation transmembrane transporter activity [GO:0008324] | 2.8E-04 | 43 |
| secondary active transmembrane transporter activity [GO:0015291] | 4.0E-03 | 25 |
| peptide transmembrane transporter activity [GO:1904680] | 5.0E-03 | 14 |
| inorganic phosphate transmembrane transporter activity [GO:0005315] | 1.4E-02 | 6 |
| amide transmembrane transporter activity [GO:0042887] | 4.1E-02 | 15 |
| iron ion transmembrane transporter activity [GO:0005381] | 4.8E-02 | 7 |

**Supplementary Table 3. 133 mRNAs common to Mitochondria tagging and Mitochondrial Profiling: GO Terms.**

| Biological Process GO Terms | p-Value | Matches |
|---|---|---|
| mitochondrion organization [GO:0007005] | 2.3E-20 | 41 |
| mitochondrial transport [GO:0006839] | 3.1E-11 | 20 |
| protein localization to mitochondrion [GO:0070585] | 5.3E-10 | 17 |
| establishment of protein localization to mitochondrion [GO:0072655] | 5.3E-10 | 17 |
| mitochondrial transmembrane transport [GO:1990542] | 3.2E-09 | 16 |
| mitochondrial gene expression [GO:0140053] | 3.9E-08 | 24 |
| mitochondrial membrane organization [GO:0007006] | 3.9E-07 | 13 |
| protein targeting to mitochondrion [GO:0006626] | 3.8E-06 | 13 |
| mitochondrial RNA metabolic process [GO:0000959] | 1.0E-05 | 11 |
| mitochondrial genome maintenance [GO:0000002] | 1.3E-05 | 11 |
| protein transmembrane transport [GO:0071806] | 1.8E-05 | 14 |
| transmembrane transport [GO:0055085] | 4.7E-05 | 31 |
| iron ion homeostasis [GO:0055072] | 2.6E-04 | 11 |
| protein transmembrane import into intracellular organelle [GO:0044743] | 5.3E-04 | 11 |
| inner mitochondrial membrane organization [GO:0007007] | 6.6E-04 | 8 |
| transition metal ion homeostasis [GO:0055076] | 2.8E-03 | 12 |
| mitochondrial RNA processing [GO:0000963] | 5.1E-03 | 7 |
| mitochondrial translation [GO:0032543] | 5.9E-03 | 16 |
| intracellular protein transmembrane transport [GO:0065002] | 6.3E-03 | 11 |
| cellular iron ion homeostasis [GO:0006879] | 7.7E-03 | 9 |
| protein import into mitochondrial matrix [GO:0030150] | 3.9E-02 | 6 |
| metal ion homeostasis [GO:0055065] | 5.0E-02 | 12 |

| Cellular Component GO Terms | p-Value | Matches |
|---|---|---|
| mitochondrion [GO:0005739] | 8.4E-71 | 120 |
| mitochondrial part [GO:0044429] | 3.4E-55 | 89 |
| mitochondrial envelope [GO:0005740] | 1.5E-43 | 70 |
| mitochondrial membrane [GO:0031966] | 7.7E-43 | 67 |
| organelle envelope [GO:0031967] | 4.1E-35 | 70 |
| envelope [GO:0031975] | 4.1E-35 | 70 |
| mitochondrial inner membrane [GO:0005743] | 1.5E-34 | 51 |
| organelle inner membrane [GO:0019866] | 1.5E-33 | 51 |
| cytoplasmic part [GO:0044444] | 1.1E-22 | 126 |
| organelle membrane [GO:0031090] | 3.1E-19 | 68 |
| mitochondrial membrane part [GO:0044455] | 1.0E-15 | 30 |
| intrinsic component of mitochondrial membrane [GO:0098573] | 2.8E-12 | 18 |
| intracellular membrane-bounded organelle [GO:0043231] | 3.8E-11 | 126 |
| cytoplasm [GO:0005737] | 1.8E-10 | 128 |
| membrane-bounded organelle [GO:0043227] | 1.8E-10 | 126 |
| integral component of mitochondrial membrane [GO:0032592] | 3.0E-10 | 16 |
| intracellular organelle [GO:0043229] | 2.1E-09 | 128 |
| organelle [GO:0043226] | 2.2E-09 | 128 |
| intrinsic component of mitochondrial inner membrane [GO:0031304] | 1.4E-08 | 12 |
| mitochondrial matrix [GO:0005759] | 1.0E-07 | 24 |
| intrinsic component of organelle membrane [GO:0031300] | 3.0E-07 | 18 |
| integral component of mitochondrial inner membrane [GO:0031305] | 1.8E-06 | 10 |
| integral component of organelle membrane [GO:0031301] | 9.4E-06 | 16 |
| mitochondrial protein complex [GO:0098798] | 1.8E-05 | 20 |
| mitochondrial outer membrane [GO:0005741] | 3.5E-05 | 14 |
| organelle outer membrane [GO:0031968] | 7.1E-05 | 14 |
| outer membrane [GO:0019867] | 7.9E-05 | 14 |
| intracellular organelle part [GO:0044446] | 9.2E-05 | 95 |
| organelle part [GO:0044422] | 1.1E-04 | 95 |
| membrane [GO:0016020] | 9.8E-04 | 76 |

| | | |
|---|---|---|
| intracellular part [GO:0044424] | 3.5E-03 | 130 |
| intracellular [GO:0005622] | 3.8E-03 | 130 |
| intrinsic component of membrane [GO:0031224] | 2.0E-02 | 55 |
| membrane part [GO:0044425] | 2.6E-02 | 62 |
| i-AAA complex [GO:0031942] | 3.7E-02 | 3 |

| Molecular Function GO Terms | p-Value | Matches |
| --- | --- | --- |
| peptide transmembrane transporter activity [GO:1904680] | 6.6E-06 | 10 |
| amide transmembrane transporter activity [GO:0042887] | 1.2E-04 | 10 |
| protein transmembrane transporter activity [GO:0008320] | 2.0E-04 | 8 |
| macromolecule transmembrane transporter activity [GO:0022884] | 2.0E-04 | 8 |
| protein transporter activity [GO:0008565] | 3.8E-04 | 8 |
| small molecule binding [GO:0036094] | 8.2E-03 | 42 |
| anion binding [GO:0043168] | 3.6E-02 | 41 |

**Supplementary Table 4. Mitochondria tagging Unique vs Mitochondrial Profiling: GO Terms.**

| Biological Process GO Terms | p-Value | Matches |
|---|---|---|
| metal ion transport [GO:0030001] | 5.0E-12 | 32 |
| ion transport [GO:0006811] | 7.3E-11 | 72 |
| metal ion homeostasis [GO:0055065] | 2.1E-10 | 36 |
| transmembrane transport [GO:0055085] | 5.2E-09 | 77 |
| cation homeostasis [GO:0055080] | 6.4E-09 | 40 |
| ion transmembrane transport [GO:0034220] | 6.6E-09 | 54 |
| inorganic ion homeostasis [GO:0098771] | 2.4E-08 | 38 |
| chemical homeostasis [GO:0048878] | 5.1E-08 | 46 |
| ion homeostasis [GO:0050801] | 1.2E-07 | 40 |
| cellular metal ion homeostasis [GO:0006875] | 3.4E-07 | 30 |
| transition metal ion transport [GO:0000041] | 3.9E-07 | 20 |
| inorganic ion transmembrane transport [GO:0098660] | 9.2E-07 | 38 |
| inorganic cation transmembrane transport [GO:0098662] | 1.7E-06 | 35 |
| cellular chemical homeostasis [GO:0055082] | 3.4E-06 | 39 |
| transition metal ion homeostasis [GO:0055076] | 3.4E-06 | 26 |
| cation transport [GO:0006812] | 3.9E-06 | 44 |
| cellular cation homeostasis [GO:0030003] | 4.0E-06 | 34 |
| cellular homeostasis [GO:0019725] | 2.3E-05 | 43 |
| cation transmembrane transport [GO:0098655] | 2.7E-05 | 38 |
| cellular ion homeostasis [GO:0006873] | 3.6E-05 | 34 |
| homeostatic process [GO:0042592] | 2.7E-04 | 53 |
| transport [GO:0006810] | 6.3E-04 | 143 |
| cell wall organization or biogenesis [GO:0071554] | 1.1E-03 | 47 |
| cellular transition metal ion homeostasis [GO:0046916] | 1.3E-03 | 21 |
| iron ion transport [GO:0006826] | 1.6E-03 | 11 |
| external encapsulating structure organization [GO:0045229] | 2.7E-03 | 41 |
| cell wall organization [GO:0071555] | 2.7E-03 | 41 |
| establishment of localization [GO:0051234] | 5.5E-03 | 143 |
| divalent inorganic cation transport [GO:0072511] | 7.5E-03 | 13 |
| anion transport [GO:0006820] | 8.3E-03 | 34 |

| | | |
|---|---|---|
| divalent inorganic cation homeostasis [GO:0072507] | 9.1E-03 | 12 |
| glycerophospholipid metabolic process [GO:0006650] | 1.2E-02 | 25 |
| regulation of biological quality [GO:0065008] | 1.9E-02 | 71 |
| iron ion homeostasis [GO:0055072] | 2.0E-02 | 16 |
| glycerolipid metabolic process [GO:0046486] | 2.5E-02 | 26 |
| divalent metal ion transport [GO:0070838] | 2.6E-02 | 12 |
| anion transmembrane transport [GO:0098656] | 3.0E-02 | 22 |
| localization [GO:0051179] | 4.0E-02 | 155 |
| fungal-type cell wall organization or biogenesis [GO:0071852] | 4.3E-02 | 36 |
| cellular divalent inorganic cation homeostasis [GO:0072503] | 4.7E-02 | 11 |

| Cellular Component GO Terms | p-Value | Matches |
|---|---|---|
| cell periphery [GO:0071944] | 1.3E-16 | 130 |
| endoplasmic reticulum [GO:0005783] | 3.5E-16 | 114 |
| intrinsic component of membrane [GO:0031224] | 1.4E-15 | 198 |
| endomembrane system [GO:0012505] | 8.4E-14 | 149 |
| membrane part [GO:0044425] | 4.4E-13 | 217 |
| membrane [GO:0016020] | 2.0E-12 | 250 |
| plasma membrane [GO:0005886] | 2.6E-12 | 94 |
| storage vacuole [GO:0000322] | 2.9E-12 | 82 |
| fungal-type vacuole [GO:0000324] | 2.9E-12 | 82 |
| lytic vacuole [GO:0000323] | 3.3E-12 | 82 |
| integral component of membrane [GO:0016021] | 4.2E-11 | 180 |
| vacuole [GO:0005773] | 4.4E-10 | 85 |
| endoplasmic reticulum part [GO:0044432] | 1.2E-09 | 71 |
| extracellular region [GO:0005576] | 1.4E-09 | 34 |
| endoplasmic reticulum subcompartment [GO:0098827] | 3.6E-09 | 68 |
| organelle subcompartment [GO:0031984] | 7.4E-09 | 82 |
| endoplasmic reticulum membrane [GO:0005789] | 3.2E-08 | 65 |
| nuclear outer membrane-endoplasmic reticulum membrane network [GO:0042175] | 1.4E-07 | 65 |
| cytoplasmic part [GO:0044444] | 1.4E-06 | 320 |
| cell wall [GO:0005618] | 8.0E-06 | 31 |
| external encapsulating structure [GO:0030312] | 8.0E-06 | 31 |
| fungal-type cell wall [GO:0009277] | 1.1E-05 | 30 |
| plasma membrane part [GO:0044459] | 2.4E-04 | 33 |
| vacuolar membrane [GO:0005774] | 2.9E-04 | 45 |
| anchored component of membrane [GO:0031225] | 5.8E-04 | 18 |
| vacuolar part [GO:0044437] | 9.7E-04 | 45 |
| intrinsic component of plasma membrane [GO:0031226] | 1.1E-03 | 26 |
| cell part [GO:0044464] | 4.0E-03 | 443 |
| bounding membrane of organelle [GO:0098588] | 4.0E-03 | 75 |
| integral component of plasma membrane [GO:0005887] | 4.4E-03 | 24 |

| | | |
|---|---|---|
| cytoplasm [GO:0005737] | 4.4E-03 | 381 |
| cell [GO:0005623] | 6.7E-03 | 443 |
| fungal-type vacuole membrane [GO:0000329] | 1.3E-02 | 35 |
| lytic vacuole membrane [GO:0098852] | 1.3E-02 | 35 |
| membrane-bounded organelle [GO:0043227] | 4.3E-02 | 364 |
| intracellular organelle [GO:0043229] | 4.3E-02 | 383 |
| intracellular membrane-bounded organelle [GO:0043231] | 4.6E-02 | 360 |
| organelle [GO:0043226] | 4.6E-02 | 383 |

| Molecular Function GO Terms | p-Value | Matches |
|---|---|---|
| metal ion transmembrane transporter activity [GO:0046873] | 2.5E-11 | 26 |
| transmembrane transporter activity [GO:0022857] | 3.4E-10 | 70 |
| transporter activity [GO:0005215] | 5.6E-09 | 71 |
| inorganic molecular entity transmembrane transporter activity [GO:0015318] | 8.5E-09 | 52 |
| ion transmembrane transporter activity [GO:0015075] | 2.6E-08 | 54 |
| inorganic cation transmembrane transporter activity [GO:0022890] | 3.3E-06 | 35 |
| transition metal ion transmembrane transporter activity [GO:0046915] | 1.3E-05 | 15 |
| cation transmembrane transporter activity [GO:0008324] | 1.3E-05 | 39 |

**Supplementary Table 5. Mitochondrial Profiling Unique vs Mitochondria tagging: GO Terms.**

| Biological Process GO Terms | p-Value | Matches |
|---|---|---|
| mitochondrial gene expression [GO:0140053] | 6.7E-09 | 35 |
| mitochondrial RNA metabolic process [GO:0000959] | 8.5E-09 | 17 |
| mitochondrial translation [GO:0032543] | 3.6E-05 | 27 |
| small molecule metabolic process [GO:0044281] | 4.7E-05 | 70 |
| respiratory electron transport chain [GO:0022904] | 9.2E-05 | 13 |
| cellular respiration [GO:0045333] | 1.2E-04 | 20 |
| tRNA aminoacylation for mitochondrial protein translation [GO:0070127] | 1.5E-04 | 8 |
| amino acid activation [GO:0043038] | 5.0E-04 | 12 |
| tRNA aminoacylation [GO:0043039] | 5.0E-04 | 12 |
| tRNA aminoacylation for protein translation [GO:0006418] | 1.5E-03 | 11 |
| energy derivation by oxidation of organic compounds [GO:0015980] | 1.7E-03 | 23 |
| ornithine biosynthetic process [GO:0006592] | 1.9E-02 | 4 |
| electron transport chain [GO:0022900] | 2.1E-02 | 13 |
| mitochondrion organization [GO:0007005] | 4.8E-02 | 29 |

| Cellular Component GO Terms | p-Value | Matches |
|---|---|---|
| mitochondrion [GO:0005739] | 9.4E-45 | 163 |
| mitochondrial part [GO:0044429] | 8.9E-43 | 118 |
| mitochondrial envelope [GO:0005740] | 2.9E-27 | 82 |
| mitochondrial membrane [GO:0031966] | 1.4E-25 | 76 |
| mitochondrial inner membrane [GO:0005743] | 1.8E-24 | 60 |
| organelle inner membrane [GO:0019866] | 2.4E-23 | 60 |
| organelle envelope [GO:0031967] | 1.2E-20 | 85 |
| envelope [GO:0031975] | 1.2E-20 | 85 |
| mitochondrial matrix [GO:0005759] | 2.8E-10 | 40 |
| mitochondrial membrane part [GO:0044455] | 7.8E-08 | 32 |
| organelle membrane [GO:0031090] | 2.3E-07 | 86 |
| cytoplasmic part [GO:0044444] | 3.7E-05 | 205 |
| mitochondrial protein complex [GO:0098798] | 4.3E-03 | 26 |

| | | |
|---|---|---|
| chloroplast [GO:0009507] | 4.3E-03 | 4 |
| chloroplast thylakoid [GO:0009534] | 4.3E-03 | 4 |
| chloroplast thylakoid membrane [GO:0009535] | 4.3E-03 | 4 |
| plastid [GO:0009536] | 4.3E-03 | 4 |
| thylakoid [GO:0009579] | 4.3E-03 | 4 |
| plastid thylakoid [GO:0031976] | 4.3E-03 | 4 |
| photosynthetic membrane [GO:0034357] | 4.3E-03 | 4 |
| thylakoid membrane [GO:0042651] | 4.3E-03 | 4 |
| chloroplast part [GO:0044434] | 4.3E-03 | 4 |
| plastid part [GO:0044435] | 4.3E-03 | 4 |
| thylakoid part [GO:0044436] | 4.3E-03 | 4 |
| plastid thylakoid membrane [GO:0055035] | 4.3E-03 | 4 |
| respiratory chain [GO:0070469] | 9.9E-03 | 10 |
| membrane [GO:0016020] | 1.5E-02 | 140 |
| respiratory chain complex [GO:0098803] | 1.8E-02 | 9 |
| membrane part [GO:0044425] | 2.1E-02 | 118 |
| mitochondrial respiratory chain [GO:0005746] | 2.9E-02 | 9 |
| extrinsic component of mitochondrial inner membrane [GO:0031314] | 3.0E-02 | 8 |
| integral component of mitochondrial membrane [GO:0032592] | 4.3E-02 | 12 |

| Molecular Function GO Terms | p-Value | Matches |
|---|---|---|
| aminoacyl-tRNA ligase activity [GO:0004812] | 6.5E-04 | 11 |
| ligase activity, forming carbon-oxygen bonds [GO:0016875] | 6.5E-04 | 11 |
| catalytic activity, acting on a tRNA [GO:0140101] | 6.8E-04 | 18 |
| electron transfer activity [GO:0009055] | 1.1E-02 | 12 |

**Supplementary Table 6. ER tagging Unique vs Mitochondria Tagging: GO Terms.**

| Biological Process GO Terms | p-Value | Matches |
|---|---|---|
| glycoprotein metabolic process [GO:0009100] | 1.4E-25 | 55 |
| glycosylation [GO:0070085] | 1.4E-25 | 55 |
| glycoprotein biosynthetic process [GO:0009101] | 1.5E-24 | 52 |
| protein glycosylation [GO:0006486] | 7.4E-24 | 50 |
| macromolecule glycosylation [GO:0043413] | 7.4E-24 | 50 |
| mannosylation [GO:0097502] | 1.2E-20 | 36 |
| transmembrane transport [GO:0055085] | 8.8E-20 | 135 |
| transport [GO:0006810] | 2.0E-18 | 276 |
| establishment of localization [GO:0051234] | 4.5E-17 | 279 |
| localization [GO:0051179] | 6.0E-17 | 307 |
| protein O-linked glycosylation [GO:0006493] | 1.0E-14 | 21 |
| ion transport [GO:0006811] | 1.3E-14 | 110 |
| lipid metabolic process [GO:0006629] | 3.0E-13 | 100 |
| protein N-linked glycosylation [GO:0006487] | 2.5E-12 | 30 |
| membrane lipid metabolic process [GO:0006643] | 7.9E-12 | 40 |
| cellular lipid metabolic process [GO:0044255] | 1.7E-11 | 92 |
| ion transmembrane transport [GO:0034220] | 3.3E-11 | 80 |
| anion transport [GO:0006820] | 1.3E-10 | 65 |
| organic substance transport [GO:0071702] | 1.4E-10 | 193 |
| membrane lipid biosynthetic process [GO:0046467] | 2.8E-10 | 34 |
| lipid biosynthetic process [GO:0008610] | 1.2E-09 | 67 |
| carbohydrate derivative biosynthetic process [GO:1901137] | 2.5E-09 | 77 |
| nitrogen compound transport [GO:0071705] | 2.0E-08 | 170 |
| organic anion transport [GO:0015711] | 1.1E-07 | 52 |
| cation transport [GO:0006812] | 1.2E-07 | 65 |
| anion transmembrane transport [GO:0098656] | 2.6E-06 | 38 |
| protein lipidation [GO:0006497] | 2.7E-06 | 24 |
| lipoprotein biosynthetic process [GO:0042158] | 2.7E-06 | 24 |
| lipoprotein metabolic process [GO:0042157] | 4.6E-06 | 24 |
| protein mannosylation [GO:0035268] | 6.8E-06 | 11 |

| | | |
|---|---|---|
| GPI anchor metabolic process [GO:0006505] | 1.0E-05 | 19 |
| glycolipid biosynthetic process [GO:0009247] | 1.0E-05 | 19 |
| GPI anchor biosynthetic process [GO:0006506] | 1.2E-05 | 18 |
| organic acid transport [GO:0015849] | 1.4E-05 | 33 |
| glycolipid metabolic process [GO:0006664] | 1.5E-05 | 20 |
| liposaccharide metabolic process [GO:1903509] | 1.5E-05 | 20 |
| cation transmembrane transport [GO:0098655] | 3.1E-05 | 53 |
| carboxylic acid transport [GO:0046942] | 4.2E-05 | 32 |
| carbohydrate derivative metabolic process [GO:1901135] | 4.7E-05 | 88 |
| cell wall macromolecule metabolic process [GO:0044036] | 4.8E-05 | 23 |
| maintenance of protein localization in endoplasmic reticulum [GO:0035437] | 5.0E-05 | 10 |
| protein localization to endoplasmic reticulum [GO:0070972] | 1.2E-04 | 24 |
| cell wall mannoprotein biosynthetic process [GO:0000032] | 1.3E-04 | 13 |
| mannoprotein metabolic process [GO:0006056] | 1.3E-04 | 13 |
| mannoprotein biosynthetic process [GO:0006057] | 1.3E-04 | 13 |
| cell wall glycoprotein biosynthetic process [GO:0031506] | 1.3E-04 | 13 |
| sphingolipid metabolic process [GO:0006665] | 2.7E-04 | 22 |
| protein retention in ER lumen [GO:0006621] | 3.6E-04 | 9 |
| protein O-linked mannosylation [GO:0035269] | 3.6E-04 | 9 |
| response to endoplasmic reticulum stress [GO:0034976] | 5.0E-04 | 30 |
| phosphatidylinositol biosynthetic process [GO:0006661] | 9.5E-04 | 20 |
| response to unfolded protein [GO:0006986] | 9.6E-04 | 21 |
| organic acid transmembrane transport [GO:1903825] | 1.7E-03 | 24 |
| cell wall macromolecule biosynthetic process [GO:0044038] | 3.4E-03 | 19 |
| cellular component macromolecule biosynthetic process [GO:0070589] | 3.4E-03 | 19 |
| phospholipid biosynthetic process [GO:0008654] | 3.7E-03 | 34 |
| response to topologically incorrect protein [GO:0035966] | 3.8E-03 | 25 |
| phospholipid metabolic process [GO:0006644] | 3.9E-03 | 43 |
| sphingolipid biosynthetic process [GO:0030148] | 4.9E-03 | 17 |
| carboxylic acid transmembrane transport [GO:1905039] | 5.1E-03 | 23 |
| cell wall biogenesis [GO:0042546] | 5.9E-03 | 33 |
| ammonium transport [GO:0015696] | 6.4E-03 | 13 |
| ammonium transmembrane transport [GO:0072488] | 8.7E-03 | 11 |

| | | |
|---|---|---|
| amino acid transport [GO:0006865] | 9.1E-03 | 21 |
| drug transmembrane transport [GO:0006855] | 1.0E-02 | 20 |
| drug transport [GO:0015893] | 1.1E-02 | 27 |
| glycerolipid biosynthetic process [GO:0045017] | 1.1E-02 | 27 |
| amino acid transmembrane transport [GO:0003333] | 1.2E-02 | 18 |
| nucleobase transport [GO:0015851] | 1.2E-02 | 8 |
| regulation of membrane lipid distribution [GO:0097035] | 1.5E-02 | 12 |
| cell wall organization or biogenesis [GO:0071554] | 1.6E-02 | 65 |
| glycerophospholipid biosynthetic process [GO:0046474] | 1.7E-02 | 26 |
| glycerolipid metabolic process [GO:0046486] | 1.8E-02 | 37 |
| maintenance of protein localization in organelle [GO:0072595] | 2.0E-02 | 10 |
| ERAD pathway [GO:0036503] | 2.0E-02 | 20 |
| cellular response to unfolded protein [GO:0034620] | 2.7E-02 | 17 |
| lipid translocation [GO:0034204] | 3.6E-02 | 11 |
| endoplasmic reticulum unfolded protein response [GO:0030968] | 4.0E-02 | 14 |
| oligosaccharide-lipid intermediate biosynthetic process [GO:0006490] | 4.5E-02 | 9 |

| Cellular Component GO Terms | p-Value | Matches |
|---|---|---|
| intrinsic component of membrane [GO:0031224] | 9.1E-135 | 516 |
| integral component of membrane [GO:0016021] | 5.9E-131 | 500 |
| membrane part [GO:0044425] | 8.8E-104 | 526 |
| endoplasmic reticulum [GO:0005783] | 1.4E-100 | 298 |
| endomembrane system [GO:0012505] | 1.9E-87 | 367 |
| membrane [GO:0016020] | 2.6E-79 | 554 |
| organelle subcompartment [GO:0031984] | 1.7E-64 | 219 |
| endoplasmic reticulum part [GO:0044432] | 6.5E-62 | 187 |
| endoplasmic reticulum subcompartment [GO:0098827] | 3.7E-58 | 178 |
| endoplasmic reticulum membrane [GO:0005789] | 1.0E-56 | 174 |
| nuclear outer membrane-endoplasmic reticulum membrane network [GO:0042175] | 1.1E-55 | 176 |
| lytic vacuole [GO:0000323] | 8.7E-36 | 164 |
| storage vacuole [GO:0000322] | 2.8E-35 | 163 |
| fungal-type vacuole [GO:0000324] | 2.8E-35 | 163 |
| vacuole [GO:0005773] | 7.7E-35 | 177 |
| bounding membrane of organelle [GO:0098588] | 3.5E-30 | 187 |
| Golgi apparatus [GO:0005794] | 4.5E-22 | 107 |
| vacuolar part [GO:0044437] | 8.6E-22 | 105 |
| vacuolar membrane [GO:0005774] | 4.0E-19 | 98 |
| organelle membrane [GO:0031090] | 3.2E-18 | 226 |
| cytoplasmic part [GO:0044444] | 7.1E-18 | 584 |
| fungal-type vacuole membrane [GO:0000329] | 2.7E-16 | 81 |
| lytic vacuole membrane [GO:0098852] | 2.7E-16 | 81 |
| whole membrane [GO:0098805] | 2.9E-16 | 147 |
| intrinsic component of endoplasmic reticulum membrane [GO:0031227] | 3.6E-16 | 44 |
| cell periphery [GO:0071944] | 5.0E-16 | 193 |
| integral component of endoplasmic reticulum membrane [GO:0030176] | 1.6E-15 | 43 |
| Golgi membrane [GO:0000139] | 1.4E-14 | 60 |
| Golgi subcompartment [GO:0098791] | 5.7E-14 | 68 |
| plasma membrane [GO:0005886] | 8.0E-14 | 143 |

| | | |
|---|---|---|
| Golgi apparatus part [GO:0044431] | 6.5E-13 | 72 |
| intrinsic component of plasma membrane [GO:0031226] | 3.8E-09 | 47 |
| integral component of plasma membrane [GO:0005887] | 2.8E-08 | 44 |
| integral component of Golgi membrane [GO:0030173] | 6.7E-08 | 20 |
| intrinsic component of Golgi membrane [GO:0031228] | 6.7E-08 | 20 |
| plasma membrane part [GO:0044459] | 2.2E-07 | 54 |
| intracellular membrane-bounded organelle [GO:0043231] | 3.9E-07 | 650 |
| membrane-bounded organelle [GO:0043227] | 1.1E-06 | 655 |
| organelle part [GO:0044422] | 2.6E-05 | 480 |
| endoplasmic reticulum lumen [GO:0005788] | 6.8E-05 | 12 |
| intracellular organelle part [GO:0044446] | 8.7E-05 | 476 |
| mannosyltransferase complex [GO:0031501] | 1.8E-04 | 12 |
| dolichyl-phosphate-mannose-protein mannosyltransferase complex [GO:0031502] | 6.1E-04 | 7 |
| endoplasmic reticulum-Golgi intermediate compartment [GO:0005793] | 2.3E-03 | 14 |
| nuclear membrane [GO:0031965] | 3.3E-03 | 27 |
| dolichyl-phosphate-mannose-protein mannosyltransferase Pmt1p-Pmt2p dimer complex [GO:0097582] | 4.7E-03 | 6 |
| COPII-coated ER to Golgi transport vesicle [GO:0030134] | 1.0E-02 | 17 |
| intracellular organelle [GO:0043229] | 4.7E-02 | 667 |

| Molecular Function GO Terms | p-Value | Matches |
|---|---|---|
| transporter activity [GO:0005215] | 1.6E-25 | 126 |
| transmembrane transporter activity [GO:0022857] | 1.0E-24 | 119 |
| transferase activity, transferring hexosyl groups [GO:0016758] | 8.2E-22 | 49 |
| mannosyltransferase activity [GO:0000030] | 9.4E-21 | 34 |
| transferase activity, transferring glycosyl groups [GO:0016757] | 1.4E-19 | 53 |
| ion transmembrane transporter activity [GO:0015075] | 1.0E-16 | 86 |
| inorganic molecular entity transmembrane transporter activity [GO:0015318] | 1.2E-10 | 71 |
| anion transmembrane transporter activity [GO:0008509] | 3.5E-10 | 42 |
| cation transmembrane transporter activity [GO:0008324] | 2.2E-09 | 58 |
| active transmembrane transporter activity [GO:0022804] | 6.0E-09 | 53 |
| alpha-1,2-mannosyltransferase activity [GO:0000026] | 1.4E-07 | 12 |
| organic anion transmembrane transporter activity [GO:0008514] | 1.6E-07 | 34 |
| drug transmembrane transporter activity [GO:0015238] | 1.3E-06 | 26 |
| organic acid transmembrane transporter activity [GO:0005342] | 1.7E-06 | 28 |
| secondary active transmembrane transporter activity [GO:0015291] | 2.0E-06 | 33 |
| carboxylic acid transmembrane transporter activity [GO:0046943] | 6.2E-06 | 27 |
| dolichyl-phosphate-mannose-protein mannosyltransferase activity [GO:0004169] | 6.5E-04 | 7 |
| ammonium transmembrane transporter activity [GO:0008519] | 6.9E-04 | 11 |
| antiporter activity [GO:0015297] | 2.4E-03 | 15 |
| nucleobase transmembrane transporter activity [GO:0015205] | 2.7E-03 | 8 |
| nucleobase-containing compound transmembrane transporter activity [GO:0015932] | 3.4E-03 | 13 |
| alpha-1,3-mannosyltransferase activity [GO:0000033] | 5.7E-03 | 6 |
| L-tyrosine transmembrane transporter activity [GO:0005302] | 5.7E-03 | 6 |
| exopeptidase activity [GO:0008238] | 6.2E-03 | 14 |
| amino acid transmembrane transporter activity [GO:0015171] | 7.3E-03 | 17 |
| carbohydrate derivative transmembrane transporter activity [GO:1901505] | 1.5E-02 | 12 |
| inorganic cation transmembrane transporter activity [GO:0022890] | 1.7E-02 | 37 |
| aromatic amino acid transmembrane transporter activity [GO:0015173] | 1.9E-02 | 7 |
| signaling receptor activity [GO:0038023] | 2.2E-02 | 9 |
| nucleotide-sugar transmembrane transporter activity [GO:0005338] | 5.0E-02 | 5 |

**Supplementary Table 7. Mitochondria Tagging Unique vs ER tagging: GO Terms.**

| Biological Process GO Terms | p-Value | Matches |
|---|---|---|
| mitochondrion organization [GO:0007005] | 1.1E-10 | 49 |
| mitochondrial gene expression [GO:0140053] | 2.3E-08 | 38 |
| protein localization to mitochondrion [GO:0070585] | 1.6E-05 | 19 |
| establishment of protein localization to mitochondrion [GO:0072655] | 1.6E-05 | 19 |
| mitochondrial translation [GO:0032543] | 3.9E-05 | 30 |
| mitochondrial transport [GO:0006839] | 7.3E-05 | 21 |
| mitochondrial genome maintenance [GO:0000002] | 2.8E-04 | 14 |
| protein targeting to mitochondrion [GO:0006626] | 4.1E-03 | 15 |
| mitochondrial transmembrane transport [GO:1990542] | 9.4E-03 | 15 |
| protein transmembrane transport [GO:0071806] | 9.8E-03 | 17 |
| protein targeting [GO:0006605] | 1.8E-02 | 28 |
| positive regulation of mitochondrial translation [GO:0070131] | 2.6E-02 | 7 |
| carboxylic acid metabolic process [GO:0019752] | 4.1E-02 | 45 |

| Cellular Component GO Terms | p-Value | Matches |
|---|---|---|
| mitochondrion [GO:0005739] | 2.4E-43 | 175 |
| mitochondrial part [GO:0044429] | 4.7E-39 | 122 |
| mitochondrial envelope [GO:0005740] | 6.2E-23 | 82 |
| mitochondrial membrane [GO:0031966] | 1.3E-21 | 76 |
| organelle envelope [GO:0031967] | 1.1E-17 | 87 |
| envelope [GO:0031975] | 1.1E-17 | 87 |
| mitochondrial inner membrane [GO:0005743] | 1.4E-16 | 54 |
| organelle inner membrane [GO:0019866] | 2.3E-16 | 55 |
| mitochondrial matrix [GO:0005759] | 5.1E-16 | 51 |
| cytoplasm [GO:0005737] | 3.6E-10 | 292 |
| mitochondrial protein complex [GO:0098798] | 2.8E-09 | 39 |
| cytoplasmic part [GO:0044444] | 9.8E-09 | 241 |
| intracellular [GO:0005622] | 1.5E-08 | 318 |
| mitochondrial membrane part [GO:0044455] | 1.2E-07 | 34 |
| intracellular part [GO:0044424] | 1.5E-07 | 317 |
| cell part [GO:0044464] | 2.7E-07 | 320 |
| cell [GO:0005623] | 4.5E-07 | 320 |
| intracellular organelle [GO:0043229] | 1.9E-06 | 288 |
| organelle membrane [GO:0031090] | 2.0E-06 | 92 |
| organelle [GO:0043226] | 2.0E-06 | 288 |
| membrane-bounded organelle [GO:0043227] | 2.9E-05 | 273 |
| intrinsic component of mitochondrial membrane [GO:0098573] | 8.2E-05 | 17 |
| intracellular membrane-bounded organelle [GO:0043231] | 9.8E-05 | 269 |
| organellar ribosome [GO:0000313] | 1.4E-04 | 19 |
| mitochondrial ribosome [GO:0005761] | 1.4E-04 | 19 |
| integral component of mitochondrial membrane [GO:0032592] | 1.2E-03 | 15 |

| | | |
|---|---|---|
| nucleoid [GO:0009295] | 2.3E-03 | 9 |
| mitochondrial nucleoid [GO:0042645] | 2.3E-03 | 9 |
| intrinsic component of mitochondrial inner membrane [GO:0031304] | 3.4E-03 | 11 |
| intracellular organelle part [GO:0044446] | 4.4E-02 | 196 |

| Molecular Function GO Terms | p-Value | Matches |
| --- | --- | --- |
| protein transmembrane transporter activity [GO:0008320] | 2.1E-02 | 9 |
| macromolecule transmembrane transporter activity [GO:0022884] | 2.1E-02 | 9 |
| peptide transmembrane transporter activity [GO:1904680] | 3.1E-02 | 10 |
| protein transporter activity [GO:0008565] | 4.1E-02 | 9 |

**Supplementary Table 8. Yeast Dual-tagged RNAs: GO Terms.**

| Biological Process GO Terms | p-Value | Matches |
|---|---|---|
| ion transport [GO:0006811] | 7.1E-22 | 67 |
| transmembrane transport [GO:0055085] | 2.8E-19 | 70 |
| metal ion transport [GO:0030001] | 3.3E-18 | 31 |
| ion transmembrane transport [GO:0034220] | 8.6E-17 | 50 |
| metal ion homeostasis [GO:0055065] | 2.4E-16 | 34 |
| cation homeostasis [GO:0055080] | 1.0E-14 | 37 |
| inorganic ion homeostasis [GO:0098771] | 1.1E-13 | 35 |
| ion homeostasis [GO:0050801] | 2.0E-13 | 37 |
| cellular metal ion homeostasis [GO:0006875] | 1.1E-12 | 29 |
| cation transport [GO:0006812] | 1.5E-12 | 41 |
| inorganic ion transmembrane transport [GO:0098660] | 4.6E-12 | 35 |
| transition metal ion homeostasis [GO:0055076] | 8.1E-12 | 26 |
| chemical homeostasis [GO:0048878] | 8.5E-12 | 39 |
| transition metal ion transport [GO:0000041] | 1.1E-11 | 20 |
| cellular cation homeostasis [GO:0030003] | 2.0E-11 | 32 |
| inorganic cation transmembrane transport [GO:0098662] | 3.7E-11 | 32 |
| cation transmembrane transport [GO:0098655] | 1.7E-10 | 35 |
| cellular ion homeostasis [GO:0006873] | 2.1E-10 | 32 |
| cellular chemical homeostasis [GO:0055082] | 4.4E-10 | 34 |
| cellular homeostasis [GO:0019725] | 4.8E-10 | 38 |
| cell wall organization or biogenesis [GO:0071554] | 1.3E-09 | 43 |
| cellular transition metal ion homeostasis [GO:0046916] | 5.7E-09 | 22 |
| homeostatic process [GO:0042592] | 3.3E-08 | 44 |
| external encapsulating structure organization [GO:0045229] | 4.0E-08 | 37 |
| cell wall organization [GO:0071555] | 4.0E-08 | 37 |
| iron ion transport [GO:0006826] | 2.1E-07 | 12 |
| anion transport [GO:0006820] | 4.9E-07 | 31 |

| | | |
|---|---|---|
| transport [GO:0006810] | 5.7E-07 | 100 |
| lipid metabolic process [GO:0006629] | 1.1E-06 | 42 |
| fungal-type cell wall organization or biogenesis [GO:0071852] | 1.4E-06 | 33 |
| establishment of localization [GO:0051234] | 1.8E-06 | 101 |
| localization [GO:0051179] | 7.1E-06 | 109 |
| iron ion homeostasis [GO:0055072] | 8.7E-06 | 16 |
| anion transmembrane transport [GO:0098656] | 9.0E-06 | 21 |
| divalent inorganic cation transport [GO:0072511] | 1.0E-05 | 13 |
| cellular lipid metabolic process [GO:0044255] | 2.2E-05 | 38 |
| membrane lipid metabolic process [GO:0006643] | 3.0E-05 | 18 |
| divalent metal ion transport [GO:0070838] | 6.3E-05 | 12 |
| membrane lipid biosynthetic process [GO:0046467] | 6.3E-05 | 16 |
| fungal-type cell wall organization [GO:0031505] | 7.2E-05 | 28 |
| lipid biosynthetic process [GO:0008610] | 8.2E-05 | 29 |
| glycerophospholipid metabolic process [GO:0006650] | 1.7E-04 | 21 |
| glycerolipid metabolic process [GO:0046486] | 2.2E-04 | 22 |
| regulation of biological quality [GO:0065008] | 2.2E-04 | 52 |
| phospholipid metabolic process [GO:0006644] | 2.2E-04 | 24 |
| divalent inorganic cation homeostasis [GO:0072507] | 2.6E-04 | 11 |
| glycolipid metabolic process [GO:0006664] | 1.3E-03 | 11 |
| liposaccharide metabolic process [GO:1903509] | 1.3E-03 | 11 |
| manganese ion transport [GO:0006828] | 2.1E-03 | 6 |
| cellular divalent inorganic cation homeostasis [GO:0072503] | 2.2E-03 | 10 |
| organic anion transport [GO:0015711] | 3.6E-03 | 22 |
| glycolipid biosynthetic process [GO:0009247] | 4.1E-03 | 10 |
| cellular iron ion homeostasis [GO:0006879] | 5.6E-03 | 12 |
| amino acid transmembrane transport [GO:0003333] | 6.8E-03 | 11 |
| glycerolipid biosynthetic process [GO:0045017] | 8.5E-03 | 15 |
| phosphatidylinositol metabolic process [GO:0046488] | 1.3E-02 | 14 |
| amino acid transport [GO:0006865] | 1.3E-02 | 12 |

| | | |
|---|---|---|
| protein lipidation [GO:0006497] | 2.2E-02 | 11 |
| lipoprotein biosynthetic process [GO:0042158] | 2.2E-02 | 11 |
| phospholipid biosynthetic process [GO:0008654] | 2.3E-02 | 17 |
| lipoprotein metabolic process [GO:0042157] | 2.7E-02 | 11 |
| glycerophospholipid biosynthetic process [GO:0046474] | 2.9E-02 | 14 |
| cofactor transport [GO:0051181] | 3.0E-02 | 9 |
| sodium ion transmembrane transport [GO:0035725] | 3.4E-02 | 5 |
| manganese ion transmembrane transport [GO:0071421] | 3.4E-02 | 5 |
| GPI anchor metabolic process [GO:0006505] | 3.9E-02 | 9 |

| Cellular Component GO Terms | p-Value | Matches |
|---|---|---|
| intrinsic component of membrane [GO:0031224] | 1.2E-58 | 192 |
| membrane part [GO:0044425] | 1.3E-47 | 195 |
| integral component of membrane [GO:0016021] | 1.0E-46 | 175 |
| membrane [GO:0016020] | 1.1E-39 | 205 |
| endoplasmic reticulum [GO:0005783] | 1.0E-31 | 103 |
| cell periphery [GO:0071944] | 6.8E-30 | 111 |
| endomembrane system [GO:0012505] | 3.7E-25 | 121 |
| storage vacuole [GO:0000322] | 1.8E-20 | 71 |
| fungal-type vacuole [GO:0000324] | 1.8E-20 | 71 |
| lytic vacuole [GO:0000323] | 2.0E-20 | 71 |
| plasma membrane [GO:0005886] | 3.0E-20 | 79 |
| endoplasmic reticulum part [GO:0044432] | 7.9E-20 | 66 |
| endoplasmic reticulum subcompartment [GO:0098827] | 1.0E-18 | 63 |
| organelle subcompartment [GO:0031984] | 3.2E-18 | 73 |
| vacuole [GO:0005773] | 4.0E-18 | 73 |
| endoplasmic reticulum membrane [GO:0005789] | 4.6E-17 | 60 |
| nuclear outer membrane-endoplasmic reticulum membrane network [GO:0042175] | 2.6E-16 | 60 |
| extracellular region [GO:0005576] | 6.5E-14 | 31 |
| intrinsic component of plasma membrane [GO:0031226] | 2.2E-10 | 28 |
| cytoplasmic part [GO:0044444] | 6.4E-10 | 205 |
| plasma membrane part [GO:0044459] | 7.4E-10 | 32 |
| cell wall [GO:0005618] | 1.6E-09 | 28 |
| external encapsulating structure [GO:0030312] | 1.6E-09 | 28 |
| integral component of plasma membrane [GO:0005887] | 2.6E-09 | 26 |
| fungal-type cell wall [GO:0009277] | 3.3E-09 | 27 |
| bounding membrane of organelle [GO:0098588] | 9.8E-08 | 61 |
| vacuolar membrane [GO:0005774] | 3.5E-07 | 37 |

| | | |
|---|---|---|
| anchored component of membrane [GO:0031225] | 9.5E-07 | 17 |
| vacuolar part [GO:0044437] | 1.2E-06 | 37 |
| whole membrane [GO:0098805] | 2.9E-06 | 54 |
| organelle membrane [GO:0031090] | 6.6E-06 | 78 |
| integral component of endoplasmic reticulum membrane [GO:0030176] | 1.5E-04 | 16 |
| intrinsic component of endoplasmic reticulum membrane [GO:0031227] | 1.7E-04 | 16 |
| fungal-type vacuole membrane [GO:0000329] | 2.5E-03 | 26 |
| lytic vacuole membrane [GO:0098852] | 2.5E-03 | 26 |
| intracellular membrane-bounded organelle [GO:0043231] | 1.6E-02 | 217 |
| Golgi apparatus [GO:0005794] | 2.1E-02 | 29 |

| **Molecular Function GO Terms** | **p-Value** | **Matches** |
|---|---|---|
| transmembrane transporter activity [GO:0022857] | 3.1E-22 | 66 |
| transporter activity [GO:0005215] | 3.1E-20 | 66 |
| metal ion transmembrane transporter activity [GO:0046873] | 1.2E-17 | 26 |
| inorganic molecular entity transmembrane transporter activity [GO:0015318] | 9.8E-17 | 48 |
| ion transmembrane transporter activity [GO:0015075] | 1.2E-16 | 50 |
| cation transmembrane transporter activity [GO:0008324] | 2.4E-11 | 36 |
| inorganic cation transmembrane transporter activity [GO:0022890] | 3.7E-11 | 32 |
| transition metal ion transmembrane transporter activity [GO:0046915] | 3.4E-09 | 15 |
| active transmembrane transporter activity [GO:0022804] | 3.5E-04 | 24 |
| secondary active transmembrane transporter activity [GO:0015291] | 5.8E-04 | 17 |
| anion transmembrane transporter activity [GO:0008509] | 1.2E-03 | 18 |
| organic anion transmembrane transporter activity [GO:0008514] | 2.0E-03 | 16 |
| manganese ion transmembrane transporter activity [GO:0005384] | 1.6E-02 | 5 |
| sodium ion transmembrane transporter activity [GO:0015081] | 1.6E-02 | 5 |
| cofactor transmembrane transporter activity [GO:0051184] | 1.8E-02 | 7 |
| transferase activity, transferring hexosyl groups [GO:0016758] | 2.5E-02 | 14 |
| amino acid transmembrane transporter activity [GO:0015171] | 2.6E-02 | 10 |
| potassium ion transmembrane transporter activity [GO:0015079] | 3.0E-02 | 6 |

| Pathways | p-Value | Matches |
|---|---|---|
| sphingolipid metabolism | 9.9E-03 | 6 |

**Supplementary Table 9. 45 RNAs tagged about equally by both ER- and Mito-PUP.**

| Systematic Name | Feature Type | ER-PUP Tier | Mito-PUP Tier |
|---|---|---|---|
| YER019W | ORF | 1 | 1 |
| YHR071W | ORF | 1 | 1 |
| YLR056W | ORF | 1 | 1 |
| YEL017C-A | ORF | 1 | 1 |
| YHL040C | ORF | 1 | 2 |
| YNL327W | ORF | 1 | 2 |
| YCR024C-B | ORF | 1 | 2 |
| YHL047C | ORF | 1 | 3 |
| YDR534C | ORF | 1 | 3 |
| YKR072C | ORF | 1 | 4 |
| YDR481C | ORF | 2 | 3 |
| YDR072C | ORF | 2 | 3 |
| YIL117C | ORF | 2 | 3 |
| YLR188W | ORF | 2 | 3 |
| YCR015C | ORF | 3 | 2 |
| YOR306C | ORF | 3 | 2 |
| YDR422C | ORF | 3 | 3 |
| YPR140W | ORF | 3 | 3 |
| YDR319C | ORF | 3 | 3 |
| YDR270W | ORF | 3 | 3 |
| YLR138W | ORF | 3 | 3 |
| YDR073W | ORF | 3 | 3 |
| YNL186W | ORF | 3 | 3 |
| YDR320C-A | ORF | 4 | 2 |
| snR7-L | snRNA gene | 3 | 4 |
| YGR146C-A | ORF | 3 | 4 |
| YLR226W | ORF | 3 | 4 |

| | | | |
|---|---|---|---|
| YLR404W | ORF | 3 | 4 |
| YIL090W | ORF | 3 | 4 |
| YBR262C | ORF | 3 | 4 |
| YDR124W | ORF | 4 | 3 |
| YKR052C | ORF | 4 | 3 |
| YDR374W-A | ORF | 4 | 3 |
| YGR171C | ORF | 3 | 5 |
| YNR045W | ORF | 4 | 4 |
| YHL028W | ORF | 4 | 4 |
| YPL216W | ORF | 4 | 4 |
| YMR195W | ORF | 4 | 4 |
| YHR143W-A | ORF | 4 | 4 |
| YDL232W | ORF | 4 | 4 |
| YJR082C | ORF | 5 | 3 |
| YNL211C | ORF | 4 | 5 |
| YBL051C | ORF | 4 | 5 |
| YOR384W | ORF | 5 | 5 |
| YEL061C | ORF | 5 | 5 |

**Supplementary Table 10. Dual-localization conservation in yeast and HEK293T cells.**

**<u>Yeast (BY4742):</u>**

| Systematic Name | ER-PUP Tier | Mito-PUP Tier |
|---|---|---|
| YKL008C | 1 | 2 |
| YGL054C | 1 | 2 |
| YDR107C | 2 | 2 |
| YHR079C | 1 | 3 |
| YPR124W | 1 | 3 |
| YIL005W | 1 | 3 |
| YEL042W | 1 | 3 |
| YJR116W | 1 | 3 |
| YKL140W | 2 | 3 |
| YCR026C | 2 | 3 |
| YBR086C | 2 | 3 |
| YOR316C | 2 | 3 |
| YML072C | 2 | 3 |
| YHR050W | 2 | 3 |
| YOR288C | 2 | 3 |
| YDR270W | 3 | 3 |
| YOR344C | 3 | 3 |
| YDR319C | 3 | 3 |
| YKR044W | 3 | 3 |
| YIL023C | 3 | 3 |
| YGR055W | 1 | 4 |
| YBR068C | 1 | 4 |
| YBR229C | 1 | 4 |
| YLR207W | 1 | 4 |
| YMR296C | 1 | 4 |
| YBR219C | 1 | 4 |

| | | |
|---|---|---|
| YIL030C | 2 | 4 |
| YBR029C | 2 | 4 |
| YOL003C | 2 | 4 |
| YDR456W | 2 | 4 |
| YML019W | 2 | 4 |
| YML013W | 3 | 4 |
| YOR154W | 3 | 4 |
| YOR175C | 1 | 5 |
| YBR296C | 1 | 5 |
| YJL019W | 3 | 5 |
| YGR036C | 3 | 5 |
| YGR171C | 3 | 5 |
| YDR038C | 3 | 5 |
| YPR079W | 3 | 5 |
| YPL176C | 3 | 5 |

**HEK293T Cells (from ref 25):**

| ENSEMBL ID | OMM Tier | ERM Tier |
|---|---|---|
| ENSG00000171714 | 3 | 2 |
| ENSG00000172292 | 3 | 2 |
| ENSG00000197296 | 3 | 2 |
| ENSG00000102158 | 3 | 2 |
| ENSG00000196950 | 3 | 2 |
| ENSG00000136868 | 4 | 2 |
| ENSG00000065308 | 4 | 2 |
| ENSG00000180776 | 4 | 2 |
| ENSG00000139641 | 5 | 2 |
| ENSG00000113194 | 5 | 2 |
| ENSG00000152078 | 5 | 2 |
| ENSG00000071537 | 3 | 3 |
| ENSG00000125827 | 3 | 3 |
| ENSG00000117868 | 4 | 3 |
| ENSG00000065923 | 4 | 3 |
| ENSG00000094975 | 4 | 3 |
| ENSG00000072274 | 4 | 3 |
| ENSG00000166479 | 4 | 3 |
| ENSG00000165240 | 5 | 3 |
| ENSG00000100528 | 5 | 3 |
| ENSG00000178607 | 5 | 3 |
| ENSG00000170385 | 5 | 3 |
| ENSG00000139514 | 5 | 3 |
| ENSG00000092068 | 5 | 3 |
| ENSG00000167130 | 3 | 4 |
| ENSG00000089597 | 3 | 4 |
| ENSG00000197081 | 3 | 4 |

| | | |
|---|---|---|
| ENSG00000197594 | 4 | 4 |
| ENSG00000104635 | 4 | 4 |
| ENSG00000141424 | 4 | 4 |
| ENSG00000197818 | 4 | 4 |
| ENSG00000143797 | 5 | 4 |
| ENSG00000100926 | 5 | 4 |
| ENSG00000011376 | 1 | 5 |
| ENSG00000182372 | 4 | 5 |
| ENSG00000107798 | 4 | 5 |
| ENSG00000114988 | 4 | 5 |
| ENSG00000198689 | 4 | 5 |
| ENSG00000101337 | 4 | 5 |
| ENSG00000001561 | 5 | 5 |
| ENSG00000144136 | 5 | 5 |
| ENSG00000164828 | 5 | 5 |

**Supplementary Table 11. Dual-, and ER- or Mito-PUP-tagged complex components.**

**Class A: Individual components dual-tagged (31 Complexes)**

acyl-CoA ceramide synthase complex

| Both | ER-PUP |
|---|---|
| Lac1p | Lag1p |

alpha-1,6-mannosyltransferase complex

| Both | ER-PUP |
|---|---|
| Anp1p | Mnn11p |
| Van1p | Hoc1p |
| | Mnn10p |
| | Mnn9p |

DASH complex

| Both | ER-PUP |
|---|---|
| Dad4p | Dad1p |
| | Dad3p |

DNF1-LEM3 P4-ATPase complex

| Both | ER-PUP |
|---|---|
| Dnf1p | Lem3p |

FET3-FTR1 high affinity iron permease complex

| Both | ER-PUP |
|---|---|
| Ftr1p | Fet3p |

Glucosidase II complex

| Both | ER-PUP |
|---|---|

| | |
|---|---|
| Rot2p | Gtb1p |

Glycosylphosphatidylinositol-N-acetylglucosaminyltransferase complex

| Both | ER-PUP |
|---|---|
| Gpi1p | Gpi15p |
| Eri1p | |
| Gpi2p | |

GPI-anchor transamidase complex

| Both | ER-PUP |
|---|---|
| Gpi16p | Gaa1p |
| | Gab1p |
| | Gpi17p |
| | Gpi8p |

HRD1 ubiquitin ligase complex

| Both | ER-PUP |
|---|---|
| Hrd3p | Hrd1p |
| Ubx2p | Der1p |
| Usa1p | Yos9p |

Luminal surveillance complex

| Both | ER-PUP |
|---|---|
| Hrd3p | Kar2p |
| | Yos9p |

Mannosyl phosphorylinositol ceramide synthase CSH1-CSG2

| Both | ER-PUP |
|---|---|
| Csg2p | Csh1p |

Oligosaccharyl transferase complex variant 1

| Both | ER-PUP |
|---|---|
| Ost6p | Wbp1p |
| Ost4p | Stt3p |
| | Ost1p |
| | Swp1p |

Oligosaccharyl transferase complex variant 2

| Both | ER-PUP |
|---|---|
| Ost4p | Wbp1p |
| | Stt3p |
| | Ost1p |
| | Swp1p |
| | Ost3p |

OPY2-MSB2 osmosensory complex

| Both | ER-PUP |
|---|---|
| Msb2p | Opy2p |

SPOTS complex

| Both | ER-PUP |
|---|---|
| Lcb1p | Lcb2p |
| | Orm1p |
| | Orm2p |
| | Sac1p |

SWI/SNF chromatin remodelling complex

| Both | ER-PUP |
|---|---|
| Snf11p | Snf6p |

60S cytosolic large ribosomal subunit

| Both | Mito-PUP |
|---|---|
| Rpl30p | Rpp0p |
| | Rpl39p |
| | Rpl10p |

1,3-beta-D-glucan synthase complex

| Both | ER-PUP | Mito-PUP |
|---|---|---|
| Gsc2p | Fks1p | Rho1p |

BUR1-BUR2 kinase complex

| Both |
|---|
| Bur2p |

Doa10 ubiquitin ligase complex

| Both |
|---|
| Ssm4p |
| Ubx2p |

Exomer complex

| Both |
|---|
| Bch2p |

FET5-FTH1 high affinity iron exporter complex

| Both |
|---|
| Fth1p |
| Fet5p |

Glycosylphosphatidylinositol-mannosyltransferase II complex

| Both |
|---|

Gpi18p

Ire1 complex

Both

Ire1p

Mannosyl phosphorylinositol ceramide synthase SUR1-CSG2

Both

Csg2p

Sur1p

MICOS complex

Both

Mic12p

NuA3 histone acetyltransferase complex

Both

Eaf6p

NuA4 histone acetyltransferase complex

Both

Eaf6p

PCL5-PHO85 kinase complex

Both

Pcl5p

Seipin complex

Both

Sei1p

UBP3-BRE5 ubiquitin hydrolase complex

Both

Bre5p

**Class B: Components tagged at either ER or Mitochondria (13 complexes).**

37S mitochondrial small ribosomal subunit

| ER-PUP | Mito-PUP |
| --- | --- |
| Sws2p | Mrp17p |
| | Rsm26p |
| | Mrps5p |
| | Rsm23p |
| | Mrps18p |

40S cytosolic small ribosomal subunit

| ER-PUP | Mito-PUP |
| --- | --- |
| Rps12p | Rps20p |

ESCRT-II complex

| ER-PUP | Mito-PUP |
| --- | --- |
| Snf8p | Vps36p |

Golgi transport complex

| ER-PUP | Mito-PUP |
| --- | --- |
| Cog8p | Cog6p |

MEX67-MTR2 nuclear RNA export factor complex

| ER-PUP | Mito-PUP |
| --- | --- |
| Mtr2p | Mex67p |

Nuclear pore complex

| ER-PUP | Mito-PUP |
| --- | --- |

| | |
|---|---|
| Pom34p | Nup60p |
| Ndc1p | Gle2p |
| Pom152p | |
| Nup2p | |

PAS complex

| ER-PUP | Mito-PUP |
|---|---|
| Vac7p | Fab1p |

PRP19-associated complex

| ER-PUP | Mito-PUP |
|---|---|
| Cdc40p | Cwc22p |
| Prp46p | |
| Prp9p | |
| Cwc25p | |

RNA polymerase I upstream activating factor complex

| ER-PUP | Mito-PUP |
|---|---|
| Rrn10p | Hhf2p |
| Uaf30p | |
| Rrn5p | |

SEC62-SEC63 complex

| ER-PUP | Mito-PUP |
|---|---|
| Sec66p | Sec63p |
| Sec62p | |

SMC5-SMC6 SUMO ligase complex

| ER-PUP | Mito-PUP |
|---|---|
| Nse5p | Kre29p |

Translocon complex

| ER-PUP | Mito-PUP |
| --- | --- |
| Sec61p | Sec63p |
| Sec66p | |
| Sec62p | |

TRAPPII protein complex

| ER-PUP | Mito-PUP |
| --- | --- |
| Trs31p | Trs65p |

**Class C: Components ER- or Mito-tagged**

| | |
|---|---|
| RED | Component in more than one complex |
| YELLOW | Same complex |

**ER-tagged components:**

| Standard Name | Complex | ER-PUP Tier |
|---|---|---|
| Aga1p | a-agglutinin | 1 |
| Apc5p | Anaphase-Promoting Complex variant 1 | 4 |
| Apc5p | Anaphase-Promoting Complex variant 2 | 4 |
| Apc5p | Anaphase-Promoting Complex variant 3 | 4 |
| Apc5p | Anaphase-Promoting Complex variant 4 | 4 |
| Asi1p | Asi complex | 4 |
| Asi2p | Asi complex | 3 |
| Asi3p | Asi complex | 3 |
| Atg31p | ATG1 kinase complex | 4 |
| Atg31p | Atg17-Atg31-Atg29 complex | 4 |
| Cnl1p | BLOC-1 complex | 5 |
| Ssn8p | CKM complex | 4 |
| Cks1p | CLN1-CDC28 kinase complex | 2 |
| Cln1p | CLN1-CDC28 kinase complex | 3 |
| Cks1p | CLN2-CDC28 kinase complex | 2 |
| Cln2p | CLN2-CDC28 kinase complex | 3 |
| Cks1p | CLN3-CDC28 kinase complex | 2 |
| Ctf19p | COMA complex | 3 |
| Cox14p | COX1 pre-assembly complex | 1 |
| Cue1p | CUE1-UBC7 ubiquitin-conjugating enzyme complex | 2 |
| Ela1p | CUL3-HRT1/ELC1/ELA1 ubiquitin ligase complex | 4 |

| | | |
|---|---|---|
| Nus1p | Dehydrodolichyl diphosphate synthase complex variant RER2 | 2 |
| Nus1p | Dehydrodolichyl diphosphate synthase complex variant SRT1 | 2 |
| Lif1p | DNA ligase IV complex | 4 |
| Rev7p | DNA polymerase zeta complex | 4 |
| Lem3p | DNF2-LEM3 P4-ATPase complex | 4 |
| Dnf3p | DNF3-CRF1 P4-ATPase complex | 3 |
| Ynr048Wp | DNF3-CRF1 P4-ATPase complex | 4 |
| Sld3p | DPB11-SLD3-SLD2 DNA replication complex | 4 |
| Cdc50p | DRS2-CDC50 P4-ATPase complex | 1 |
| Drs2p | DRS2-CDC50 P4-ATPase complex | 3 |
| Ldb18p | Dynactin | 4 |
| Slm4p | EGO complex | 5 |
| Emp24p | EMP24 complex | 1 |
| Erp1p | EMP24 complex | 1 |
| Erp2p | EMP24 complex | 1 |
| Erv25p | EMP24 complex | 1 |
| Emc1p | Endoplasmic Reticulum Membrane Complex | 1 |
| Emc3p | Endoplasmic Reticulum Membrane Complex | 1 |
| Mmm1p | ERMES complex | 2 |
| Erv41p | ERV41-ERV46 retrograde receptor complex | 2 |
| Erv46p | ERV41-ERV46 retrograde receptor complex | 2 |
| Far11p | FAR complex | 3 |
| Rad3p | General transcription factor complex TFIIH | 4 |
| Get1p | GET complex | 1 |
| Rad7p | Global genome repair CUL3/RAD7/RAD16/ELC1 ubiquitin ligase complex | 5 |
| Dug3p | Glutathione hydrolase complex | 3 |
| Gpi14p | Glycosylphosphatidylinositol-mannosyltransferase I complex | 1 |
| Hda1p | HDA1 complex | 3 |
| Hda2p | HDA1 complex | 3 |
| Sho1p | HICS complex | 2 |
| Hpa2p | Hpa2 acetyltransferase | 2 |

| | | |
|---|---|---|
| Mnl1p | HTM1-PDI1 exomannosidase complex | 1 |
| Pdi1p | HTM1-PDI1 exomannosidase complex | 1 |
| Ies3p | INO80 chromatin remodeling complex | 4 |
| Aur1p | Inositol phosphorylceramide synthase complex | 1 |
| Kei1p | Inositol phosphorylceramide synthase complex | 2 |
| Cyt1p | Mitochondrial electron transport complex III | 5 |
| Mot1p | MOT1-TBP transcription regulation complex | 2 |
| Mst28p | MST27-MST28 vesicle formation complex | 4 |
| Ncb2p | Negative cofactor 2 complex | 3 |
| Spo7p | Nem1-Spo7 phosphatase complex | 4 |
| Neo1p | NEO1-MON2-ARL1-DOP1 membrane remodeling complex | 2 |
| Pop8p | Nucleolar ribonuclease MRP complex | 4 |
| Pop8p | Nucleolar ribonuclease P complex | 4 |
| Rad4p | Nucleotide excision repair factor 2 complex | 4 |
| Rad7p | Nucleotide excision repair factor 4 complex | 5 |
| Erf2p | Palmitoyltransferase ERF2/SHR5 complex | 3 |
| Pcl10p | PCL10-PHO85 kinase complex | 4 |
| Pcl7p | PCL7-PHO85 kinase complex | 3 |
| Pex13p | Peroxisomal docking Pex13/Pex14/Pex 15 complex | 2 |
| Pex17p | Peroxisomal docking Pex13/Pex14/Pex 15 complex | 3 |
| Pex2p | Pex2/Pex10/Pex12 ubiquitin ligase complex | 2 |
| Pex21p | Pex7-Pex21 receptor complex | 3 |
| Pdr16p | Phosphatidylinositol transporter complex | 4 |
| Pmt1p | PMT1-PMT2 dolichyl-phosphate-mannose-protein mannosyltransferase complex | 1 |
| Pmt2p | PMT1-PMT2 dolichyl-phosphate-mannose-protein mannosyltransferase complex | 1 |
| Pmt1p | PMT1-PMT3 dolichyl-phosphate-mannose-protein mannosyltransferase complex | 1 |
| Pmt3p | PMT1-PMT3 dolichyl-phosphate-mannose-protein mannosyltransferase complex | 2 |
| Pmt4p | PMT4 dolichyl-phosphate-mannose-protein mannosyltransferase complex | 1 |
| Pmt2p | PMT5-PMT2 dolichyl-phosphate-mannose-protein mannosyltransferase complex | 1 |
| Pmt5p | PMT5-PMT2 dolichyl-phosphate-mannose-protein mannosyltransferase complex | 1 |
| Pmt3p | PMT5-PMT3 dolichyl-phosphate-mannose-protein mannosyltransferase complex | 2 |

| | | |
|---|---|---|
| Pmt5p | PMT5-PMT3 dolichyl-phosphate-mannose-protein mannosyltransferase complex | 1 |
| Msb4p | Polarisome | 2 |
| Rad55p | RAD55-RAD57 complex | 4 |
| Sas5p | SAS acetyltransferase complex | 5 |
| Scc4p | SCC2-SCC4 cohesin loader complex | 4 |
| Sec61p | SEC61 translocon complex | 1 |
| Prp9p | SF3A complex | 4 |
| Hsh49p | SF3B complex | 2 |
| Shu1p | Shu complex | 3 |
| Sec11p | Signal peptidase complex | 1 |
| Spc3p | Signal peptidase complex | 1 |
| Srp102p | Signal recognition particle receptor | 1 |
| Srp102p | Signal recognition particle receptor complex | 1 |
| Ssh1p | SSH1 translocon complex | 1 |
| Red1p | Synaptonemal complex | 5 |
| Tel2p | TEL2-TTI1-TTI2 complex | 5 |
| Est3p | Telomerase holoenzyme complex | 3 |
| Trs31p | TRAPPI protein complex | 4 |
| Trs31p | TRAPPIII protein complex | 4 |
| Sen2p | tRNA-intron endonuclease complex | 5 |
| Dsc2p | TUL1 E3 ubiquitin ligase complex | 2 |
| Tul1p | TUL1 E3 ubiquitin ligase complex | 2 |
| Ubx3p | TUL1 E3 ubiquitin ligase complex | 3 |
| Prp42p | U1 snRNP | 4 |
| Hsh49p | U2 snRNP | 2 |
| Prp9p | U2 snRNP | 4 |
| Alg14p | UDP-N-acetylglucosamine transferase complex | 4 |
| Utp5p | UTP-A complex | 1 |
| Stv1p | Vacuolar proton translocating ATPase complex, Golgi variant | 2 |
| Vma16p | Vacuolar proton translocating ATPase complex, Golgi variant | 1 |
| Vma3p | Vacuolar proton translocating ATPase complex, Golgi variant | 1 |

| | | |
|---|---|---|
| Vma16p | Vacuolar proton translocating ATPase complex, vacuole variant | 1 |
| Vma3p | Vacuolar proton translocating ATPase complex, vacuole variant | 1 |
| Vph1p | Vacuolar proton translocating ATPase complex, vacuole variant | 2 |
| Vps55p | VPS55-VPS68 sorting complex | 3 |
| Vps68p | VPS55-VPS68 sorting complex | 4 |
| Whi2p | WHI2-PSR1 phosphatase complex | 3 |
| Whi2p | WHI2-PSR2 phosphatase complex | 3 |

| | |
|---|---|
| RED | Same mRNA in more than one complex |
| YELLOW | Multiple mRNAs tagged for same complex |

**Mitochondria-tagged components:**

| Standard Name | Complex | Mito-PUP Tier |
|---|---|---|
| Mrp20p | 54S mitochondrial large ribosomal subunit | 5 |
| Mrpl11p | 54S mitochondrial large ribosomal subunit | 5 |
| Mrpl22p | 54S mitochondrial large ribosomal subunit | 3 |
| Mrpl24p | 54S mitochondrial large ribosomal subunit | 3 |
| Mrpl25p | 54S mitochondrial large ribosomal subunit | 3 |
| Mrpl32p | 54S mitochondrial large ribosomal subunit | 2 |
| Mrpl39p | 54S mitochondrial large ribosomal subunit | 2 |
| Mrpl3p | 54S mitochondrial large ribosomal subunit | 3 |
| Mrpl6p | 54S mitochondrial large ribosomal subunit | 4 |
| Rtc6p | 54S mitochondrial large ribosomal subunit | 5 |
| Mrs1p | bI3 intron splicing factor complex | 3 |
| Rad6p | BRE1-RAD6 ubiquitin ligase complex | 3 |
| Tpk1p | cAMP-dependent protein kinase complex variant 1 | 4 |
| Tpk1p | cAMP-dependent protein kinase complex variant 4 | 4 |
| Tpk1p | cAMP-dependent protein kinase complex variant 5 | 4 |
| Cbp3p | CBP3-CBP6 complex | 4 |
| Cbp6p | CBP3-CBP6 complex | 3 |
| Hap3p | CCAAT-binding factor complex | 4 |
| Bem1p | CLA4-BEM1-CDC24 polarity complex | 3 |
| Cat5p | CoQ biosynthetic complex | 3 |
| Cka2p | CURI complex variant 1 | 4 |
| Rrp7p | CURI complex variant 1 | 4 |
| Rrp7p | CURI complex variant 2 | 4 |

| | | |
|---|---|---|
| Cka2p | CURI complex variant 3 | 4 |
| Rrp7p | CURI complex variant 3 | 4 |
| Pri1p | DNA polymerase alpha:primase complex | 3 |
| Dpb4p | DNA polymerase epsilon complex | 1 |
| Hse1p | ESCRT-0 complex | 4 |
| Sec6p | Exocyst | 5 |
| Cap1p | F-actin capping protein complex | 1 |
| Mgm1p | FZO1-MGM1-UGO1 complex | 3 |
| Vps54p | GARP complex | 3 |
| Tfa1p | General transcription factor complex TFIIE | 4 |
| Vid24p | GID ubiquitin ligase complex | 4 |
| Gcv1p | Glycine decarboxylase multienzyme complex | 3 |
| Ssa1p | HAP1 transcriptional repressor complex - variant 1 | 2 |
| Hsc82p | HMC complex | 2 |
| Ssa1p | HMC complex | 2 |
| Dpb4p | ISW2 chromatin remodeling complex | 1 |
| Pbi2p | LMA1 complex, TRX1 variant | 3 |
| Pbi2p | LMA1 complex, TRX2 variant | 3 |
| Mcm5p | MCM complex | 3 |
| Mcm7p | MCM complex | 4 |
| Iqg1p | MIH complex | 2 |
| Imp2'p | Mitochondrial inner membrane peptidase complex | 4 |
| Tim44p | Mitochondrial inner membrane pre-sequence translocase complex | 2 |
| Tim50p | Mitochondrial inner membrane pre-sequence translocase complex | 4 |
| Idh1p | Mitochondrial isocitrate dehydrogenase complex (NAD+) | 3 |
| Idh2p | Mitochondrial isocitrate dehydrogenase complex (NAD+) | 3 |
| Tom40p | Mitochondrial outer membrane translocase core complex | 1 |
| Tom40p | Mitochondrial outer membrane translocase holocomplex | 1 |
| Tom70p | Mitochondrial outer membrane translocase holocomplex | 1 |
| Mas2p | Mitochondrial processing peptidase complex | 3 |
| Atp15p | Mitochondrial proton-transporting ATP synthase complex | 2 |

| | | |
|---|---|---|
| Atp2p | Mitochondrial proton-transporting ATP synthase complex | 2 |
| Atp4p | Mitochondrial proton-transporting ATP synthase complex | 1 |
| Sam50p | Mitochondrial sorting and assembly machinery complex | 3 |
| Rad50p | MRE11-RAD50-XRS2 meiotic recombination initiation complex | 5 |
| Rad6p | MUB1-RAD6-UBR2 ubiquitin ligase complex | 3 |
| Vac17p | MYO2-VAC17-VAC8 transport complex | 3 |
| Btt1p | Nascent polypeptide-associated complex, BTT1-EGD2 variant | 4 |
| Nfs1p | NFS1-ISD11 cysteine desulphurase complex | 3 |
| Hhf2p | Nucleosome, variant HTA1-HTB1 | 5 |
| Hta1p | Nucleosome, variant HTA1-HTB1 | 2 |
| Htb1p | Nucleosome, variant HTA1-HTB1 | 2 |
| Hhf2p | Nucleosome, variant HTA1-HTB2 | 5 |
| Hta1p | Nucleosome, variant HTA1-HTB2 | 2 |
| Hhf2p | Nucleosome, variant HTA2-HTB1 | 5 |
| Htb1p | Nucleosome, variant HTA2-HTB1 | 2 |
| Hhf2p | Nucleosome, variant HTA2-HTB2 | 5 |
| Hhf2p | Nucleosome, variant HTZ1-HTB1 | 5 |
| Htb1p | Nucleosome, variant HTZ1-HTB1 | 2 |
| Hhf2p | Nucleosome, variant HTZ1-HTB2 | 5 |
| Ctr9p | PAF1 complex | 5 |
| Pcl1p | PCL1-PHO85 kinase complex | 2 |
| Vps15p | Phosphatidylinositol 3-kinase complex I | 5 |
| Vps15p | Phosphatidylinositol 3-kinase complex II | 5 |
| Rad6p | RAD6-RAD18 ubiquitin ligase complex | 3 |
| Sth1p | RSC complex variant RSC1 | 2 |
| Sth1p | RSC complex variant RSC2 | 2 |
| Rsp5p | RSP5-BUL1 ubiquitin ligase complex | 5 |
| Bul2p | RSP5-BUL2 ubiquitin ligase complex | 3 |
| Rsp5p | RSP5-BUL2 ubiquitin ligase complex | 5 |
| Cpr1p | SET3C histone deacetylase complex | 2 |
| Sit4p | SIT4-SAP155 phosphatase complex | 5 |

| | | |
|---|---|---|
| Sap185p | SIT4-SAP185 phosphatase complex | 3 |
| Sit4p | SIT4-SAP185 phosphatase complex | 5 |
| Sit4p | SIT4-SAP190 phosphatase complex | 5 |
| Dig1p | Ste12/Dig1/Dig2 transcription regulation complex | 3 |
| Bdf1p | Swr1 chromatin remodelling complex | 3 |
| Sit4p | TAP42-RRD1-SIT4 phosphatase complex | 5 |
| Dig1p | Tec1/Ste12/Dig1 transcription regulation complex | 3 |
| Toa2p | Transcription factor TFIIA complex | 3 |
| Snu66p | U4/U6.U5 tri-snRNP complex | 5 |
| Rad6p | UBR1-RAD6 ubiquitin ligase complex | 3 |
| Cka2p | UTP-C complex variant 1 | 4 |
| Rrp7p | UTP-C complex variant 1 | 4 |
| Rrp7p | UTP-C complex variant 2 | 4 |
| Cka2p | UTP-C complex variant 3 | 4 |
| Rrp7p | UTP-C complex variant 3 | 4 |
| Sec9p | Vesicular t-SNARE complex, SSO1 variant | 3 |
| Sec9p | Vesicular t-SNARE complex, SSO2 variant | 3 |

**Supplementary Table 12. Tagged non-mRNAs.**

**Dual-tagged:**

| Systematic Name | ER-PUP Tier | Mito-PU Tier |
|---|---|---|
| snR11 | 1 | 3 |
| tK(UUU)D | 2 | 4 |
| snR7-L | 3 | 4 |
| snR84 | 3 | 5 |
| snR37 | 4 | 5 |

**ER-tagged Unique:**

| SGD ID | Feature Type | ER-PUP Tier |
|---|---|---|
| SCR1 | ncRNA gene | 1 |
| tD(GUC)O | tRNA gene | 2 |
| snR35 | snoRNA gene | 2 |
| tQ(UUG)H | tRNA gene | 2 |
| snR47 | snoRNA gene | 2 |
| snR33 | snoRNA gene | 2 |
| snR31 | snoRNA gene | 2 |
| tK(UUU)L | tRNA gene | 2 |
| tH(GUG)E1 | tRNA gene | 3 |
| tV(AAC)K2 | tRNA gene | 3 |
| tH(GUG)H | tRNA gene | 3 |
| snR7-S | snRNA gene | 3 |
| snR128 | snoRNA gene | 3 |
| RPR1 | ncRNA gene | 3 |
| snR45 | snoRNA gene | 3 |
| RUF20 | ncRNA gene | 3 |
| snR6 | snRNA gene | 3 |
| NME1 | snoRNA gene | 3 |
| tT(AGU)B | tRNA gene | 3 |
| snR67 | snoRNA gene | 3 |
| tD(GUC)G1 | tRNA gene | 3 |
| tM(CAU)J2 | tRNA gene | 3 |
| tD(GUC)M | tRNA gene | 3 |
| tK(UUU)O | tRNA gene | 3 |
| snR32 | snoRNA gene | 3 |
| snR30 | snoRNA gene | 3 |

| | | |
|---|---|---|
| snR48 | snoRNA gene | 3 |
| SRG1 | ncRNA gene | 3 |
| snR4 | snoRNA gene | 3 |
| snR63 | snoRNA gene | 3 |
| tR(UCU)G3 | tRNA gene | 4 |
| snR66 | snoRNA gene | 4 |
| tR(UCU)G1 | tRNA gene | 4 |
| tG(GCC)G1 | tRNA gene | 4 |
| snR55 | snoRNA gene | 4 |
| snR3 | snoRNA gene | 4 |
| snR64 | snoRNA gene | 4 |
| tK(UUU)G1 | tRNA gene | 4 |
| snR42 | snoRNA gene | 4 |
| tS(AGA)H | tRNA gene | 4 |
| tY(GUA)F1 | tRNA gene | 4 |
| tN(GUU)O2 | tRNA gene | 4 |
| tX(XXX)L | tRNA gene | 4 |
| tG(GCC)O2 | tRNA gene | 4 |
| snR69 | snoRNA gene | 4 |

**Mitochondria-tagged Unique:**

| Systematic Name | Mito-PU Tier |
|---|---|
| snR190 | 3 |
| tH(GUG)G2 | 4 |
| tN(GUU)K | 5 |
